# STAT1-Mediated Regulation of IL-17A/CEBPB/NF-κB Axis in HIV-1 Infected Human Cerebral Organoids Reveals Therapeutic Targets for Neuroprotection

**DOI:** 10.64898/2025.12.30.697044

**Authors:** Karthick Chennakesavan, Malaroviyam Samikkannu, Shreyan Gupta, Olasunkanmi Israel Tayo, Arpan Acharya, James J. Cai, Siddappa N. Byrareddy, Thangavel Samikkannu

## Abstract

HIV-associated neurological complications remain a major concern in people living with HIV (PLWH), even under effective viral suppression with combination antiretroviral therapy (cART), underscoring unresolved mechanisms driving HIV-related neurodegeneration. To address this gap, we used HIV-1–infected human cerebral organoids (hCOs) containing microglia as a physiologically relevant three-dimensional model of the central nervous system (CNS) and validated key findings in simian immunodeficiency virus (SIV)-infected, cART-treated rhesus macaque model. Integrating single-nucleus RNA sequencing with ATAC sequencing, we identified cell-type-specific alterations in dual innate immune–driven inflammatory signaling pathways across neural and glial populations. Microglia were preferentially infected and activated, initiating IL-17A–mediated cascades involving IFN-γ/STAT1, CEBPB, and NF-κB pathways, which converged with cGAS–STING and Notch–NEURL1 signaling, both associated with neuroinflammation and synaptic dysfunction. cART suppressed viral replication, reduced IL-17A–driven neuroinflammation, improved mitochondrial function, and restored synaptic gene expression. Collectively, these findings identify IL-17A, STAT1, CEBPB, and cGAS–STING as key molecular drivers of HIV-associated neuroinflammation and establish a novel mechanistic link between IL-17A signaling and microglial remodeling, providing a translational framework for the development of CNS-targeted therapies in PLWH.

## INTRODUCTION

According to the 2024 Joint United Nations Programme on HIV/AIDS (UNAIDS) report, an estimated 40.8 million people worldwide are living with HIV (PLWH), with 1–2 million new infections occurring annually [1]. Despite the widespread availability of highly effective combination antiretroviral therapy (cART), only ∼50% of PLWH achieve undetectable plasma HIV viral loads, mainly due to incomplete adherence, pharmacokinetic variability, drug resistance, or immune dysfunction. HIV-associated neurocognitive disorder (HAND) remains a major complication, affecting 30–50% of PLWH, most often in its milder forms, even in the long-acting cART era. Persistent low-level HIV replication in the brain, despite systemic viral suppression, is a recognized driver of chronic neuroinflammation [2].

Single-cell and single-nucleus RNA sequencing (scRNA-seq and snRNA-seq) have significantly advanced our ability to resolve cell-type–specific gene expression and molecular pathways, particularly when applied to human cerebral organoids (hCOs) [3, 4]. Integrating transcriptomic, immune response, and inflammatory profiles from single-cell datasets enables detailed dissection of cellular complexity in the human brain and the mechanisms driving neurodegenerative disease progression [4, 5]. hCOs recapitulate key features of the human central nervous system (CNS), encompassing diverse resident cell types such as neurons, astrocytes, oligodendrocytes, and microglia, while preserving spatial organization and higher-order functional architecture [5, 6]. These characteristics make hCOs a powerful platform for modeling neurodevelopment, neurodegeneration, and intercellular interactions in both physiological and pathological contexts.

To clarify HIV neuropathogenesis, it is essential to identify CNS-resident cells that mediate immune responses. In the brain parenchyma, immune defense is primarily carried out by glial cells, which are long-lived myeloid cells that reside permanently within brain tissue [7].

Microglia, the brain’s resident macrophages, are highly susceptible to HIV infection and contribute to the establishment of long-lived viral reservoirs. Once infected, microglia support persistent viral replication and initiate a chronic inflammatory milieu, releasing pro-inflammatory cytokines and inducing type I interferon-stimulated genes (ISGs) [8]. These ISGs perpetuate inflammation, impair synaptic plasticity, and contribute to HAND and other neurodegenerative disorders [9].

One key regulator of neuroinflammation is CCAAT/enhancer-binding protein-β (CEBPB), also known as nuclear factor interleukin-6 (NF-IL6), a transcription factor central to inflammatory signaling in the CNS [10, 11]. CEBPB coordinates glial inflammatory responses and may interact with interleukin-17A (IL-17A), a pro-inflammatory cytokine produced primarily by Th17 cells, to exacerbate neuroinflammation. IL-17A synergizes with tumor necrosis factor (TNF) to increase production of chemokines such as CCL20, which recruit additional Th17 cells and promote HIV replication [10, 12, 13]. Notably, previous studies have shown that CEBPB regulates the expression of inflammation-related factors, including monocyte chemotactic protein-1 (MCP-1), in HIV-treated astrocytes and microglia [11, 14], and plays a dual role, propagating inflammation while also contributing to resolution during tissue repair [13, 15]. Specified persistent immune activation in HIV-associated neuroinflammation despite cART, the CEBPB/IL-17A axis represents a promising but underexplored therapeutic target.

Another candidate of interest is neuralized-like protein 1A (NEURL1), an E3 ubiquitin-protein ligase involved in Notch signaling and neuronal differentiation. In glial cells, NEURL1 maintains homeostasis and alters cellular CNS inflammatory responses [16]. Dysregulation of NEURL1, potentially via impaired Notch–JAG1 signaling, has been linked to glial vulnerability under pathological conditions such as HIV infection and tumor progression [17]. The Notch pathway also influences IL-17A production [18], suggesting that the NEURL1–Notch–IL-17A axis could represent a novel mechanism linking HIV-infection to CNS inflammation and synaptic dysfunction. Despite its potential importance, NEURL1 remains poorly studied in the context of HIV neuropathogenesis.

In addition to viral and host factors, therapeutic regimens themselves may influence CNS inflammation. For example, the cART regimen ATRIPLA (Efavirenz, Emtricitabine, Tenofovir) can suppress NF-κB activation, potentially reducing inflammatory signaling and preventing neuronal injury. However, dissecting these interactions in vivo remains challenging due to the complexity of CNS immune networks. Human cerebral organoids (hCOs), combined with advanced single nuclei multiomic approaches, provide a unique platform to interrogate these pathways in a controlled yet physiologically relevant context. In this study, we investigate how cART-suppressed HIV-1 infection alters the expression and functional activity of NEURL1 and the CEBPB/IL-17A axis in hCOs, and we elucidate the downstream effects on neuroinflammation, mitochondrial function, and neuronal integrity. We further integrate data from simian immunodeficiency virus (SIV)-infected, cART-treated rhesus macaques to validate key findings and assess cell-type-specific transcriptomic changes in the inflammatory profile using snRNA-seq and snATAC-seq. This combined approach aims to identify and validate novel immunotherapeutic targets to mitigate HIV-1-associated neuroinflammation and preserve cognitive function in people living with HIV.

## MATERIALS AND METHODS

### Cell Culture

The human induced pluripotent stem cell (iPSC) line (Episomal, PBMC-derived; Alstem, Cat. #iPS15), hereafter referred to as iPS15, was maintained in feeder-free conditions using mTeSR1 Plus medium (STEMCELL Technologies, Cat. #85850) supplemented with ROCK inhibitor (Y-27632; STEMCELL Technologies, Cat. #72307). Cells were cultured on Matrigel-coated 6-well plates, with medium changes performed every other day. Cultures were incubated at 37 °C in a humidified atmosphere containing 5% CO₂.

### Cerebral Organoid Development

Human cerebral organoids were generated using the human iPSC line (iPS15) and the STEMdiff™ Cerebral Organoid Kit (STEMCELL Technologies, Cat. #08570), following the manufacturer’s instructions. Freshly prepared media were used for each developmental stage: embryoid body (EB) formation (Day 0–5), neural induction (Day 5–7), expansion (Day 7–10), and maturation (Day 10–90).

On Day 0, iPSCs were dissociated using Gentle Cell Dissociation Reagent (STEMCELL Technologies, Cat. #100-0485) and resuspended in mTeSR™ medium (STEMCELL Technologies, Cat. #100-0276). Cells were centrifuged at 300 × g for 5 minutes, and the pellet was resuspended in EB seeding medium containing 10 µM Y-27632. Approximately 9,000 cells per well were seeded into a low-attachment 96-well plate for EB formation and incubated at 37 °C with 5% CO₂. On Days 2 and 4, 100 µL of EB formation medium was added to each well.

On Day 5, 1–2 EBs per well were transferred into a low-attachment 24-well plate containing induction medium and incubated at 37 °C with 5% CO₂.

On Day 7, EBs were embedded in 15 µL of Matrigel® (Corning, Cat. #354-277) per EB on an organoid embedding sheet. A total of 4–6 embedded EBs were placed in each well of an ultra-low-attachment 6-well plate containing 3 mL of expansion medium and incubated at 37 °C with 5% CO₂.

On Day 10, the expansion medium was replaced with maturation medium, and organoids were transferred to an orbital shaker and maintained at 37 °C with 5% CO₂. Half-media changes were performed every 3–4 days until the experiments were done, as illustrated in Figure 1a.

**Figure 1.**
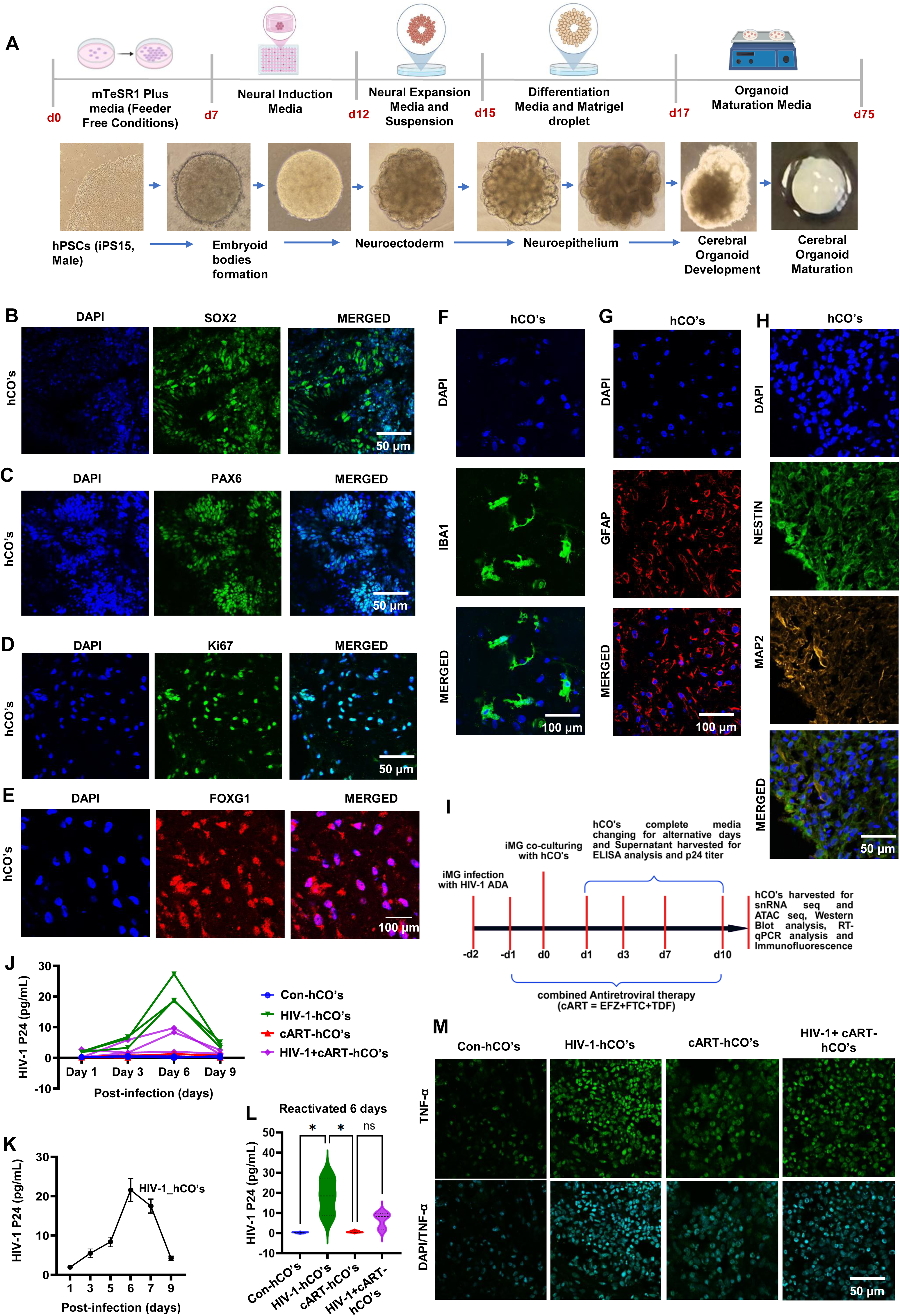
Generation and characterization of hCOs from iPS15 Line. **(A)** Diagrammatic representation illustrating the developmental stages of hCOs from iPS15 Lines. **(B-E)** Immunostaining analysis of hCOs at 60 days to assess the expression of stem cell proliferation markers, SOX2, PAX6, Ki67, and FOXG1, respectively. **(F-H)** Characterization of glial and neuronal cell populations within 60-day-old hCOs using specific markers: **(F)** Iba1, a microglial marker; **(G)** GFAP, an astrocytic marker highlighting glial cell maturation; **(H)** NESTIN, and MAP2, a neuronal marker representing mature neurons and dendritic structures. Scale bars, 50 and 100 μm. **(I)** schematic representation illustrating the experimental setup in which hCOs are infected with the HIV-1-ADA virus strain. **(J-L)** HIV p24 ELISA analysis was performed on supernatant samples collected from different experimental groups, including control, HIV-1 infected, and with or without cART treatment. A total of 6 hCOs per condition were analyzed across 3 independent experiments. **(M)** Immunofluorescence of hCOs stained with anti-TNF-α antibodies (green), a pro-inflammatory marker, and with DAPI, overexpression of TNF-α was observed in HIV-1 infected organoids compared to controls. Data are represented as mean ± SD (n = 3). One-way ANOVA followed by Tukey’s multiple comparison test. Statistically significant at *P < 0.01, and ns-Not significant.

### Derivation of microglia from iPSC cells

Hematopoietic progenitor cells (HPCs) were derived from the iPS15 line using the STEMdiff™ Hematopoietic Kit (STEMCELL Technologies, Cat. #05310) according to the manufacturer’s instructions. The HPCs were then seeded and differentiated into microglia using the STEMdiff™ Microglial Differentiation Kit (Cat. #100-0019), followed by further maturation with the STEMdiff™ Microglial Maturation Kit (Cat. #100-0020), as per the manufacturer’s protocol. Briefly, hiPSC-derived HSCs (2 × 10^5^ cells/well) were plated in Matrigel®-coated 6-well plates containing 2 ml of STEMdiff™ Microglia differentiation medium and maintained at 37 °C with 5% CO2. Cell cultures were refreshed every other day by supplementing with 1 ml of STEMdiff™ Microglia differentiation medium for 12 days. Following this initial period, the entire cell suspension was centrifuged at 300 g for 5 min and subsequently seeded into fresh Matrigel®-coated 6-well plates with 1 ml of STEMdiff™ Microglia differentiation medium, maintaining the same culture conditions for an additional 12 days with regular medium replenishment. On day 24, 1 × 10^6^ cells (at a density of 1 × 10^5^ cells/cm^2^) were harvested, centrifuged at 300 g for 5 min, and cultured in Matrigel®-coated plates with STEMdiff™ Microglia maturation medium for 4 days. On day 28, the hiPSC-derived iMGs were collected for subsequent HIV-1 infection.

### HIV-1 ADA infection and cART treatment of hCOs

The R5-tropic HIV-1 ADA strain was propagated in human PBMCs and titrated as described previously [19], and used for infecting hCOs. iPSC-derived microglia (5 × 10^4^ cells) were cultured in 6-well plates and infected with HIV-1 ADA at a multiplicity of infection (MOI) of 0.5 in microglial maturation medium supplemented with Polybrene (5 µg/mL) to enhance viral entry. After 12–16 hours of incubation, the cells were washed with PBS and maintained in fresh maturation medium for 24 hours. The HIV-1-infected microglia were then surface-seeded onto 60-day-old cerebral organoids and co-cultured for an additional 9 days to allow integration and interaction within the hCOs. The medium was partially replaced every 3 days.

In parallel, 60-day-old organoids were pre-treated for 2 hours with ATRIPLA, a combination antiretroviral therapy (cART) regimen containing 5 µM each of Efavirenz (EFV), Emtricitabine (FTC), and Tenofovir (TDF), with or without subsequent HIV-1 ADA infection, and maintained for 9 days. Culture supernatants were collected daily, and HIV p24 antigen levels were quantified using ELISA on days 1, 3, 6, and 9 post-infections.

### Single-nucleus dissociation and library preparation

For single-nucleus ATAC sequencing (snATAC-seq), nuclei were isolated using Nuclei Extraction Buffer (130-128-024, Miltenyi Biotec) supplemented with a protease inhibitor cocktail (5892791001; Roche). Human cerebral organoids were finely chopped into fragments smaller than 0.5 mm and homogenized in 2 mL of ice-cold lysis buffer using a Dounce homogenizer. The homogenate was incubated on ice for 5 minutes, followed by adding another 2 mL of lysis buffer. The lysate was passed through a 40-μm cell strainer and centrifuged at 500 × *g* for 5 minutes at 4 °C. The pellet was resuspended in fresh lysis buffer, washed with 4 mL, and incubated on ice for 5 minutes. After a second centrifugation step, the nuclei pellet was resuspended in Nuclei Buffer (10x Genomics, PN-2000153), filtered through a 5-μm cell strainer, and counted.

For single-nucleus RNA sequencing (snRNA-seq), RNase inhibitors (Promega, N2615; Life Technologies, AM2696) were added to the lysis buffer to preserve RNA integrity. The final nuclei pellet was resuspended in Nuclei Suspension Buffer (1× PBS, 1% bovine serum albumin, 0.1% RNase inhibitor) and stored in CryoStor CS10 freezing media. Libraries for both snATAC-seq and snRNA-seq were prepared using the 10x Genomics Chromium platform according to the manufacturer’s protocol.

### Single-Cell RNA and ATAC Sequencing

#### Cell Thawing and Nuclei Isolation

Cryopreserved cells were thawed following the CryoStor CS10 Thawing Protocol (StemCell Technologies, https://tinyurl.com/veyk9ab5) using pre-warmed RPMI + 10% FBS for the initial 1:10 dilution and 1X PBS + 0.04% for the washing and final resuspension steps. Thawed cells were stained with acridine orange and propidium iodide (Logos Biosystems PN-F23001) and assessed for viability, concentration, and singleness using the LUNA-FX7 Dual Fluorescence Cell Counter (Logos Biosystems). Nuclei were isolated following the 10x Genomics Low Cell Input Nuclei Isolation Protocol (PN-CG000365 Rev B, http://tinyurl.com/ycynrdhj). Approximately 40,000 pelleted cells per sample were resuspended in 1X Lysis Buffer, incubated on ice for 3 minutes, washed twice, and resuspended in a proprietary buffer. The nuclei were again assessed for viability, concentration, and singleness using the LUNA-FX7 [20].

#### 10x Genomics Multiome Assay for Dual Chromatin Accessibility and Gene Expression Profiling

Multiome library preparation was completed using the 10x Genomics Chromium X Controller and the Chromium Next GEM Single Cell Multiome ATAC + Gene Expression Reagent Bundle (10x Genomics PN-1000283) following the manufacturer’s user guide (PN-CG000338 Rev F, http://tinyurl.com/bdhe7rs4). Freshly isolated nuclei suspensions were incubated in a transposition mix to simultaneously fragment open regions of chromatin and ligate adapter sequences to the ends of the fragmented DNA. Approximately 16,500 nuclei were transposed per sample, with an ultimate recovery target of 10,000 nuclei per sample. Transposed nuclei, a master mix of reverse transcription and template-switching reagents, and gel beads coated with barcoding oligos were then microfluidically mixed into an oil medium to form gel beads in emulsion (GEMs) using the Chromium Controller instrument (10x Genomics). Individual partitioned single-nucleus reaction mixtures were formed by lysing the cells and dissolving the gel beads within the GEMs. The GEMs were incubated to generate transposed DNA fragments and full-length cDNA, both tagged with cell barcodes, unique molecular identifiers (UMIs), and 5’ and 3’ priming sites provided by the gel bead oligos. A quenching agent was added to stop the reaction, and GEMs were stored at -80°C for up to 4 weeks. The GEMs were thawed, broken, and purified using magnetic DNA-binding beads (Dynabeads MyOne SILANE beads, Invitrogen). The purified products were pre-amplified via PCR (7 cycles) with primer pairs specific to the DNA fragments (for ATAC) and cDNA (for Gene Expression). Pre-amplified products were then split into separate aliquots for ATAC and Gene Expression (GEX) library preparation.

#### ATAC Library Preparation

The Illumina P7 sequencing adapter and a uniquely identifying single index were added to pre-amplified, transposed DNA fragments via PCR (8 cycles). The completed ATAC sequencing libraries were then purified using SPRIselect beads, visualized and quantified using the Bioanalyzer High Sensitivity DNA, and pooled together before sequencing.

#### Gene Expression Library Preparation

The pre-amplified cDNA was further amplified via PCR (7 cycles) to generate enough mass for library preparation and subsequently purified with SPRIselect beads. The cDNA was subjected to fragmentation, end repair, A-tailing, double-sided size selection using SPRI select beads, and Illumina sequencing adaptor ligation. Uniquely identifiable dual indexes were added via PCR (16 cycles). The completed gene expression libraries were then purified using SPRI select beads, visualized and quantified using the Bioanalyzer High Sensitivity DNA chip, and pooled together for sequencing.

#### Sequencing and Data Analysis

ATAC and GEX pools were sequenced separately at the UNC High Throughput Sequencing Facility. Pools were denatured and diluted following standard Illumina protocol and spiked with 1% PhiX sequencing control (Illumina) before sequencing. ATAC libraries were sequenced on two NextSeq 2000 P4 flow cells to a total depth of 3.1 billion read pairs passing quality filters. GEX libraries were sequenced on a single NovaSeq 6000 S4 flow cell to a total depth of 8.0 billion read pairs passing quality filters. Libraries were sequenced in the appropriate format for ATAC (Read 1: 50 cycles, i7 index: 8 cycles, i5 Index: 24 cycles, Read 2: 50 cycles) or GEX (Read 1: 28 cycles, i7 index: 10 cycles, i5 Index: 10 cycles, Read 2: 90 cycles) accordingly. Sequencing data was aligned to 10x’s prebuilt human reference genome (GRCh38-2020-A, https://tinyurl.com/mwvy8e9m) and processed through Cell Ranger ARC v2.0.2 using default settings.

#### Single-cell multi-omics analysis

Multi-omics (ATAC + RNA) data were generated from three hCOs conditions: control (Con-hCOs), HIV-1 infected (HIV-1-hCOs), and HIV-1-infection with cART treatment (HIV-1 + cART-hCOs).

##### a. Preprocessing and Quality Control

Raw sequencing reads were processed with Cell Ranger ARC v2.0.2 (10x Genomics), aligned to GRCh38, and used to generate feature-barcode matrices. Downstream analyses were performed using Seurat v5 and Signac v1.14 in R. Seurat objects with separate RNA and ATAC assays were created per sample. EnsDb provided peak annotations for ATAC-seq*. Hsapiens.v86*. To ensure unique cell IDs, Seurat objects were merged with dataset-specific prefixes. Quality control filtering used the following thresholds: ATAC-seq—fragments >200 and <30,000; nucleosome signal <3; TSS enrichment >1. RNA-seq—UMI counts >1,000 and <30,000; genes expressed >500.

For cross-species validation of the hCO dataset, we reanalyzed publicly available snRNA-seq data of healthy control and SIV-infected, ART-treated macaque brains [21–23]. The snRNA-seq data of the control rhesus macaques (RMs) were obtained from NCBI GEO, accession # GSE233278, and the data of the SIV-infected, ART-treated RMs were obtained from NCBI GEO, accession # GSE209606, respectively. Data were imported into R using Zellkonverter [24], and converted into a Seurat object to enable downstream processing. Doublets were removed using the Doublet Finder package in R. Low-quality nuclei were removed based on three metrics: nuclei with gene count <200 or >10,000, UMI count <400 or >1000, and nuclei with mitochondria UMI proportion greater >1%. As a result, a total of 28,760 and 7217 high-quality nuclei were retained for downstream analysis in the SIV-infected ART-treated and control RMs, respectively. Downstream analysis of RMs data was carried out using the same pipeline used for analyzing the hCO dataset.

##### b. Normalization, Integration, and Dimensionality Reduction

For the hCO dataset, RNA-seq data were normalized using *NormalizeData*, and 2,000 highly variable genes were selected using *FindVariableFeatures*. PCA was applied after scaling. Harmony-based batch correction was performed on PCA embeddings using *IntegrateLayers* (method = “Harmony Integration”). For ATAC-seq data, peaks were filtered with *FindTopFeatures* (min.cutoff = 5), normalized using *RunTFIDF*, and reduced using SVD. A weighted nearest neighbor (WNN) graph was constructed via *FindMultiModalNeighbors* using Harmony-integrated RNA (top 30 PCs) and LSI-reduced ATAC (dimensions 2–40). UMAP embedding was computed using *RunUMAP*.

For the RMs data, normalization was performed using SCTransform normalization within Serat, and batch effect was corrected using Harmony, followed by PCA. The top 20 principal components (PCs) were selected for downstream analysis.

##### c. Cell Clustering and Annotation

Unsupervised clustering was performed for both datasets following dimensionality reduction. In the hCO dataset, clustering was performed using the Louvain algorithm via *FindClusters* (resolution = 0.8) on the WNN graph. For the RMs dataset, clustering was performed using the Louvain algorithm implemented via the *FindClusters* function in the Seurat package, with 0.3 - 0.5 resolution. These unsupervised clustering approaches allowed for the identification of transcriptionally distinct cell populations in the single-nucleus RNA-seq dataset.

For both datasets, cluster-specific marker genes were identified using *FindAllMarkers* (log2FC ≥ 1, min.pct = 0.1, FDR-adjusted *p* < 0.01). Manual annotation was performed using known cell-type markers from the literature and public databases. Broad categories included astrocytes (Astro), oligodendrocytes (Oligo), neural stem cells (NSCs), neurons, microglia, oligodendrocyte precursor cells (OPCs), and cycling radial glia cells (cRGCs).

##### d. Differential Expression, Accessibility, and Pathway Analysis

For the hCO dataset, differential expression (DE) and accessibility (DA) were analyzed per cell type. Comparisons were made between HIV-1-hCOs vs. Con-hCOs and HIV-1+cART-hCOs vs. HIV-1-hCOs using *FindMarkers* (Wilcoxon test; log2FC ≥ 1; min.pct = 0.05), *p*-values < 0.05 (Bonferroni) were considered significant. For ATAC-seq, differentially accessible regions (DARs) were identified using the same criteria. Nearest genes to DARs were assigned using *ClosestFeature*. Functional enrichment was performed using the fgsea R package with Gene Ontology (GO) biological processes from MSigDB v2024.1. Pathways with adjusted *p* < 0.05 and ≥2 genes were considered significant. Leading-edge genes were extracted, and network plots were generated per cell type to visualize pathway–gene relationships. KEGG pathway gene sets were also used to examine expression trends in pathways of interest. Enrichr analysis was performed using DEGs for microglia.

RMs data, differential gene expression analysis was performed for each cluster using Scanpy’s rank genes group’s function (Wilcoxon rank-sum test; adjusted p-value < 0.05 and log fold change > 0.25). To reduce cell-level variability and improve robustness, pseudobulk gene expression profiles were generated by aggregating cells at the sample level within each cell type. Functional enrichment analysis of biological processes, GO Biological Process pathways, was performed using the same fgsea pipeline as previously described (based on MSigDB v2024.1). Likewise, upregulated genes were mapped from gene symbols to ENTREZ gene IDs using the bitr function from the clusterProfiler R package, using the Bioconductor package with the org.Hs.eg.db database used as the annotation database. KEGG pathway enrichment was then performed using the enrich KEGG function, specifying Homo sapiens (organism = “hsa”) and a significance threshold of p<0.05, following the same criteria and visualization strategy as in the iPSC organoids.

### ELISA Assay

**p24 ELISA** – HIV viral load in supernatants from hCOs was quantified using the HIV p24 SimpleStep ELISA® Kit (Abcam, Cat# ab218268), following the manufacturer’s instructions. Supernatants were collected every three days up to 9 days post-HIV infection, including during cART treatment. Briefly, 50 µL of sample or standard was added to each well of the ELISA plate, followed by 50 µL of antibody cocktail. The plate was incubated at room temperature for 1 hour. Wells were then washed three times with wash buffer. Subsequently, 100 µL of TMB substrate was added to each well and incubated for 10 minutes in the dark on a shaker at 400 rpm. The reaction was stopped by adding 100 µL of stop solution and incubating for 1 minute on the shaker. Absorbance was measured at 450 nm using a microplate reader [25].

### Cytokine Quantification by Sandwich ELISA

Supernatants were collected from HIV-1–infected and cART-treated hCOs on day 6 post-infection for cytokine analysis. IL-6, IL-1β, and IL-4 levels were quantified using ELISA kits specific to each cytokine, according to the respective manufacturers’ protocols. Briefly, 50 µL of each sample or standard was added in duplicate to wells of a 96-well plate pre-coated with capture antibody. An equal volume (50 µL) of biotin-conjugated detection antibody was then added to each well, followed by a 2-hour incubation at room temperature. Wells were washed three times with the provided wash buffer. Streptavidin-HRP conjugate (100 µL) was added to each well and incubated for 30 minutes at room temperature. After additional washes, TMB substrate solution was added, and the plate was incubated in the dark for 30 minutes. The reaction was stopped by adding 100 µL of stop solution per well, and absorbance was measured at 450 nm using a microplate reader. Cytokine concentrations were calculated using standard curves generated with known concentrations of each cytokine.

### Western Blot analysis

At the end of the experimental time course, hCOs were harvested and washed twice with ice-cold phosphate-buffered saline (PBS) to remove residual media and debris. hCOs were then lysed using RIPA Lysis and Extraction Buffer (Thermo Scientific, Cat# 89901), supplemented with protease and phosphatase inhibitor cocktails according to the manufacturer’s instructions. Lysates were clarified by centrifugation at 14,000 × g for 15 minutes at 4 °C, subjected to SDS-PAGE using 4–20% gradient gels, and transferred to polyvinylidene difluoride (PVDF) membranes using a Western blotting system (Bio-Rad). Membranes were blocked with EveryBlot Blocking Buffer (Cat# 12010020, Bio-Rad) for 10 minutes, followed by incubation with specific primary antibodies at 4 °C overnight. After primary antibody incubation, membranes were washed with TBST and incubated with HRP-conjugated secondary antibodies at room temperature for 1 hour. Protein signals were visualized using the Clarity Max Western ECL Substrate system (Cat# 1705062, Bio-Rad). Images were acquired and analyzed using ImageJ software (National Institutes of Health, Bethesda, MD, USA). Protein expression levels (band intensities) were normalized to β-actin and expressed as fold change relative to the control group [26]. Details of the primary antibodies used are listed in Supplementary Table S1.

### Immunocytochemistry

The hCOs were fixed in 4% paraformaldehyde overnight at 4 °C and washed thrice with PBS. After fixation, the hCOs were dehydrated in 30% sucrose solution at 4 °C. Subsequently, the hCOs were embedded in Tissue-Tek O.C.T. compound (Cat# 4583) and frozen on dry ice. The frozen hCOs were sectioned into 10 μm slices using a cryostat and mounted onto glass microscope slides. The sections were blocked with blocking buffer containing 4% normal donkey serum (Cat# S-30-M) and 0.1% Triton X-100 in 1× PBS for 1 hour at room temperature. Next, the sections were incubated overnight at 4 °C with primary antibodies (1:500 dilution, listed in Supplementary Table S2) diluted in blocking buffer without Triton X-100. After three washes with PBS, the sections were incubated for 1 hour at room temperature in a light-protected humidified chamber with secondary antibodies diluted 1:1000: Alexa Fluor 488 (Donkey Anti-Mouse), Alexa Fluor 488 (Donkey Anti-Rabbit), Alexa Fluor 555 (Donkey Anti-Mouse), Alexa Fluor 555 (Donkey Anti-Rabbit), and Alexa Fluor 647 (Donkey Anti-Chicken). Finally, the slides were cover slipped with DAPI mounting medium (Invitrogen, Cat# P36935). Fluorescence signals were captured using a Carl Zeiss LSM 880 confocal microscope.

### Nitrite assay

After HIV-1 infection and cART treatment of hCOs, supernatants were collected, and nitrite levels were quantified using the Griess reagent kit (Cat# G7921). Briefly, 100 µL of supernatant was mixed with 100 µL of Griess reagent to generate a colorimetric reaction. After incubating for 30 minutes at room temperature, absorbance was measured at 540 nm. Nitrite concentrations were calculated using a sodium nitrite standard curve and expressed in µM.

### RT-PCR

Briefly, hCOs were homogenized, and RNA was extracted using Trizol reagent for semi-quantitative RT-PCR analysis. Total RNA was solubilized in RNase-free water and quantified in duplicate by measuring the optical density (OD) at 260 nm. RNA purity was confirmed by examining the OD260/OD280 ratio using a Nanodrop Microvolume Spectrophotometer (Invitrogen). For cDNA synthesis, 2 µg of RNA was reverse transcribed using the iScript™ Advanced cDNA Synthesis Kit (Cat# 1725038, Bio-Rad), following the manufacturer’s instructions. RT-qPCR was performed using diluted cDNA (1:20) and SYBR Green master mix. Samples were loaded into a 96-well PCR plate and run on a CFX96™ Real-Time System (Bio-Rad) with the following cycling conditions: initial denaturation at 95°C for 20–30 seconds (1 cycle), denaturation at 95°C for 1–3 seconds (40 cycles), annealing/extension at 60°C for 20 seconds (40 cycles), followed by a melt curve from 60°C to 95°C with gradual temperature increase (1 cycle). Primers were purchased from BioRad. Gene expression levels were calculated by subtracting the target genes’ cycle threshold (Ct) values from those of the housekeeping gene (GAPDH). Relative gene expression was calculated using the 2^-ΔΔCt^ method and reported as fold changes compared to the control group.

### Statistics

Statistical analyses were performed using GraphPad Prism version 8. Differences between the control, HIV-1 infected, and cART-treated groups (n =6-8 organoids per group, from three independent experiments), were evaluated by one-way ANOVA followed by Tukey’s multiple comparison test. For the rhesus macaque (RM) study, we used n = 4 animals, each from a different individual. Values are expressed as mean ± SD, with significance set at p < 0.05. Gene set enrichment analysis was conducted using 10,000 permutations; pathways with a Benjamini-Hochberg adjusted p-value < 0.05 and a minimum of two genes were considered significantly enriched.

## RESULTS

### Development, maturation, and differentiation of human cerebral organoids (hCOs): Stem cell markers, proliferation, and glial-neuronal characterization

The formation of hCOs is a complex process that begins with deriving cells from induced pluripotent stem cell lines (iPSCs). These cells self-organize to form distinct brain regions within a 3D structure [27, 28]. In this study, human iPSCs (# iPS15, Alstem, USA) were differentiated into hCOs using previously established protocols [28]. The development and maturation of brain organoids over 60 days were monitored using stem cell and proliferation markers (Fig. 1A-H). Early cultures displayed undifferentiated stem cell features, followed by the formation of embryoid bodies and progressive structural organization. By day 60, mature organoids exhibited well-defined tissue-like architecture with distinct internal compartments (Fig. 1A).

In parallel, we evaluated whether the established brain organoids accurately replicate key aspects of human brain development. To achieve this, we performed immunofluorescence staining for stem cell proliferation markers, including SOX2, PAX6, Ki67, FOXG1, and Nestin, which were observed on day 60. SOX2, PAX6, and Ki67 expression were developed in neural progenitor cells (NPCs) within the proliferative zones of the cerebral organoids, marking active neurogenesis and supporting the expansion and migration of neuronal precursors. At this stage, the organoids displayed well-defined neuroepithelial architecture, with expanding NPC populations driving early cortical-like patterning, as shown in Fig. 1B-D.

We next characterized neuronal and glial differentiation in hCOs by assessing lineage-specific markers. FOXG1 expression was detected in day 60 organoids, indicating neuronal maturation (Fig. 1E). The microglial marker Iba1 was expressed in the matured organoids, with Iba1⁺ cells predominantly localized to outer regions and displaying ramified, homeostatic morphologies (Fig. 1F). Astrocyte differentiation was confirmed by GFAP expression, with GFAP⁺ cells distributed in the organoids and exhibiting characteristic stellate morphology, consistent with the establishment of a supportive glial network (Fig. 1G). Neuronal differentiation was further assessed using Nestin and MAP2; Nestin expression marked neural progenitors, whereas MAP2 expression increased with maturation. MAP2⁺ neurons displayed elongated, branched processes indicative of functional neuronal maturation (Fig. 1H).

### Incorporation of HIV-1 ADA virus-infected microglia into hCOs

Our primary objective in developing the hCOs model is to investigate HIV-associated neuroinflammation in a system that closely mimics *in vivo* HIV pathogenesis. A key element of HIV-1 neuropathogenesis is the presence of HIV-infected human microglia derived from induced pluripotent stem cells (iPSCs) [29, 30]. HIV-1 exhibits extensive genetic variability, leading to significant differences among viral strains in their biological properties, including replicative capacity, syncytium-inducing potential, and tropism for specific target cells. Most primary isolates can infect both cell types; however, individual strains often display a distinct preference for either lymphocytes or macrophages, reflecting their coreceptor usage and influencing clinical progression and neuropathogenesis [31]. Among these, the HIV-1 ADA strain is a well-characterized, macrophage-tropic and neurovirulent subtype B isolate, frequently used as a model to study HIV-1 infection in the CNS and to evaluate antiretroviral drug efficacy in macrophage-based systems [21, 32].

To model this process under physiologically relevant conditions, we generated human microglia from induced pluripotent stem cells (iPSCs) and infected them with the macrophage-tropic, neurovirulent HIV-1 ADA strain. To replicate *in vivo* HIV neuropathogenesis, infected microglia were co-cultured with hCOs, and the cultures were maintained for 9 days post-infection (dpi), following treatment with cART (Fig. 1I). HIV-1 infection was confirmed through HIV p24 ELISA analysis (Fig. 1J-L), compared with control hCOs. As shown in Figure 1J, HIV p24 levels were significantly elevated in the infected hCOs on dpi 6 compared to earlier time points. The gradual increase in p24 protein levels across time reflects the progression of viral replication within the brain organoids. To evaluate the efficacy of cART, supernatants were collected every 3 days and analyzed for p24 levels by ELISA. cART treatment significantly reduced p24 antigen levels compared with HIV-1 infected controls (Fig. 1K-L). Immunostaining further confirmed that cART therapy markedly decreased TNF-α levels in HIV-1 infected hCOs relative to other groups (Fig. 1M).

### Single-cell transcriptional and chromatin accessibility changes in HIV-1 ADA-infected hCOs using integrated multiomic analysis

To evaluate multiplexed, long-term organoid modeling under HIV-1 infection and cART, samples were pooled across all conditions before single-cell encapsulation (Fig. 2A; Methods). We performed batch correction on both snATAC-seq and snRNA-seq datasets using the R package Harmony Integration. Subsequently, Seurat v5 and Signac v1.14 R packages were employed to characterize cellular composition in the snRNA-seq data, annotate cells based on transcriptional profiles, and guide the snATAC-seq analysis (Fig. 2B-C). The snRNA-seq analysis identified all major cell types within the cerebral organoids, using lineage-specific markers (Fig. 2B). Detected cell populations included neural stem cells (NSCs), midbrain dopaminergic neurons (mDNs), neurons, cycling radial glial cells (cRGCs), oligodendrocytes (oligos), astrocytes (astro), microglia, and oligodendrocyte precursor cells (OPCs). These populations were visualized using Uniform Manifold Approximation and Projection (UMAP; Fig. 2B, C).

**Figure 2.**
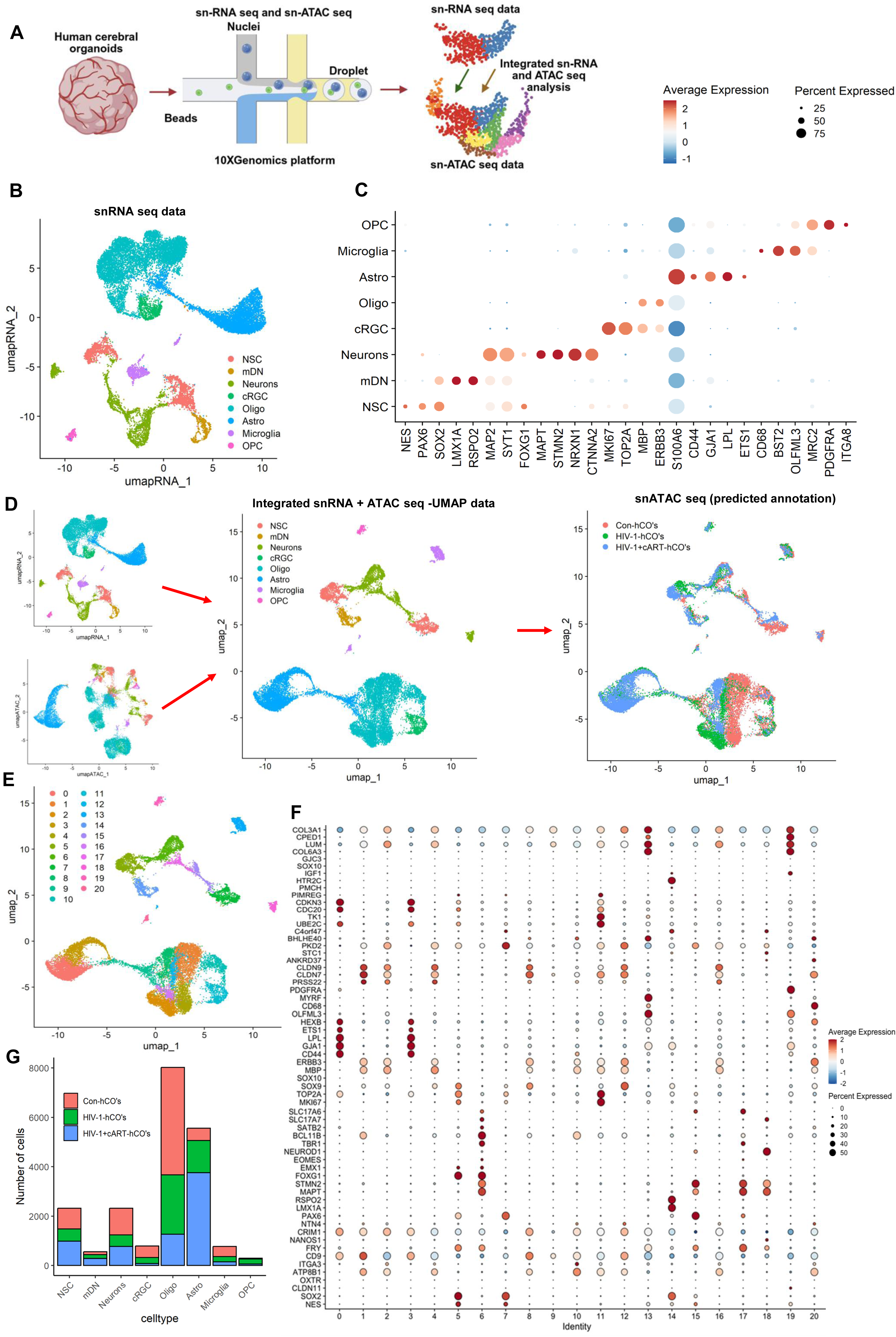
Single-cell transcriptional and chromatin accessibility profiling on HIV-1 infected hCOs with cART. **(A)** Graphical illustration of experimental methodology. Single-cell RNA sequencing (scRNA-seq) and ATAC-seq were performed on ∼60-day-old hCOs following HIV-1 infection with cART regimen (n = 6). **(B)** UMAP plots of snRNA-seq dataset. NSC- neural stem cells, mDN – midbrain dopaminergic neurons, Neurons, cRGC- cycling radial glial cells, oligo – oligodendrocytes, astro – astrocytes, microglia, OPC- oligodendrocyte progenitor cells. **(C)** Dot plot of snRNA-seq dataset showing gene expression patterns of cluster-enriched markers. The diameter of the dot corresponds to the proportion of cells expressing the indicated gene, and the density of the dot corresponds to the average expression across all cell types. **(D)** Multi-omics integration strategy for processing the snATAC-seq dataset. Following integrated snRNA-seq and snATAC-seq were filtered using a 97% prediction score threshold for cell-type assignment. **(E)** UMAP plot of snATAC-seq dataset with gene activities-based cell type markers. **(F)** The diameter of the dot corresponds to the proportion of cells with detected activity of the indicated gene, and the density of the dot corresponds to the average gene activity relative to all cell types. **(G)** Proportion of cell numbers shown in c, in different treatment conditions.

snATAC-seq captures the chromatin-accessibility profile of individual cells [33]. Since cell-type-specific chromatin accessibility remains understudied, we leveraged our annotated snRNA-seq dataset to predict cell identities in the snATAC-seq data using Seurat’s label-transfer framework. We generated a gene-activity matrix representing chromatin accessibility across gene bodies and promoters from the snATAC-seq data as the query, then identified transfer anchors between the snRNA-seq reference and this matrix. The resulting prediction scores indicated that most snATAC-seq cells were confidently matched to a single cell type (Fig. 2D). Comparison of these label-transfer predictions with curated annotations of unsupervised clusters (Fig. 2E, F) confirmed representation of all major cell types. It demonstrated that snATAC-seq provides resolution comparable to snRNA-seq for cell-type assignment. Interestingly, snATAC-seq identified 20 subpopulations within the cerebral organoid clusters (Fig. 2E), and the proportions of these clusters remained consistent across different treatment conditions (Fig. 2F).

Following HIV-1 infection, we observed notable shifts in cell type proportions, with microglia, astrocytes, oligodendrocytes, NSCs, and neurons comprising 27.8%, 23.5%, 30%, 21.4%, and 20.3% of the population, respectively, and a noticeable increase in OPCs and mDNs compared to uninfected controls (Fig. 2G).

For cross-species validation of the hCOs data, we reanalyzed previously published snRNA-seq datasets from healthy controls and SIV-infected, ART-treated rhesus macaques as described in the Materials and Methods (Fig. 3A) [23, 34]. The UMAP projection of aggregated cells are shown in Fig. 3B. The major cell populations identified include astrocytes, cerebellar glutamatergic neurons, choroid plexus epithelial cells, cortical projection neurons, cortical pyramidal neurons, GABAergic neurons, glutamatergic neurons, mature oligodendrocytes, microglia, myelinating oligodendrocytes, neurons, and OPCs (Fig. 3C). The canonical markers used for cell type annotations (Fig. 3D). Similar to HIV-1 infected hCOs, in the snRNA-seq data from the brains of SIV-infected, cART-treated RMs revealed considerable change in cell population composition compared to healthy controls.

**Figure 3.**
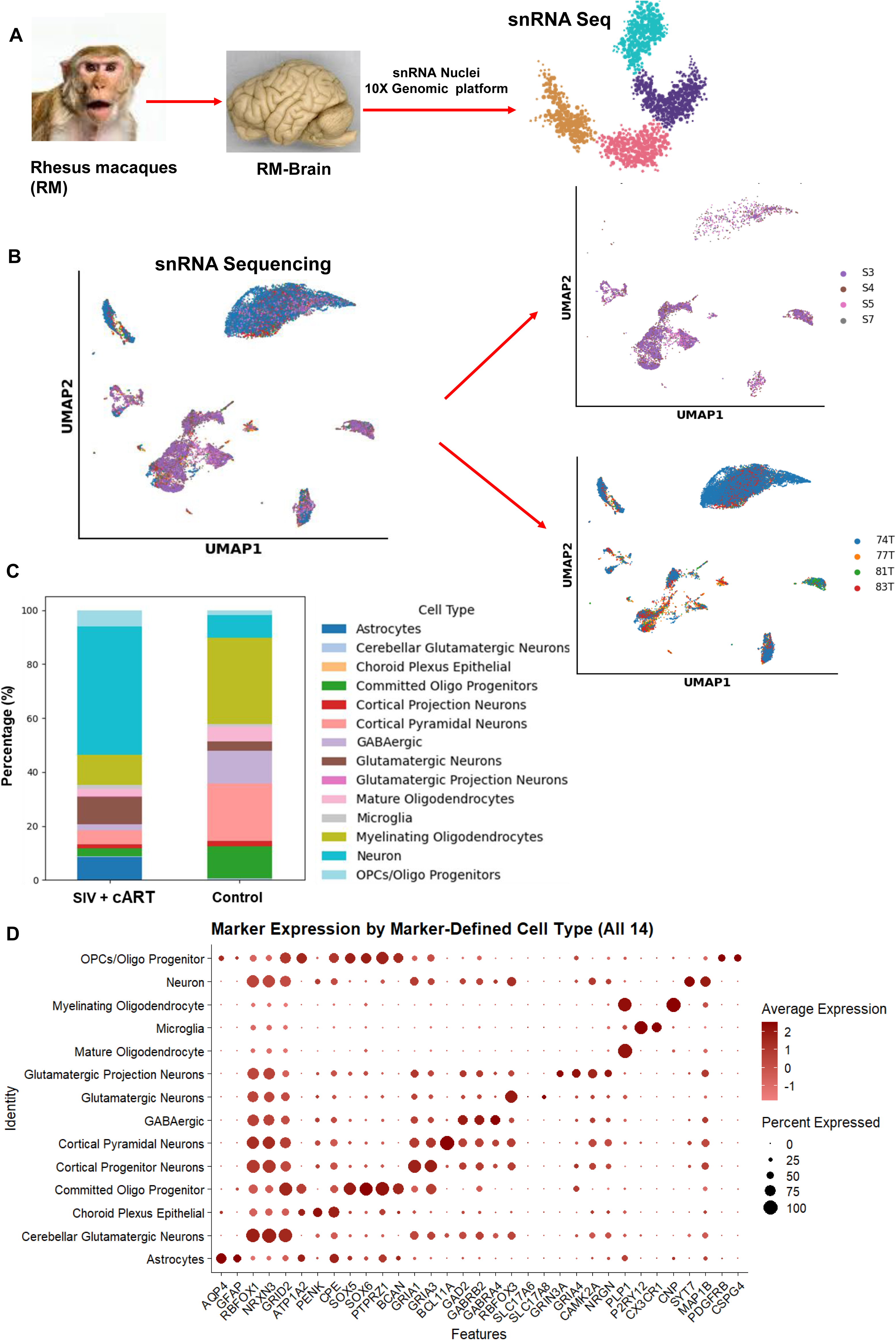
Single-cell transcriptomic profiling and cell type annotation in a macaque model of SIV infection. **(A)** Experimental schema. Four macaques were infected with 200 TCID50 of SIVmac251, and 5 weeks post-infection, a daily single subcutaneous injection (1 ml/kg body weight) of antiretroviral therapy (ART) was initiated. The ART regimen consisted of two reverse transcriptase inhibitors (FTC: 40 mg/ml and TFV: 20 mg/ml) and one integrase inhibitor (DTG: 2.5 mg/ml). After 30 weeks of ART, the animals were necropsied post perfusion with PBS containing 1 U/mL heparin to clear the blood from the brain. The brain tissues were then harvested and snap-frozen in LN2 for single-nuclei transcriptomic analysis. **(B)** UMAP plots illustrating transcriptional clustering of cells colored by annotated cell types. **(C)** Bar plot showing the proportion of identified cell types in SIV-infected, ART-treated rhesus macaques (RMs) and healthy controls. **(D)** Dot plot summarizing the expression of representative marker genes across clusters used for cell type annotations. The diameter of the dot corresponds to the proportion of cells expressing the indicated gene, and the density of the dot corresponds to the average expression across all cell types.

### Dual innate immune pathways drive robust HIV-1–associated inflammatory responses in hCOs and rhesus macaque brains

To assess the molecular impact of HIV-1 infection in hCOs, we performed differential gene expression analysis using fgsea (R)[35] across annotated cell types, comparing HIV-1-infected samples to uninfected controls and HIV-1 + cART-treated groups. This revealed widespread transcriptional changes across neurons and glial cells (Supplemental data file S1). DEG analysis revealed robust transcriptional changes across multiple cell types in response to HIV-1 infection. Remarkably, genes involved in immune activation and cellular stress were significantly upregulated, highlighting the persistent inflammatory state associated with HIV-1-neuropathogenesis. Enrichment analysis of the DEGs identified key biological pathways, including cytokine and interferon signaling, Toll-like receptor (TLR) signaling, mitochondrial electron transport, and glial cell development and differentiation, across all subtypes (Supplementary Fig. S1). These pathways are consistent with chronic immune activation and metabolic stress observed in HIV-1 infected hCOs. Mitochondrial dysfunction, reflected by changes in genes involved in electron transport, may underlie cellular energy deficits and oxidative stress. Furthermore, dysregulation of glial cell development pathways indicates potential disturbances in astrocyte and oligodendrocyte function, which may contribute to impaired neuro-glial homeostasis. The top 10 differentially expressed genes (ranked by adjusted p-value and fold change) in each major cell type are presented in Supplementary Figure S2A, providing a comprehensive view of the transcriptional alterations underlying these functional shifts.

Similarly, we performed single-nuclei ATAC-seq (snATAC-seq), and the data showed differential chromatin accessibility across conditions and cell types (Supplementary Fig. S2B) using fgsea in R and the Molecular Signatures Database (MSigDB) v2024.1. In microglia, top-enriched pathways included TLR signaling and TGF-β response and regulation, whereas astrocytes showed altered accessibility to genes related to synaptic signaling (Supplementary Fig. S2B). Volcano plots indicate that S100A8 and S100A9, known DAMP (damage-associated molecular pattern) genes, were significantly downregulated in HIV-1 infected hCOs across all cell types but were restored to near-control levels following cART treatment (Fig. 4A-F). In addition to several nuclear genes, MTRNR2L1, MTRNR2L6, AMBRA1, and ZNF90 were upregulated in HIV-1 infected hCOs. The altered expression of MTRNR2L1 and MTRNR2L6, genes likely reflecting a mitochondrial stress–adaptive response, results in disrupted oxidative phosphorylation. Treatment with cART significantly reduced this expression, particularly in astrocytes, oligodendrocytes, and cRGC (Fig. 4G-L).

**Figure 4.**
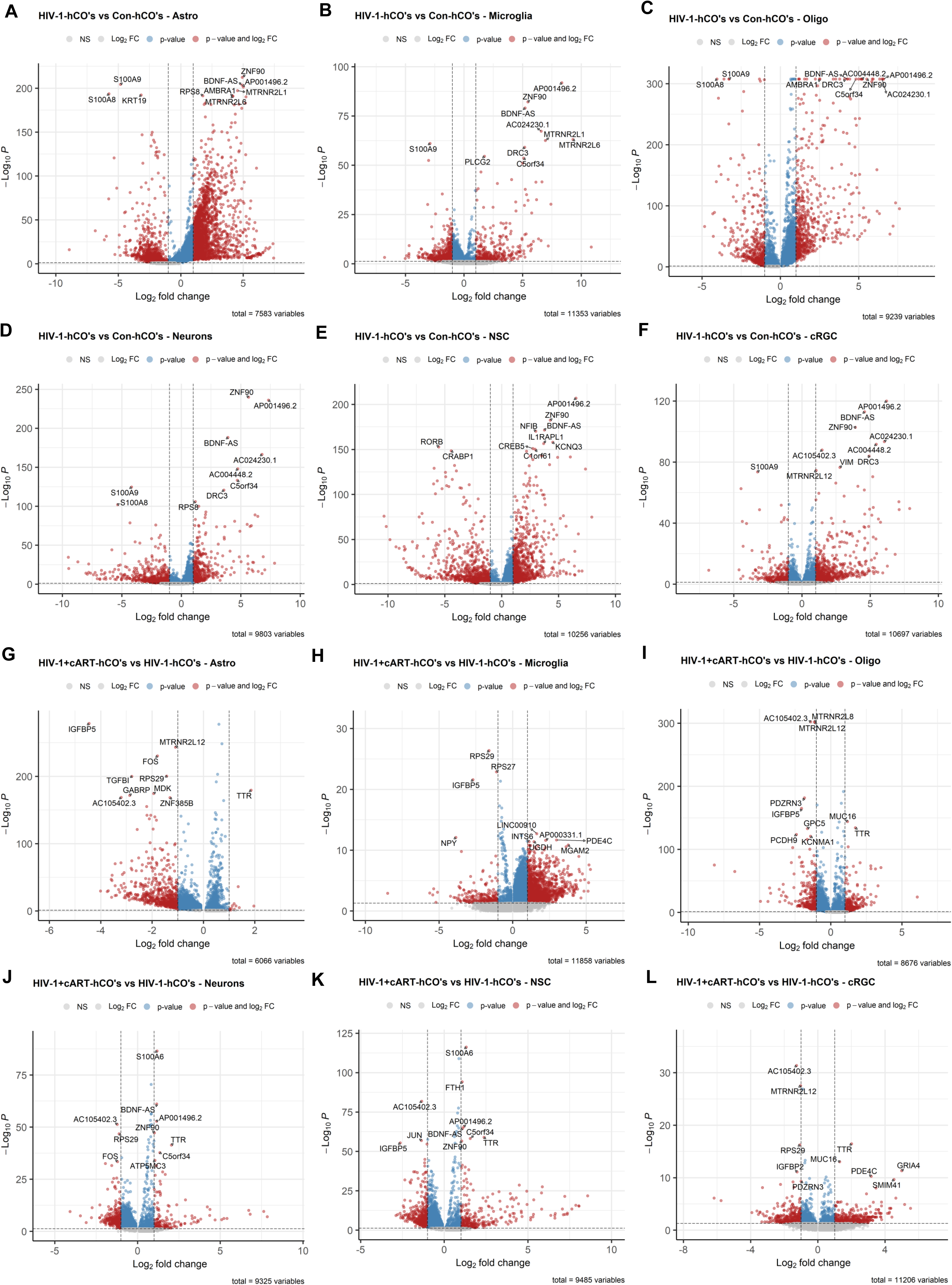
Transcriptional Profile of hCOs and HIV-1 infected hCOs. (A-L) Volcano plot of genes that are significantly (P ≤ 0.05; Wilcoxon rank sum test, Bonferroni adjusted) different when comparing HIV-1 infected vs. Control (A-F) and HIV-1 + cART vs. HIV-1 infection (G-L). X-axis displays the log2 fold-change of each significant gene.

KEGG analysis revealed significant dysregulation in multiple signaling cascades, including chemokine, TGF-β, Type I interferon, and oxidative phosphorylation pathways, when comparing HIV-1-infected hCOs to both control and cART-treated groups (Fig. 5A-D). Additionally, complementary GO enrichment analysis further indicated that key regulators of T-cell activation and inflammatory responses are selectively upregulated in infected hCOs (Supplemental data file S1). Notably, IL-17A, CEBPB, STAT1, FOXO3, and CSF1R are enriched in cytokine-cytokine receptor interaction (Supplementary Fig. S3A). Notably, IGF2R, previously linked to microglial activation and HIV-1 persistence in the CNS, was upregulated, while SLC11A1, a HAND-associated marker implicated in synaptic dysfunction and apoptosis, emerged as a key feature in our integrated multi-omics analysis (Supplemental data file S1). Remarkably, many DEGs in infected-hCOs mapped to antiviral defense, innate immune regulation, and interferon-mediated pathways, with a pronounced signature of proinflammatory cytokine and interferon-γ signaling observed in the current study (Supplementary Fig. S3B).

**Figure 5.**
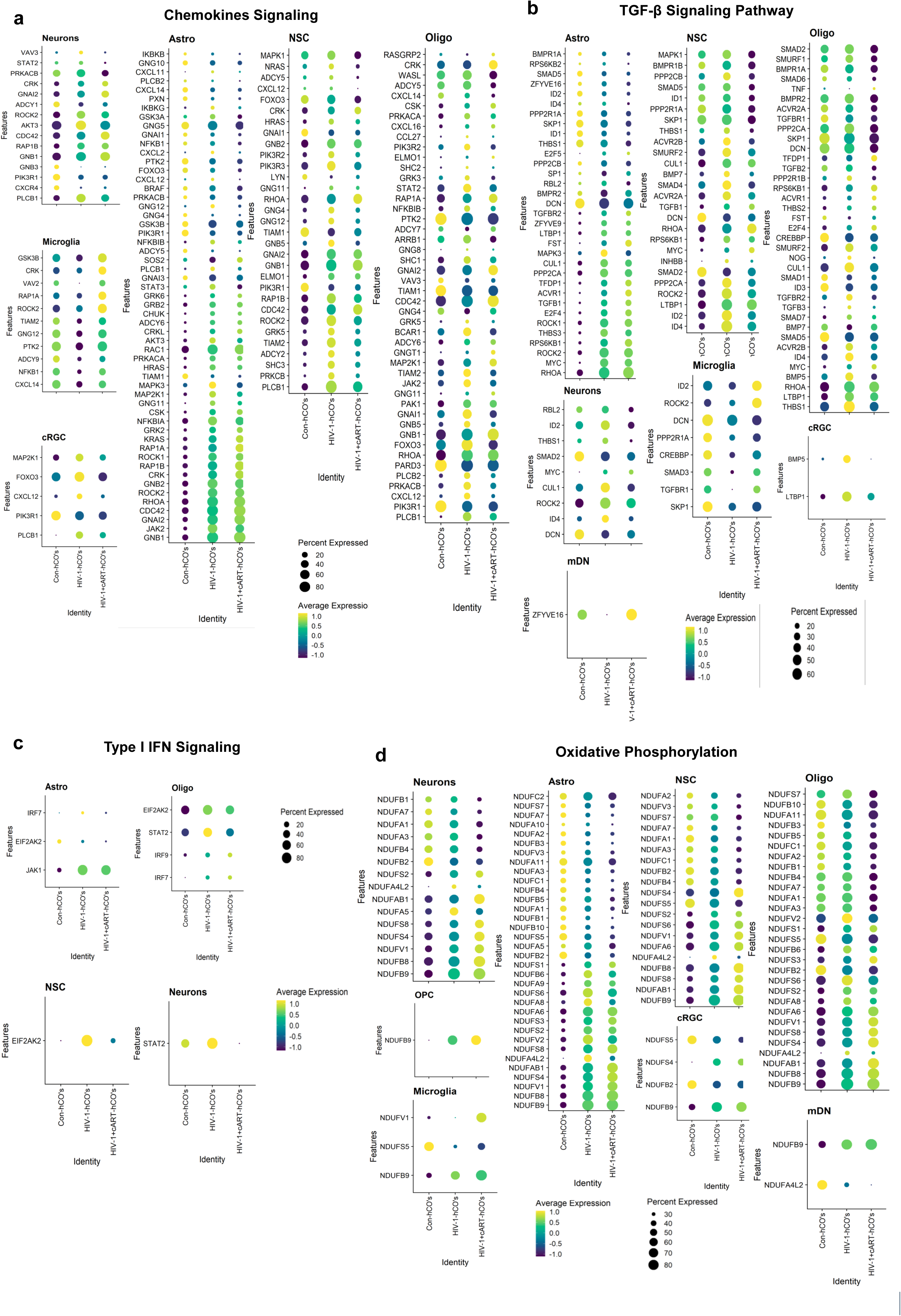
Functional enrichment analysis of upregulated genes in HIV-1 infected hCOs, highlighting key biological processes and pathways. **A-D** Kyoto Encyclopedia of Genes and Genomes (KEGG) pathways, offering a comprehensive view of the molecular mechanisms altered by HIV-1-infection. This analysis aids in identifying transcriptional regulators, cellular responses, and metabolic pathways affected by viral infection. **A** chemokines signaling, B TGF-β signaling pathways, **C** oxidative phosphorylation, and **D** Type I IFN signaling.

To extend these observations to the macaque model, we analyzed snRNA-seq data from SIV-infected cART-treated RMs and compared them to uninfected controls (Supplementary Fig. S4A-D). The analysis focused on immune-related pathways in chemokine signaling, TGF-β signaling, and Type I interferon signaling, with broader enrichment scans filtered by relevant keywords (Supplementary Fig. S5A-D). Additionally, pathways associated with immune responses, inflammatory responses, synaptic regulation, and the cell cycle are enriched across major cell types, as shown in Supplementary Figure S6. These findings parallel our observations in hCOs, reinforcing the presence of shared molecular disruptions in both human and macaque models of HIV/SIV neuropathogenesis.

To further elucidate transcriptomic inflammatory profiles changes, we performed differential peak accessibility analysis using the fgsea R package [35], identifying prominent transcriptional regulators such as NF-κB2, NEURL1, CEBPB, and STAT1 showing significant differential accessibility peaks in neurons and glial cells (Fig. 6). Additional analysis revealed that altered accessibility in immune-associated transcription factors, including STAT3, STAT4, IRF3, IRF9, and TFEB (Supplementary Fig. S7), was increased among all cell types. Moreover, we observed a significant reduction in S100A8 and S100A9 gene expression and chromatin accessibility in HIV-1 infected hCOs compared to other experimental groups, suggesting that these changes may exert anti-HIV effects in microglia, astrocytes, and oligodendrocytes (Supplementary Fig. S7). Together, our transcriptomic data reveal widespread, cell-type-specific immune activation and altered inflammatory response to HIV-1 which was inhibited by cART treatment. Recent studies using primary DCs provide evidence that cytosolic S100A9 suppresses HIV replication by inhibiting reverse transcription [36]. Consistent with this, mature monocyte-derived Langerhans cells exhibit reduced intracellular S100A9 levels and increased susceptibility to HIV-1 infection [37], underscoring that HIV-1 infection outcomes can be highly cell–type–dependent. However, the individual roles of S100A8/A9 proteins in regulating HIV-1 replication in primary cells remain poorly understood.

**Figure 6.**
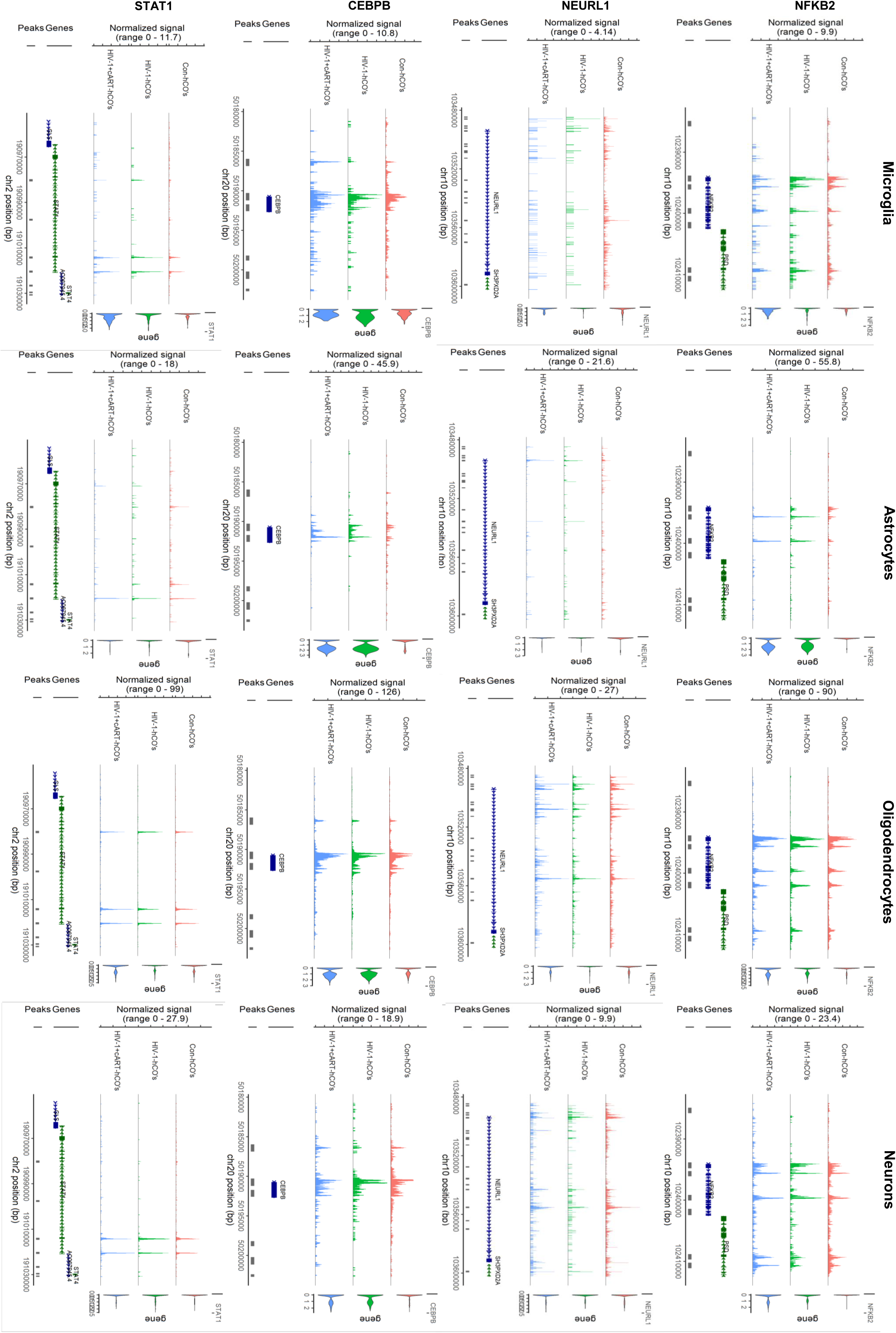
Single-nucleus ATAC-seq reveals differential chromatin accessibility and transcription factor regulation in HIV-infected brain organoids. Single-nucleus ATAC-seq was performed on cerebral organoids under control, HIV-1 infected, and cART-treated conditions to assess chromatin accessibility at the single-cell level. Peak intensities represent open chromatin regions, with higher peaks indicating greater accessibility. Analysis was performed across major cell types, including microglia, astrocytes, oligodendrocyte-lineage cells, and neurons. Differential peak analysis revealed dynamic changes in regulatory elements, particularly at loci associated with immune-related transcription factors, including NFKB2 (targeted gene locus on Chr 10), NEURL1 locus on Chr 10, CEBPB locus on Chr 20, and STAT1 locus on Chr 2. HIV-1 infection was associated with increased chromatin accessibility at NFKB2 and STAT1 loci, alongside reduced accessibility at the NEURL1 locus, suggesting impaired negative regulation of Notch signaling. Notably, CEBPB exhibited increased accessibility in myeloid-enriched clusters, aligning with inflammatory activation. Treatment with cART partially restored chromatin structure near NEURL1 and dampened accessibility near NFKB1, indicating modulation of inflammation-related regulatory programs. Gene peaks were annotated to nearby genes using reference genome coordinates (hg38), and cell-type–specific differences were visualized using integrated clustering and trajectory inference.

### Dynamic neuroinflammatory responses in HIV-1 infected hCOs: Biphasic activation following infection and cART treatment

Single-nuclei RNA-seq revealed that HIV-1 infected hCOs exhibited elevated expression of transcriptional regulators such as ATF4, CEBPB, and CREB5, as well as ISGs, including IFI6 and IFITM1 (Fig. 7A). To complement our hCOs data, we also examined single-cell transcriptomic data from the macaque model. This data revealed similar signatures, reinforcing the relevance of our finding across species (Fig. 7B). The overlap in cell-type-specific transcriptional changes, particularly in pathways related to inflammation, immune activation, and neuronal dysfunction, underscores a conserved response to viral infection and treatment. Consistent with our transcriptomic findings, qPCR validation further confirms a significant upregulation of proinflammatory cytokines, including TNF-α, IL-1β, IL-6, CEBPB, and C-X-C motif chemokine ligand 10 (CXCL10) levels were upregulated in HIV-1-infected hCOs compared to controls (Fig. 7C).

**Figure 7.**
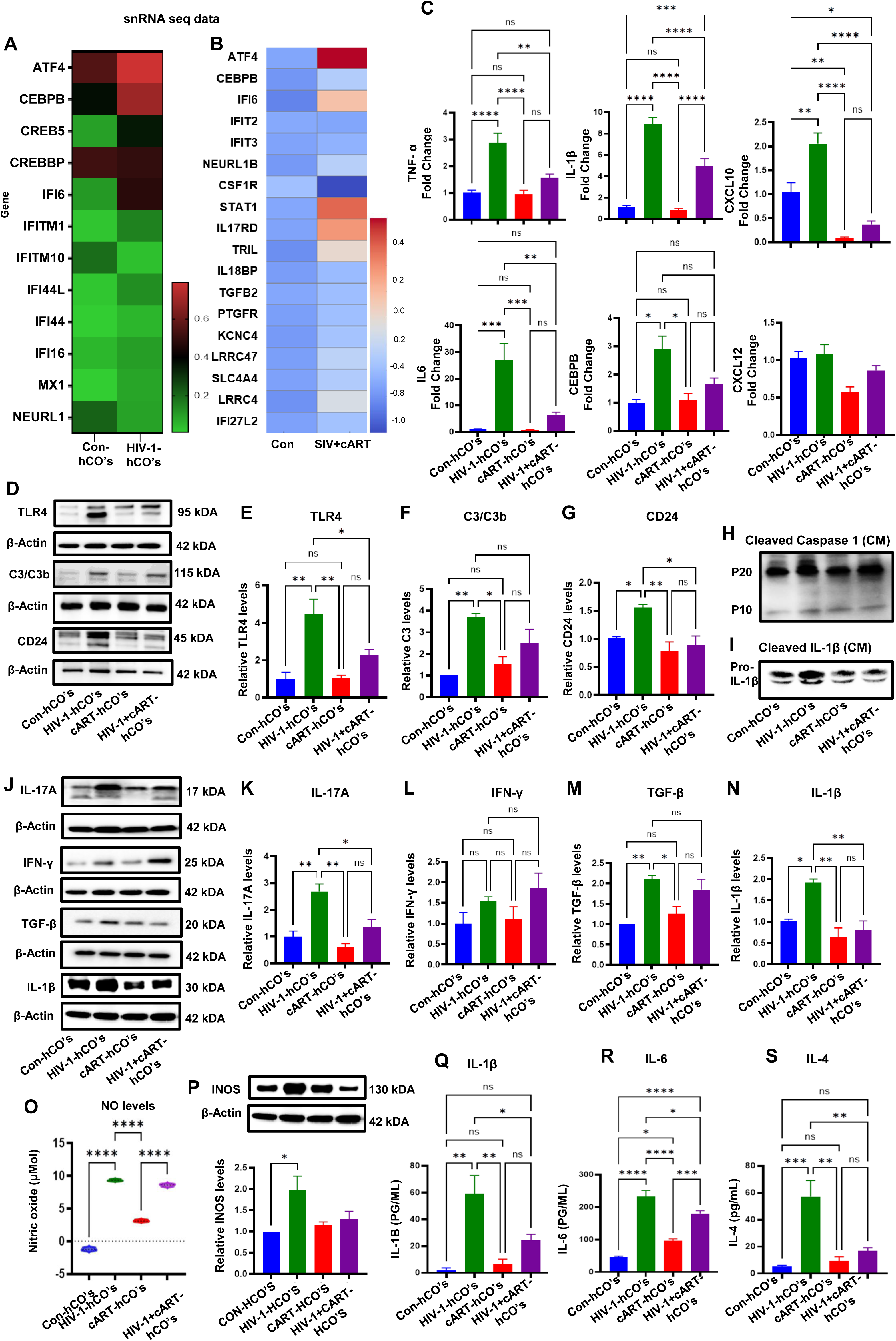
Exploration of HIV-1 infection dynamics in hCOs with and without cART treatment. **(A)** Heat map showing the expression of proinflammatory marker genes in HIV-1 infected hCOs vs Control hCOs using snRNA sequencing analysis. The graph represents log2FC ratios of [HIV-1-hCO/control]. **(B)** A curated panel of immune and inflammation-associated genes of interest, from control and SIV+cART treated RMs. Log-transformed, normalized expression values were averaged across samples per group and visualized using the heatmap R package. Row-wise scaling was applied, and a diverging color scheme (blue to red) highlights differential expression. **(C)** mRNA expression of pro-inflammatory cytokines, TNF-α, IL-1β, IL-6, CEBPB, CXCL10, CXCL12 were analyzed using Real time qPCR analysis in HIV-1 infected hCOs with or without cART. Fold changes were calculated using the formula 2-ΔΔCT method. **(D-G)** Immunoblot analysis was performed to assess the expression of key inflammatory and immune response proteins in HIV-1 infected hCOs with or without cART treatment. **(E)**TLR4, **(F)** C3/C3b, **(G)** CD24. **(H)** Cleaved caspase 1, **(I)** Cleaved IL-1β was measured from conditioned media of different groups. **(J)** Western blot, left panel, and the levels of **(K)** IL-17A, **(L)** IFN-γ, **(M)** TGF-β, **(N)** IL-1β, **(P)** iNOS, were quantified from 3 independent blots with respective bar graphs (right panels). Band intensities were measured and normalized to that of the respective β-actin. **(O)** Nitrite levels were quantified in supernatant media collected at day 6 post-infection from different experimental groups. **(Q-S)** Released level of IL-1β, IL-6 and IL-4 was further quantified using ELISA analysis. Data are represented as mean ± SD (n = 3). One-way ANOVA followed by Tukey’s multiple comparison test. \**P* < 0.05, \*\**P* < 0.01, \*\*\**P* < 0.001, \*\*\*\**P* < 0.0001 are represented as statistically significant, ns – Not significant.

Emerging studies have shown that Toll-like receptors (TLR) signaling in the brain initiates an innate immune response primarily through microglia and astrocytes, contributing to neuroinflammation [38]. Among TLRs, TLR4 is distinct in activating both MyD88- and TRIF-dependent pathways, resulting in amplified and prolonged inflammatory signaling. TLR4 is also more broadly expressed in CNS cell types and has been strongly associated with HIV-induced neuroinflammation [39], making it a significant focus of this study. Here, we investigate how TLR4 activation in microglia drives pro-inflammatory cascades and examine the role of complement pathways and immune mediators in regulating these responses. Western blot analysis shows the levels of TLR4, C3/C3b, and CD24 were significantly upregulated in HIV-1 infected hCOs, indicating a sustained neuroinflammatory response (Fig. 7D-G). Similarly, immunostaining data showed increased CD68 expression in HIV-1-infected hCOs (Supplementary Fig. S8), indicating dysregulated neuroimmune signaling and heightened microglial activation. Interestingly, 9-day cART treatment significantly reduced proinflammatory cytokine expression (Fig. 7C), TLR4 signaling, and CD24, an immune checkpoint molecule, in HIV-1–infected hCOs, while no significant changes were observed in the C3 complement system (Fig. 7D). These findings suggest that cART not only suppresses HIV-1 replication but also attenuates the inflammatory response associated with HIV-1 infection. The reduction in proinflammatory cytokine levels suggests that cART may help restore immune homeostasis and mitigate the neuroinflammatory effects seen in HAND [40, 41].

Furthermore, immunoblot revealed that HIV-1 infection robustly triggered neuroinflammatory responses in hCOs, as evidenced by significantly elevated levels of IL-17A, transforming growth factor-β (TGF-β), and interferon-γ (IFN-γ) (Fig. 7J-N). To further characterize the inflammatory milieu, we assessed levels of cleaved caspase-1 and cleaved interleukin-1β (IL-1β) in the supernatant collected from HIV-1 infected organoids with cART treatment on day 6 post-infection (dpi); both cleaved forms of caspase (P10 and P20) and cleaved IL-1β were significantly increased in HIV-1 infected hCOs compared to controls (Fig. 7H, I). Additionally, we evaluated nitrite and inducible nitric oxide synthase (iNOS) levels as indicators of reactive nitrogen species (RNS) production. Nitrite levels were significantly elevated in HIV-1-infected hCOs (Fig. 7O), and immunoblotting confirmed increased iNOS expression (Fig. 7P), indicating enhanced nitric oxide production and an amplified oxidative-stress response. Notably, treatment with cART significantly reduced IL-17A expression (Fig. 7K), underscoring its therapeutic potential to attenuate HIV-1-induced neuroinflammation and oxidative stress response and to reduce the risk of neurocognitive impairment [41].

ELISA analysis further reveals that protein levels of IL-1β, IL-6, and IL-4 were correspondingly increased during HIV-1 infection (Fig. 7Q-S). These analyses were shown in supernatant collected from HIV-1 infected organoids with cART treatment on day 6 post-infection. Consistent with our qPCR and Western blot findings, both assays confirmed the overexpression of these cytokines at the gene and protein levels. Notably, cART treatment helped restore homeostatic balance by reducing HIV-associated neuroinflammation and microglial activation, as demonstrated in this study.

### HIV-1 infection induces IL-17A-driven neuroinflammation and modulates CSF1R and STAT1 signaling in hCOs

The snRNA-seq data reveal that CCL2, STAT1 and 3, and SMAD-related genes were increased in HIV-1-infected hCOs (Supplementary Fig. S9A). Similarly, we examined the expression of CSF1R (colony-stimulating factor 1 receptor) and STAT1 (signal transducer and activator of transcription 1), both of which are key components of inflammatory signaling and immune response. We observed overexpression of phospho-STAT1(Y701) and CSF1R in HIV-1 infected hCOs compared to controls (Fig. 8A, B-D), suggesting that HIV-1 infection triggers IL-17A-mediated signaling through CSF1R and STAT1. Additionally, we observed that HIV-1 infection elicited a significant upregulation of DDX5 and IFI6 (Supplementary Fig. S9A&B) in hCOs, reflecting enhanced interferon-stimulated antiviral responses and increased RNA helicase activity associated with viral replication.

**Figure 8.**
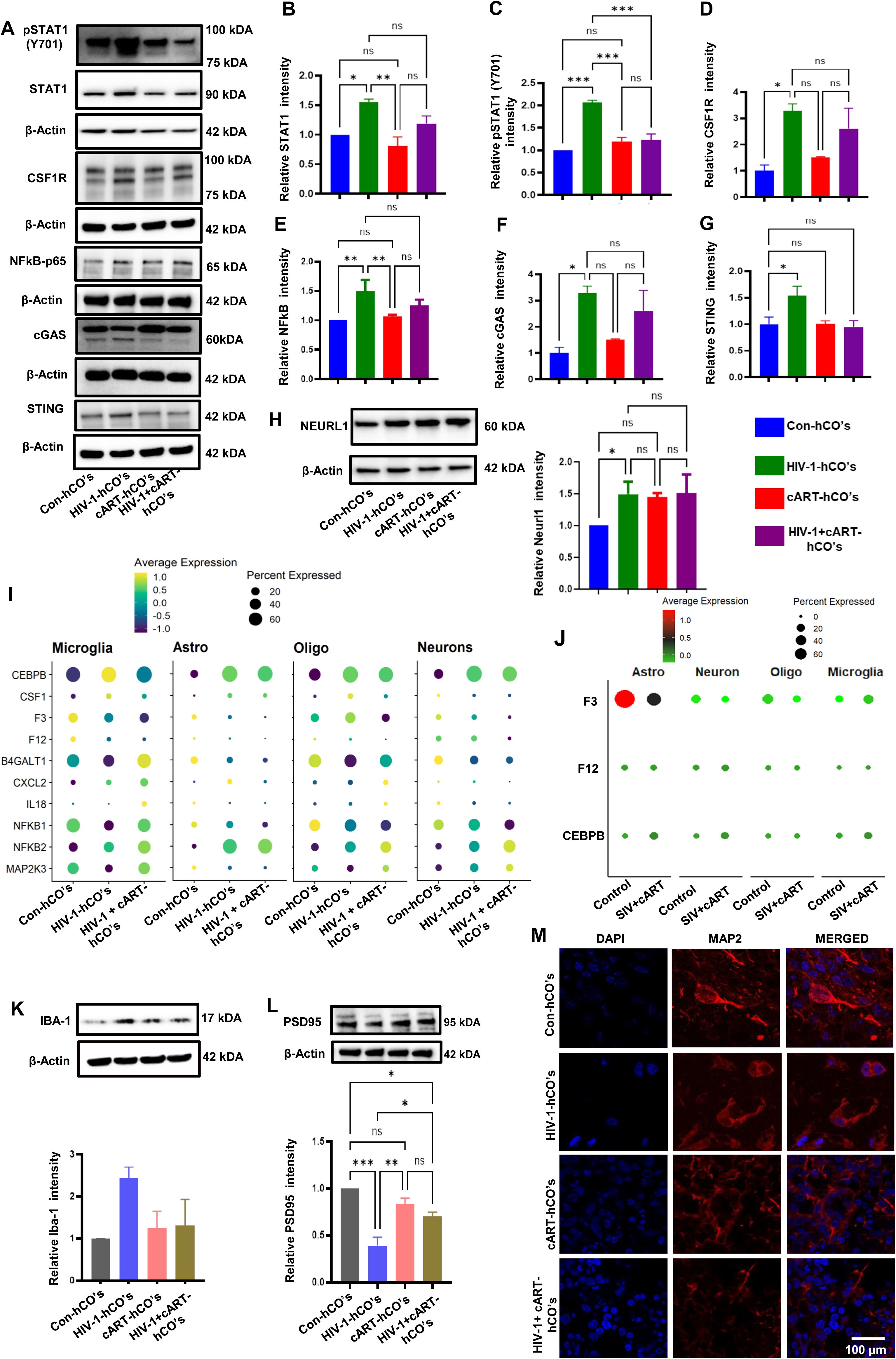
Impact of ART on STAT1 and cGAS/STING Signaling in HIV-1 infected brain organoids. **(A-G)** Western blot analysis (left panels) of key immune signaling molecules, including **(B)** STAT1, **(C)** p-STAT1 (Y701), **(D)** CSF1R, **(E)** NFKB-P65, **(F)** cGAS, and **(G)** STING, in HIV-1 infected organoids with or without cART treatment, the quantified band intensities are displayed in right panels. β-Actin served as an internal control. **(H)** Western blot analysis of NEURL1 expression in HIV-1 infected hCOs compared to control and cART, with respective bar graphs. β-Actin served as an internal control. **(I)** Integrated snRNA-seq and snATAC-seq analysis identified expression profiles of selected inflammatory markers across the various cell types among different groups. **(J)** snRNA-seq of F3, F12 and CEBPB gene expression were analyzed from control and SIV-infected cART-treated macaque brain samples. **(K)** Representative Western blot and bar graph show the quantification of Iba-1 levels in hCOs exposed to HIV-1 infection and cART treatment. Results were normalized with β-Actin. **(L)** Representative Western blot and bar graph show the quantification of PSD-95 subunit levels in hCOs exposed to HIV-1 infection and cART treatment. Results were normalized with β-Actin. **(M)** Immunolabeling fluorescence analysis was performed using antibodies against MAP2 (red). Scale bar = 100 µm. Data are presented as the mean ± SD of three independent experiments, with statistical significance determined by one-way ANOVA followed by Tukey’s multiple comparison test at \**p* < 0.05, \*\**p* < 0.01, \*\*\**p* < 0.001, ns- Not significant.

When HIV-1 infected hCOs were treated with cART, normalization of phospho-STAT1 expression was observed, suggesting that antiretroviral therapy can effectively reduce IL-17A levels. The normalization of these signaling pathways in the presence of cART highlights the therapeutic potential of antiretroviral drugs to reverse IL-17A-mediated neuroinflammation in HAND and to attenuate the downstream signaling cascade, NF-kB, via STAT1 activation (Fig. 8A, E).

Moreover, HIV-1 infection induced a marked upregulation of cyclic GMP-AMP synthase (cGAS) and stimulator of interferon genes (STING) in hCOs, indicating activation of the cytosolic DNA-sensing pathway [42]. Elevated expression of cGAS and STING in HIV-1 infected hCOs suggests that persistent sensing of viral DNA or damaged mitochondrial DNA may contribute to a sustained inflammatory milieu within the organoids. Importantly, cART treatment restored cGAS and STING expressions to levels approaching those of the control group, as shown in Fig. 8A, F-G.

Additionally, NEURL1, a newly identified gene in our study, exhibited significant transcriptional downregulation in HIV-1-infected hCOs (Fig. 7A), while paradoxically showing increased protein expression. No significant differences were detected following cART treatment (Fig. 8H). This discordance between protein abundance and transcriptional downregulation may suggest the involvement of post-transcriptional regulatory mechanisms, reflecting either a compensatory cellular response or HIV-driven translational dysregulation. Dot plot analysis of single-nucleus RNA-seq data from HIV-1–infected human cerebral organoids revealed cell-type–specific activation of innate immune and inflammatory pathways. Microglia exhibited the highest expression of inflammatory regulators, including CEBPB, CSF1, CXCL2, IL18, NFKB1, NFKB2, and MAP2K3, consistent with preferential infection and immune activation. Astrocytes and oligodendrocytes showed moderate induction, whereas neurons displayed lower expression, indicating secondary involvement (Fig. 8I). Validation in SIV-infected, cART-treated rhesus macaques demonstrated reduced inflammatory gene expression across all cell types, with residual microglial activation, confirming partial reversal of conserved neuroinflammatory programs by cART (Fig. 8J).

Indeed, to evaluate the functional implications of these molecular changes, we investigated markers of microglial activation, synaptic integrity, and neuronal health. Western blot and immunofluorescence analyses revealed increased Iba-1 expression, accompanied by marked reductions in PSD95, a postsynaptic density protein essential for synaptic signaling, and MAP2, a dendritic marker, in HIV-1 infected organoids (Fig. 8K-M). Notably, cART treatment restored the expression of both PSD95 and MAP2, suggesting that antiretroviral therapy supports healthy neuronal structure and function. Neurons, due to their high energy demands, are particularly susceptible to mitochondrial defects, which promote disease progression via oxidative stress, bioenergetic failure, and apoptosis [43, 44].

### cART attenuates HIV-1-induced mitochondrial damage and metabolic dysregulation in hCOs

Single nuclei RNA-seq data revealed an upregulation of several mitochondrial genes during HIV-1 infection, including three key NDUF family members (core subunits of mitochondrial Complex I) and six ATP synthase genes, suggesting heightened mitochondrial activity or stress responses in infected hCOs (Fig. 9A). Similar transcriptional patterns were observed in the SIV-infected, cART-treated macaque brains, where multiple Complex I subunits and ATP synthase genes were upregulated compared to uninfected controls (Fig. 9B). Despite this apparent upregulation of mitochondrial gene expression, we observed a significant downregulation of PGC1α, the transcriptional coactivator central to mitochondrial biogenesis and oxidative metabolism (Fig. 9C, D).

**Figure 9.**
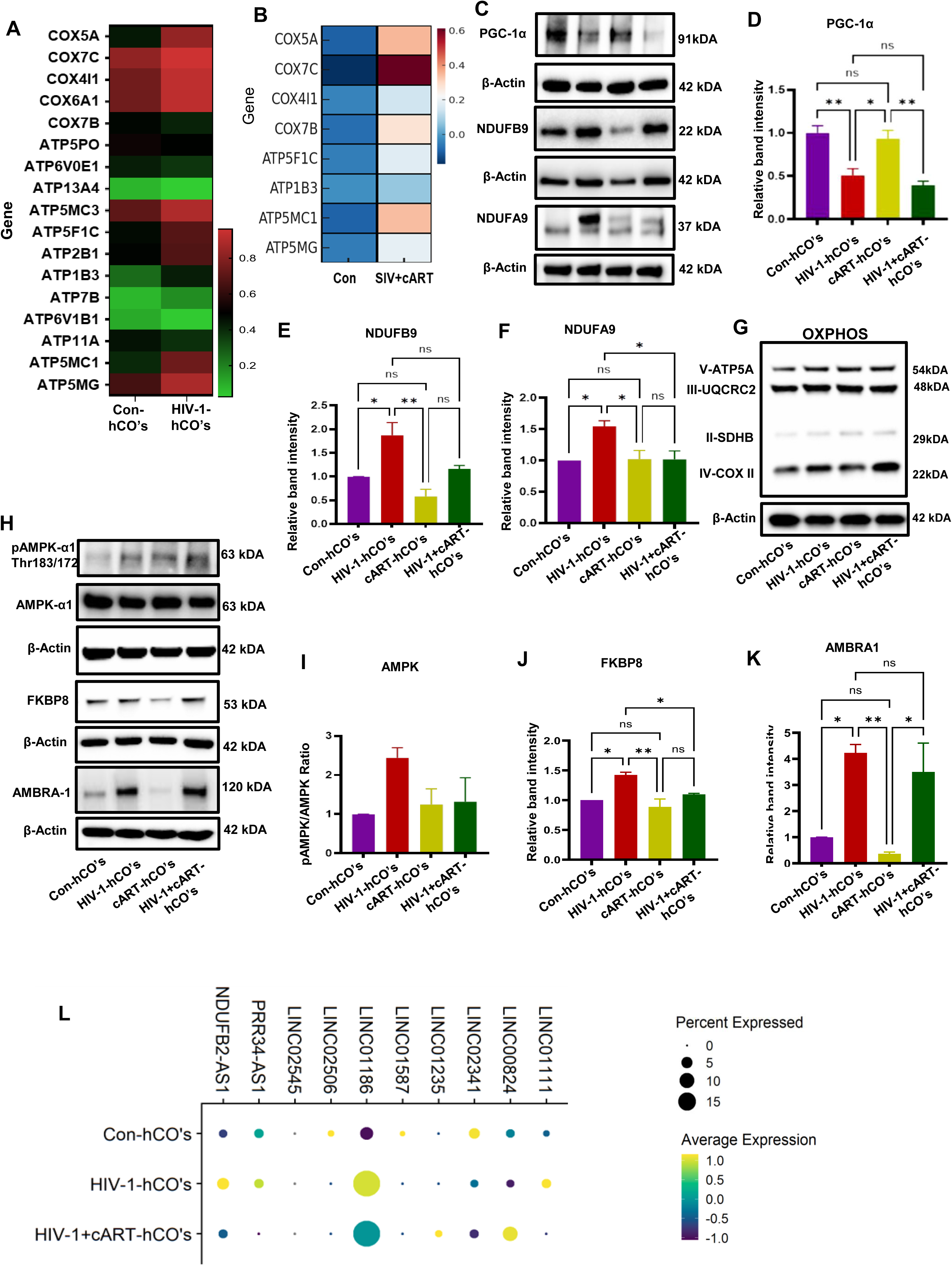
Mitochondrial complex protein and oxidative phosphorylation analysis in HIV-1 infected hCOs with and without cART. **(A)** Complex of V-related genes and ATP synthase subunit genes were analyzed using single-nuclei RNA seq using hCOs. **(B)** Heatmap was generated for mitochondrial and ATPase-related genes from control and SIV-infected and cART-treated macaques. **(C-K)** Right panels indicate the western blot analysis and respective bar graph on left panels, **(D)** PPARGC1α (PGC1α), **(E)** NDUFB9, **(F)** NDUFA9, **(G)** oxidative phosphorylation (OXPHOS) complex proteins, **(H)** Western blots analysis of AMPK and mitophagy related proteins **(I)** AMPK, **(J)** FKBP8, **(K)** AMBRA1 were analyzed from HIV-1 infected hCOs, with and without cART treatment. β-Actin served as an internal control. Data are presented as the mean ± SD of three independent experiments, with statistical significance determined by one-way ANOVA followed by Tukey’s multiple comparison test at \**p* < 0.05, \*\**p* < 0.01, ns- Not significant. **(L)** Integrated snRNA-seq and snATAC-seq analysis identified expression profiles of selected long intergenic non-coding RNAs (LINC RNAs) within microglial populations.

Interestingly, protein-level analysis confirmed elevated expression of NDUFA9 and NDUFB9, indicating mitochondrial Complex I overactivation, a condition linked to reactive oxygen species (ROS) production, cellular stress, and apoptosis in HIV-1 infected hCOs (Fig. 9E-F). These changes were accompanied by increased levels of phosphorylated AMPKα at Thr172, a metabolic stress sensor activated during energy deficits (Fig. 9H, I). Remarkably, cART treatment attenuated the expression of NDUFA9, NDUFB9, and phospho-AMPKα, suggesting a partial restoration of mitochondrial function and metabolic homeostasis.

To investigate whether oxidative phosphorylation triggers mitophagy during HIV-1 infection, we examined the expression of FKBP8 and AMBRA1 with or without cART treatment. Both FKBP8 and AMBRA1 proteins were significantly upregulated in HIV-1–infected hCOs; however, cART treatment did not significantly alter AMBRA1 expression (Fig. 9H, J-K) or oxidative phosphorylation levels (Fig. 9G).

Similarly, snRNA-seq data identified several long non-coding RNAs (lncRNAs) that could serve as therapeutic targets in disease models. Specifically, lncRNAs such as NDUFB2-AS1, PRR34-AS1, LINC02545, LINC02506, LINC01186, LINC01587, LINC01235, LINC02341, LINC00824, and LINC01111 were identified from HIV-1-infected hCOs within the microglia subtypes (Fig. 8L) and other cell types, as shown in Supplementary Fig. S10. These lncRNAs are implicated in various biological processes, including mitochondrial functions and glycolytic reprogramming [26] as well as NF-kB signaling. Due to their regulatory roles, they represent promising candidates for further investigation and potential alternative targets for cART treatment. LINC01186 and NDUFB9-AS1 were markedly upregulated in HIV-1 infected hCOs, with their expression attenuated by cART. In contrast, LINC00824 and LINC02341 were downregulated following infection, but cART treatment restored these effects (Fig. 9L).

### HIV-1 infection induces tau pathology and BIN1 upregulation: Special reference to HAND

Disease Ontology (DO) enrichment analysis of microglial subtypes in HIV-1 infected hCOs revealed significant associations with Alzheimer’s disease, neurodegeneration pathways (including Prion and Parkinson’s diseases), and the AGE-RAGE signaling cascade linked to diabetic complications (Fig. 10A). As key effectors of the CNS immune response to HIV-1, the microglia also showed upregulation of several KEGG-enriched genes, TUBB6, TUBA3C, ATP5MC1, COX5A, NDUFB9, SDHC, PSMA8, DDIT3, AMBRA1, NOX4, and WNT pathway components, suggesting a heightened susceptibility to Alzheimer-like neuropathogenesis (Fig. 10B). To further explore the neuropathological impact of HIV-1 infection in the context of HAND, we examined Tau accumulation, and BIN1 expression in hCOs. HIV-1 infection showed a significant increase in total and phosphorylated Tau levels compared to controls, suggesting that HIV-1 infection disrupts tau homeostasis and increases Tau oligomerization, which might contribute to neurotoxic outcomes associated with HAND (Fig. 10C). SynGO enrichment analysis revealed significant overrepresentation of synapse-related cellular components and biological processes. Enriched terms included presynapse, postsynapse, synapse, and postsynaptic density, along with presynaptic endocytic zone and synaptic vesicle membrane. Biological processes regulating postsynaptic organization and maintenance were enriched, indicating disrupted synaptic architecture, plasticity, and neuronal communication (Fig. 10D). Similarly, immunostaining data revealed increased phospho-Tau accumulation and BIN-1 expression in HIV-1 infected hCOs (Fig. 10E, F). This suggests a possible link between HIV-1-induced molecular changes and Tau-related pathology. Since BIN1 can alter intracellular Tau dynamics, its upregulation may contribute to neuronal damage, increased neuroinflammation, and synaptic dysfunction in the context of HIV [45]. Treatment with cART significantly reduced Tau accumulation and oligomerization, and restored BIN1 levels. These results are consistent with previous studies that suggested HIV-1 infection as a potential risk factor for neurodegeneration and cognitive decline, including the acceleration of AD-like pathology in the brain [46, 47].

**Figure 10.**
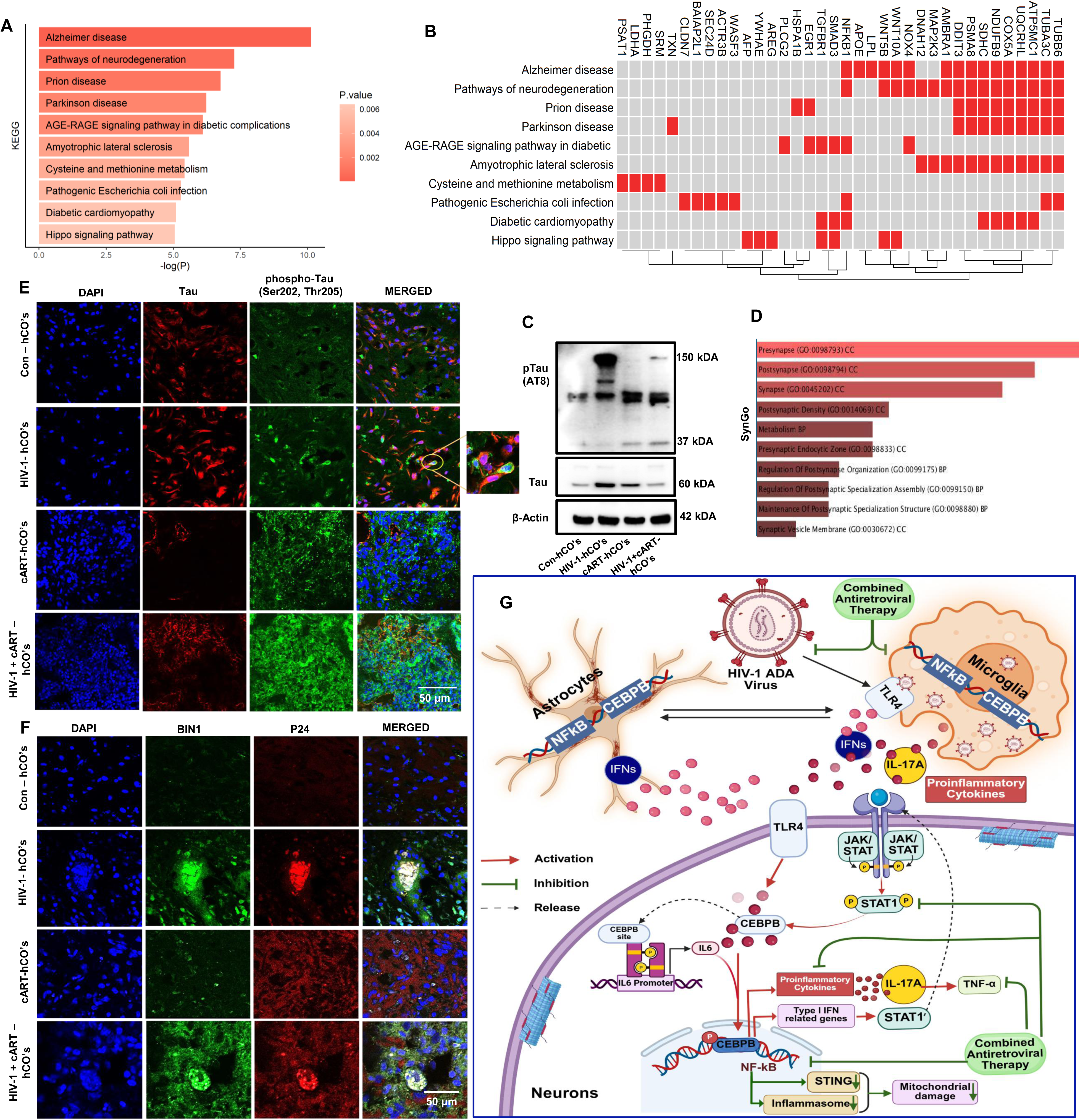
HIV-infected brain organoids induce Tau pathology and its potential link to HAND. (**A)** enriched KEGG pathway. (**B)** KEGG enrichment of the Top signal pathway in HIV infected hCOs within the microglia populations. (**C)** Western blot analysis of phospho-Tau and total Tau protein levels in HIV-infected organoids. Results indicate that Tau phosphorylation was elevated in HIV-infected hCOs but suppressed by ART treatment, suggesting a potential link between HIV infection and neurodegenerative processes. (**D)** SynGO enrichment analysis of synaptic gene signatures. Bar plot showing significantly enriched SynGO terms derived from differentially expressed genes. Enriched terms are grouped into Cellular Component (CC) and Biological Process (BP) categories. Prominent enrichment of presynaptic and postsynaptic compartments, including presynapse, postsynapse, synapse, and postsynaptic density, indicates extensive synaptic involvement. Bar length represents the degree of enrichment (–log10 adjusted p value). Only significantly enriched SynGO terms are shown. **(E)** Immunofluorescence imaging of Tau pathology in HIV-infected cerebral organoids. Double immunolabeling was conducted using antibodies against phospho-Tau (Ser 202 and Thr 205, AT8) (green) and total Tau (red) in HIV-1_hCOs with or without cART treatment, revealing the presence of hyperphosphorylated Tau. Scale bar = 50 μm. (**F)** Representative immunofluorescence image showing overexpression of BIN-1 in HIV- infected cerebral brain organoids. Double immunolabeling fluorescence analysis was performed using antibodies against BIN-1 (green) and HIV-1 p24 (red) in HIV-1-infected hCOs in the presence or absence of cART. Scale bar =50μm. (**G**) Schematic representation of HIV-1-induced neuroinflammatory signaling in microglia, astrocytes, and neurons, and the modulatory effects of cART. HIV-1 ADA virus infects microglia and astrocytes, triggering TLR4-mediated activation of transcription factors CEBPB and NF-κB. In microglia, this leads to the production and release of IL-17A, interferons and other proinflammatory cytokines. These inflammatory mediators activate TLR4 and JAK/STAT signaling pathway in neighboring neurons, leading to STAT1 phosphorylation and nuclear translocation. Astrocytes also contribute to the inflammatory milieu through CEBPB and NF-κB signaling and IFN release. IL-17A and TNF-α enhance inflammatory signaling cascades, perpetuating neuronal dysfunction. Combined antiretroviral therapy (cART) attenuates HIV-1 infection and inhibits downstream inflammatory signaling, including the release of IL-17A, TNF-α, and activation of NF-κB and STAT1. Arrows indicate activation (red), inhibition (green), and release or secretion (dashed black).

## DISCUSSION

HIV-associated neuroinflammation remains a major contributor to cognitive decline despite effective combined antiretroviral therapy (cART). The molecular pathways linking viral infection, glial activation, and neuronal dysfunction remain incompletely understood. In this study, we identify the novel mechanism of the IL-17A/CEBPB signaling axis and NEURL1 as underappreciated regulators of HIV-induced CNS dysfunction. Using human cerebral organoids (hCOs) containing microglia, along with cross-species validation in SIV-infected, ART-treated rhesus macaques, we delineate a mechanistic framework linking microglial activation, cytokine induction, synaptic impairment, mitochondrial dysfunction, and Tau pathology. Our findings highlight molecular connections that sustain CNS injury in HIV-1 and demonstrate that cART can reverse critical inflammatory and metabolic perturbations, underscoring actionable therapeutic targets.

HIV-induced neuroinflammation begins with chronic microglial activation, which rapidly propagates to astrocytes, neurons, and oligodendrocytes. This disruption of multicellular signaling networks dismantles homeostatic signaling networks, leading to neuronal dysfunction and progressive cognitive decline. Although cell-type-specific abnormalities have been documented, the integrated signaling cascades linking these effects to clinical outcomes remain poorly defined. To address this gap, we introduced HIV-1–infected iPSC-derived microglia into mature (60-day-old) cerebral organoids using a surface-seeding strategy. This approach allowed us to investigate microglia-mediated immune modulation within a physiologically mature neural environment. By focusing on the role of peripherally infected cells in seeding the brain reservoir [25, 48, 49], our model achieves high biological relevance and closely approximates the actual pathophysiology of HIV in humans.

Integration of snRNA-seq and snATAC-seq, together with publicly available datasets from macaque brains [23, 34] provided a single-cell resolution of autonomous and intercellular responses. This approach supported mapping cell composition, infection- and therapy-dependent transcriptional programs, developmental trajectories, chromatin accessibility, and active regulatory elements. Notably, cART mitigated many HIV-driven perturbations, particularly within glia–neuron networks, suggesting mechanisms by which therapy preserves CNS homeostasis.

HIV-1 infected microglia exhibited progressive increases in intracellular p24 antigen by day 6 post-infection, indicating active viral replication and the initiation of neuroinflammation. The increased p24 expression was accompanied by strong upregulation of pro-inflammatory cytokines (IL-1β, TNF-α, IL-6) and chemokines, including CXCL10 and MCP-1, consistent with previous reports of HIV-mediated neuronal injury [50]. Our hCOs recapitulated these features, exhibiting pronounced induction of CXCL10, CEBPB, and IFN-γ following HIV-1 infection. IFN-γ, in turn, activated canonical JAK/STAT signaling by phosphorylating STAT1 and driving its nuclear translocation, thereby driving ISG expression [51]. Importantly, cART treatment reduced STAT1 phosphorylation and suppressed downstream cytokine and chemokine expression, highlighting its therapeutic role in mitigating HIV-1–induced neuroinflammatory cascades.

The transcription factor CEBPB emerged as a central regulator of HIV-1–induced neuroinflammation. A key mediator of immune and inflammatory gene expression [52], CEBPB was robustly upregulated in both HIV-1–infected hCOs and SIV-infected macaque brains. CEBPB is modulated by IL-17A, a Th17-derived cytokine that engages NF-κB and MAPK signaling through its SEFIR domain [53]. This IL-17A/CEBPB axis has been implicated in autoimmune neuroinflammatory conditions, such as experimental autoimmune encephalomyelitis [10]. Previous studies demonstrate that HIV-1 activates the complement C3 promoter through NF-κB–mediated upregulation of IL-6, enhancing inflammatory processes associated with HAND [54].

Emerging evidence suggests that IL-17A enhances glial reactivity, while CEBPB integrates JAK/STAT and interferon signaling, driving sustained immune dysregulation [55]. Dysregulated CEBPB expression may therefore contribute to sustained neuroinflammation in HIV-1–infected individuals, even under effective cART. Consistent with this, HIV-1–induced NF-κB activation also upregulates CSF1R, which remains persistently expressed in the brains of virally suppressed SIV- and HIV-infected individuals, indicating that CSF1-mediated inflammation persists despite viral suppression [56, 57]. Consistent with these findings, CSF1R expression remained elevated in our brain organoid model following cART-mediated viral suppression, suggesting that microglia may adopt either a protective (M2) or pro-inflammatory (M1) phenotype depending on disease context [58]. In chronic neurodegeneration, such as HIV infection, the shift toward a protective M2 state appears impaired, contributing to sustained neuroinflammation and neuronal injury.

Notably, NEURL1, a neuronal E3 ubiquitin ligase involved in synaptic stability, was downregulated under these inflammatory conditions, suggesting that microglial CSF1R activation may indirectly influence neuronal homeostasis. Loss of NEURL1 may amplify IL-17A/STAT1 signaling and destabilize synaptic networks, consistent with studies showing that NEURL1 deficiency impairs memory and neuronal function [59]. Although a direct mechanistic link between NEURL1 and CEBPE remains unestablished, our data suggest that NEURL1 may regulate transcription by modulating upstream effectors post-translationally. Importantly, cART treatment restored NEURL1 genes and key synaptic markers (PSD95, MAP2), highlighting its ability to re-establish neuronal homeostasis amid ongoing neuroimmune stress. Collectively, persistent CSF1R signaling and altered NEURL1 expression contribute to the dysregulated neuroimmune environment in HIV-associated CNS injury, positioning NEURL1 as a key integrator of inflammatory signaling and synaptic integrity with therapeutic potential for HAND (Fig. 10G).

Consistent with neuropathology in other neurodegenerative diseases such as Alzheimer’s and Parkinson’s, HIV-1-infected hCOs exhibited pronounced mitochondrial dysfunction [60, 61]. The altered mitochondrial impairments and oxidative response were evidenced by elevated ROS, increased expression of electron transport chain (ETC) Complex I subunits (NDUFB9, NDUFA9), and activation of apoptotic pathways [26]. Treatment with cART reduced CEBPB levels, upregulated PGC1α, restored ETC function, and decreased ROS, confirming the interconnected roles of CEBPB and mitochondrial health. Notably, CEBPB regulates APOE expression and preferentially promotes the ApoE4 isoform, an established genetic risk factor for Alzheimer’s disease [62]. These findings suggest that CEBPB upregulation not only promotes HIV-induced inflammation but may also intersect with neurodegenerative pathways.

Tau pathology further underscores the convergence of HIV-1 infection and neurodegeneration. Inflammatory mediators and viral proteins can promote Tau phosphorylation, which contributes to synaptic dysfunction and HAND-like pathology [63]. The adaptor protein BIN1, a Tau interactor, links Tau phosphorylation to calcium homeostasis and synaptic regulation [64, 65]. In our study, cART treatment reduced Tau aggregation in HIV-1 infected hCOs, consistent with reports that nucleoside reverse transcriptase inhibitors such as lamivudine (3TC) mitigate Tau-driven neurodegeneration [66]. Taken together, these findings suggest that antiretroviral therapy may confer direct neuroprotective effects beyond viral suppression.

## Conclusion

This study identifies a previously unrecognized dual regulatory mechanism by which HIV-1 drives CNS dysfunction, simultaneously increasing neuroinflammation and compromising neuronal integrity through CEBPB upregulation and NEURL1 suppression. Our studies suggest that the CEBPB acts as a central integrator of IL-17A/STAT inflammatory signaling with mitochondrial dysfunction, linking ROS accumulation, complement activation, and Tau pathology to progressive neuronal injury. In parallel, loss of NEURL1 emerges as a novel driver of synaptic destabilization and cognitive vulnerability in the HIV-infected brain. Importantly, cART not only suppresses inflammatory signaling but restores nuclear-mitochondrial and synaptic regulatory pathways, as reflected by reduced CEBPB, recovery of NEURL1 expression, improved mitochondrial homeostasis, and rescue of synaptic markers. Collectively, these findings define the IL-17A-CEBPB–NEURL1 axis as a previously undefined mechanistic link between immune activation, mitochondrial stress, and synaptic failure in HIV-associated neurocognitive dysfunction, highlighting actionable therapeutic targets to preserve neuronal resilience and cognition in people living with HIV.

## Supporting information

Excel file, Full blot, Tables

## Supplementary material

## Supplementary files available online

Materials and Methods

Fig. S1 to S10

Table S1 and S2

Figure legends S1 - S10

Full blot images Fig. 7 – Fig.9

Other supplementary material for this manuscript includes the following:

Supplemental data file S1

## Acknowledgements

The authors thank Dr. Carlton Anderson and Dr. Gabrielle Cannon at the University of North Carolina Advanced Analytics Core Facility for assistance with single-nucleus RNA and ATAC sequencing. We also thank Dr. Mayur Doke for his valuable support with the organoid studies. We also thank Hemavathi Iyyappan for her assistance in preparing the supplementary tables. The Image Analysis Laboratory, College of Veterinary Medicine and Biomedical Sciences, Texas A&M University, is acknowledged for providing access to the Carl Zeiss LSM 880 confocal microscope.

## Author contributions

T.S. and K.C. designed and performed experiments, analyzed data, and drafted the manuscript. K.C. and M.S. performed Western blot experiments. J.C., S.G., and T.S. analyzed data and contributed to manuscript editing. A.A. and O.I. performed experiments and analyzed data. T.S., S.B., K.C., and J.C. provided critical scientific input. S.B. designed experiments and edited the manuscript. T.S. conceptualized, supervised, and supported the overall study, designed experiments, analyzed data, and co-wrote the manuscript. All authors read and approved the final version of the manuscript.

## Supplementary figure

**Supplementary figure S1:**
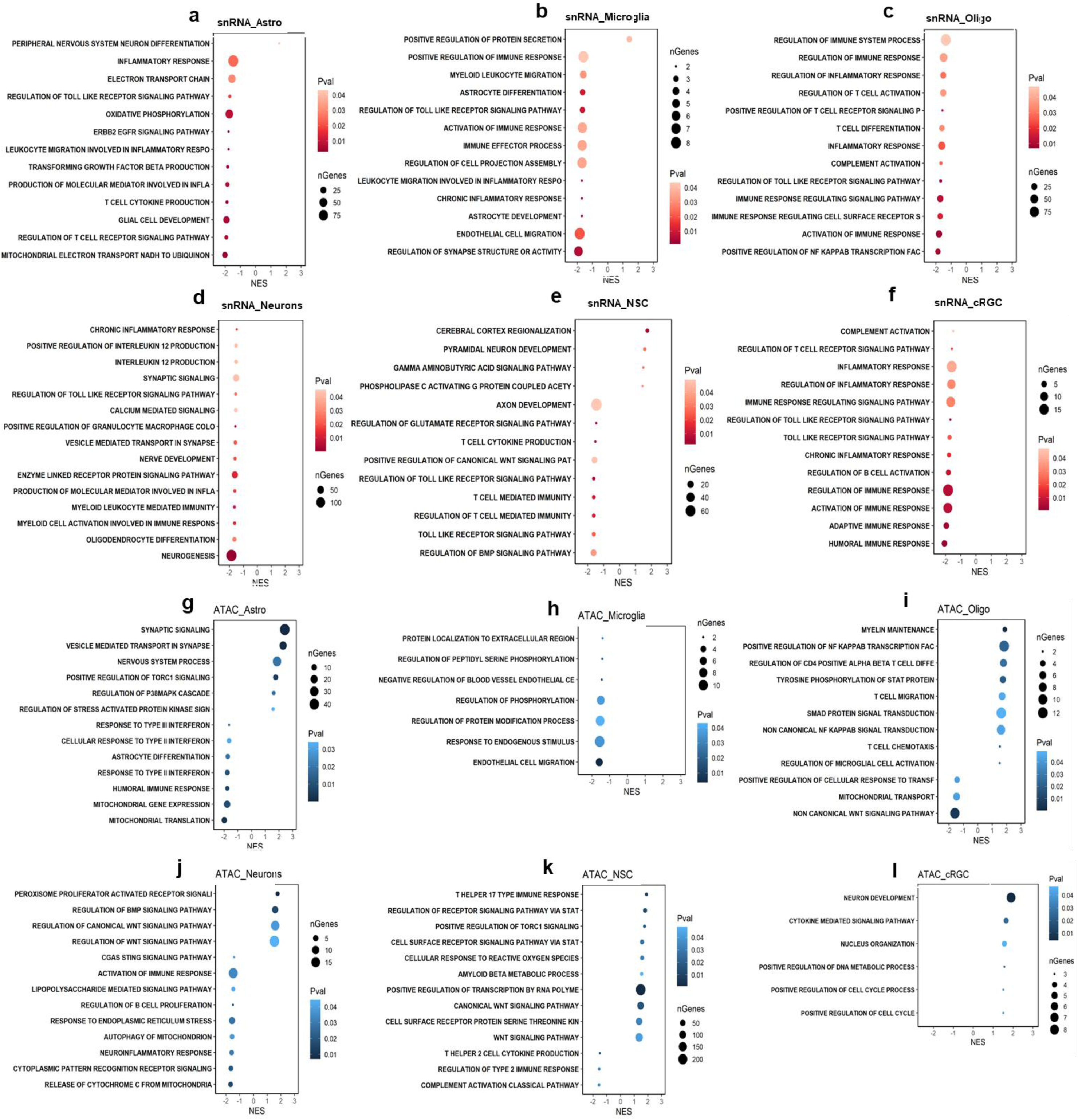
Identifying key regulators of HIV-1-infected cerebral organoids through snRNA seq and ATAC seq analysis**. (A-F)** A dot plot visualizing differentially expressed gene levels of up and down-regulated inflammatory pathways, complement activation, TLR signaling pathway, and oxidative phosphorylation in different cell types of snRNA sequencing and **(G-I)** ATAC sequencing data.

**Supplementary figure S2:**
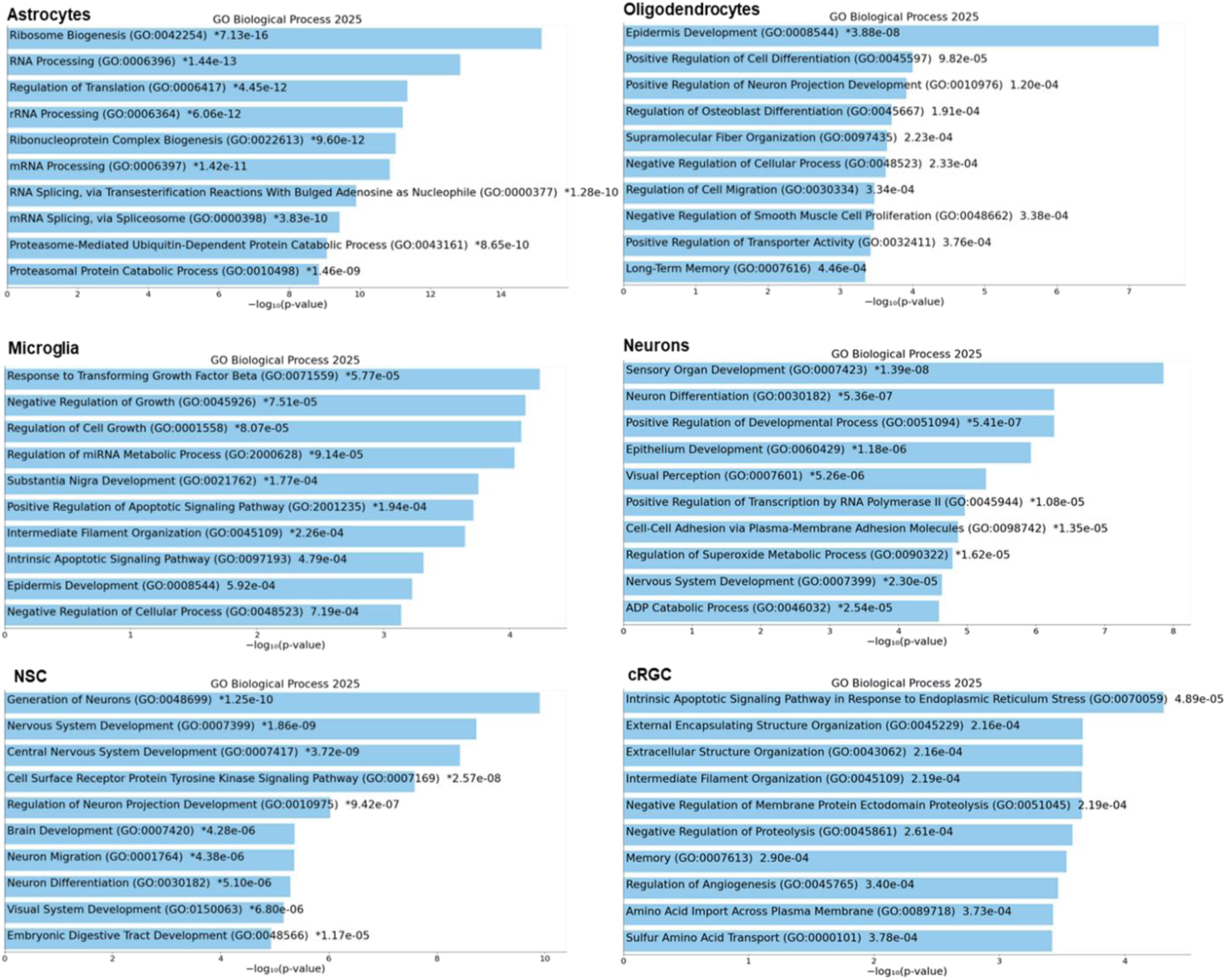
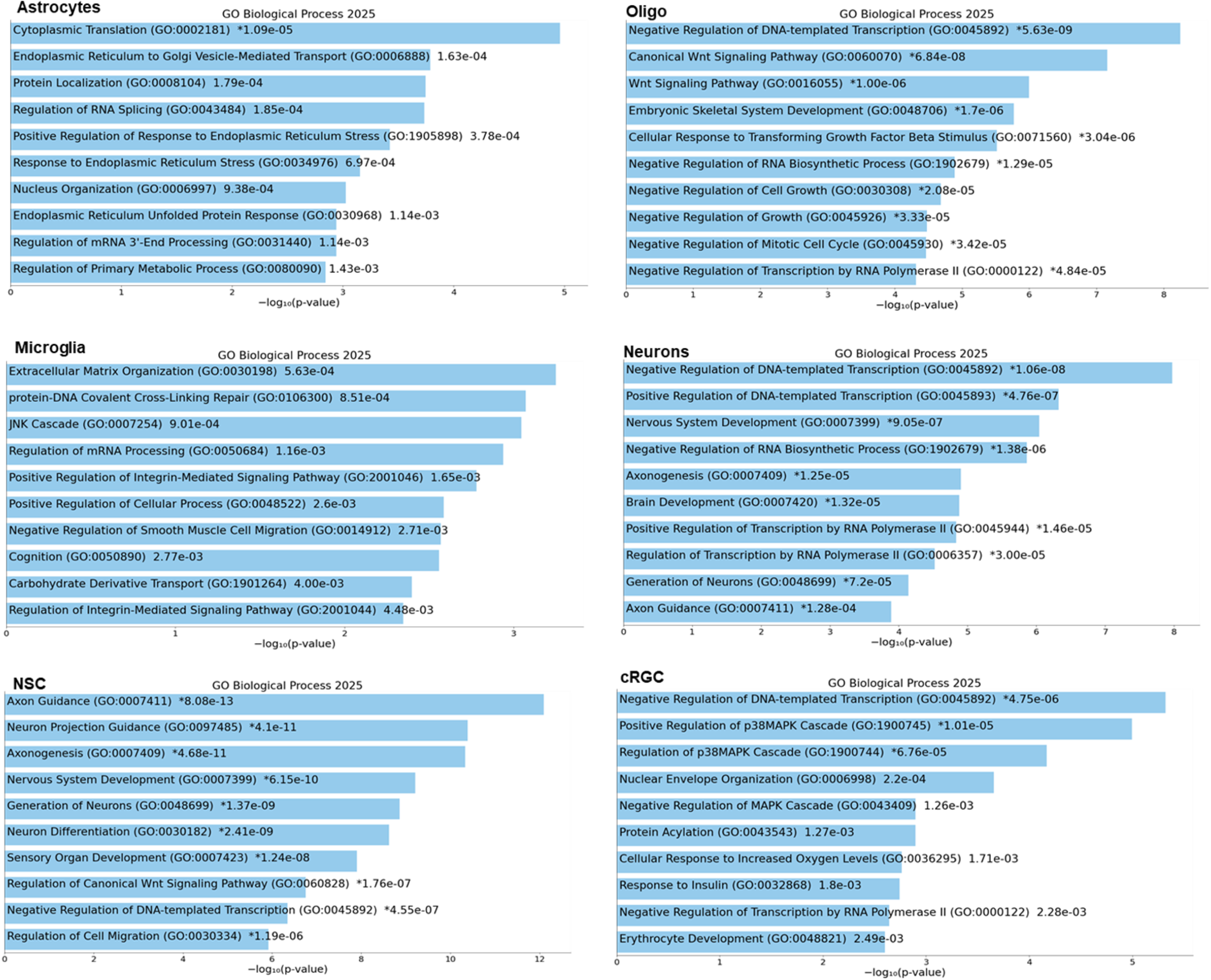
Functional enrichment analysis of differentially expressed genes. (**A)** Top 10 GO terms for the DEGs based on snRNA seq (BPs). (**B)** Top 10 GO terms for DEGS based on ATAC scores from different cell type populations of HIV-1 vs Control. GO, Gene Ontology; BP, biological process; adjusted p-value.

**Supplementary figure S3:**
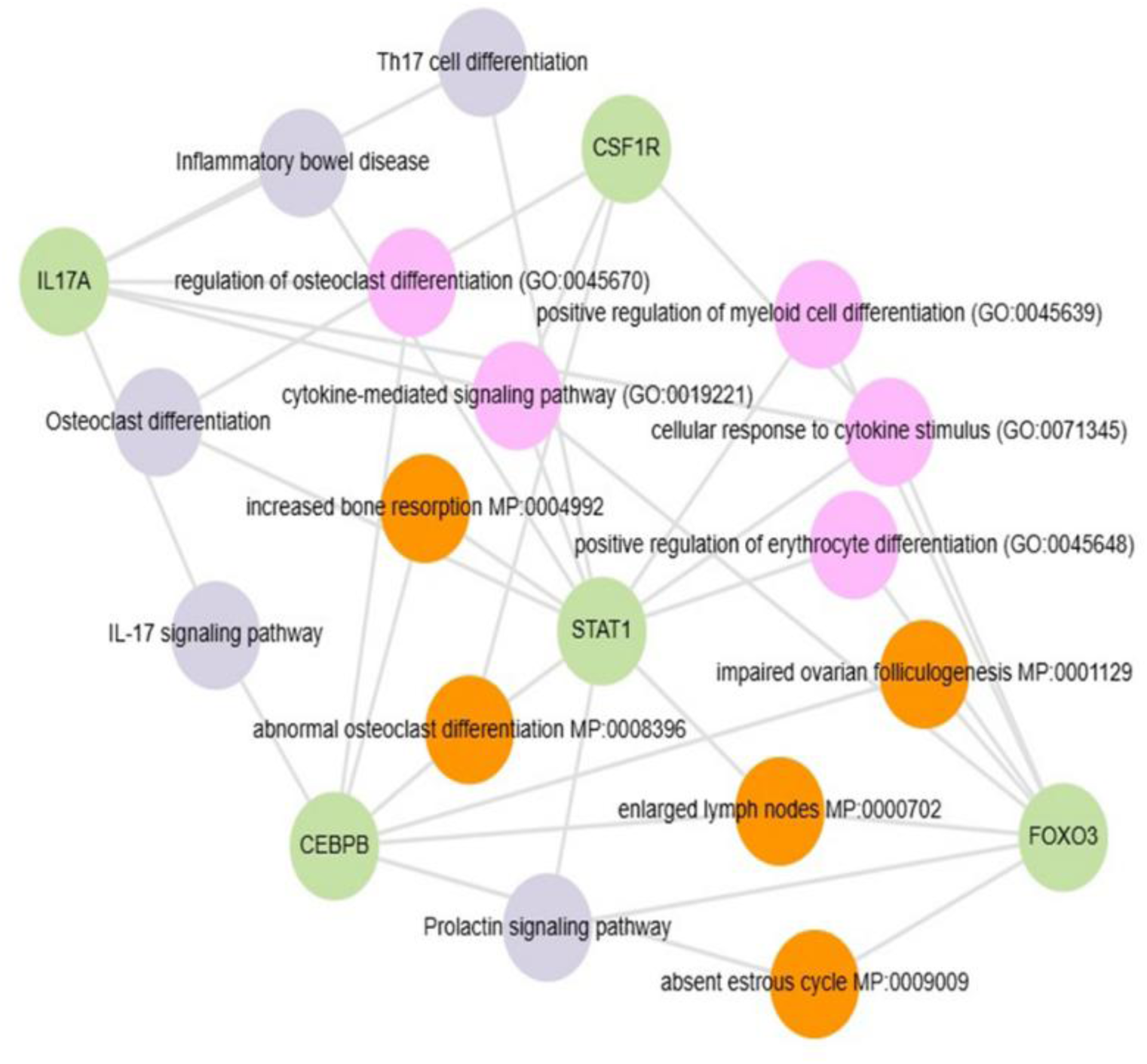
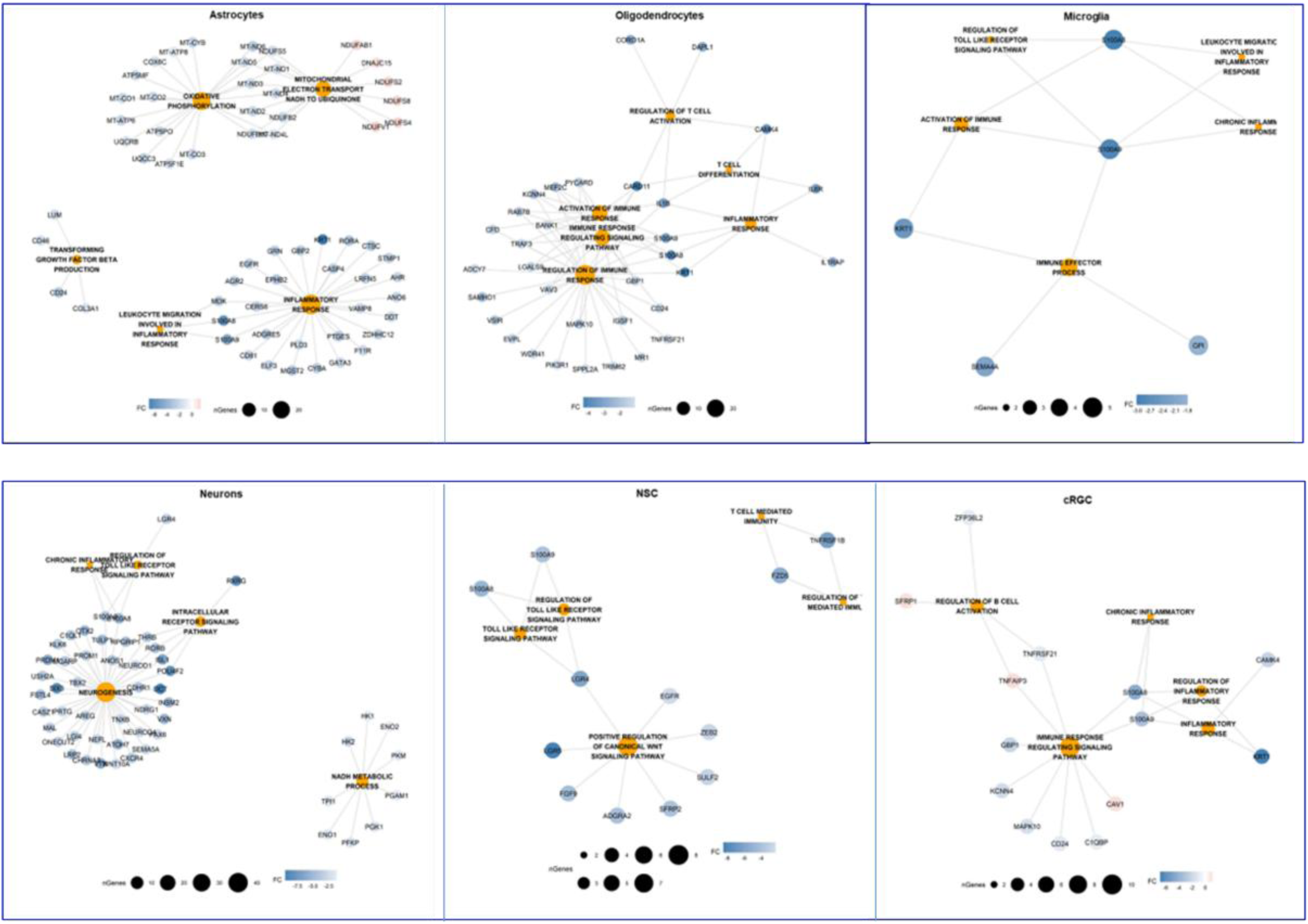
(**A)** Enrichr analysis was performed to identify the most significantly enriched GO terms. The enriched biological process terms from HIV-1-infected hCOs, show interaction between CEBPB, IL-17A, STAT1, and CSF1R signaling pathways. B. Visualization of top enriched GO pathways for the upregulated DEGs across various cell types.

**Supplementary figure S4:**
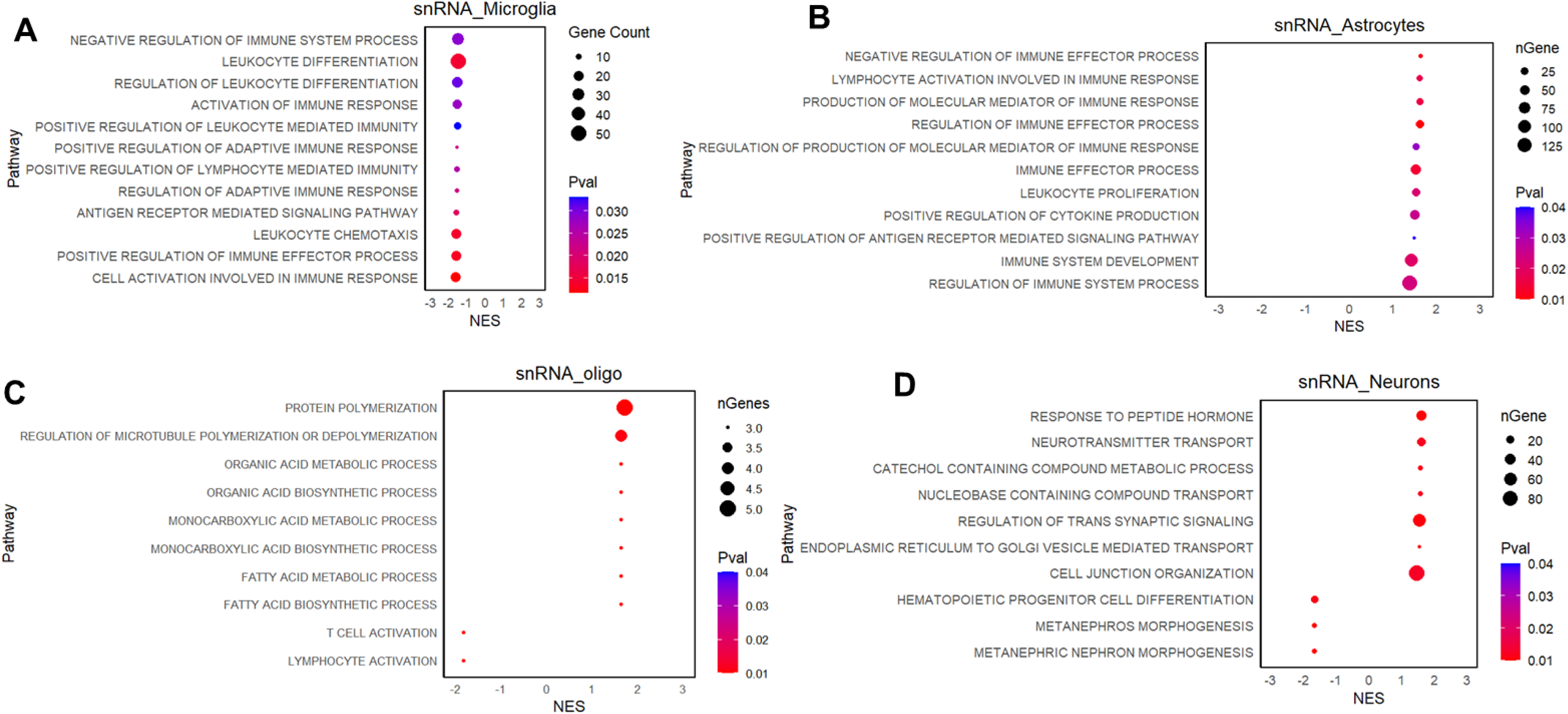
Functional enrichment analysis of biological processes in the macaque brain. (**A-D)** Cell type specific functional enrichment analysis of biological processes using upregulated immune-related genes in the macaque brain, GO Biological Process pathways were analyzed using fgsea with ranked gene lists (log_2_FC ≥ 1, p < 0.05, ≥5% expression). Significant pathways (nominal p < 0.05) were visualized as dot plots, where the diameter of the dot corresponds to gene counts.

**Supplementary figure S5.**
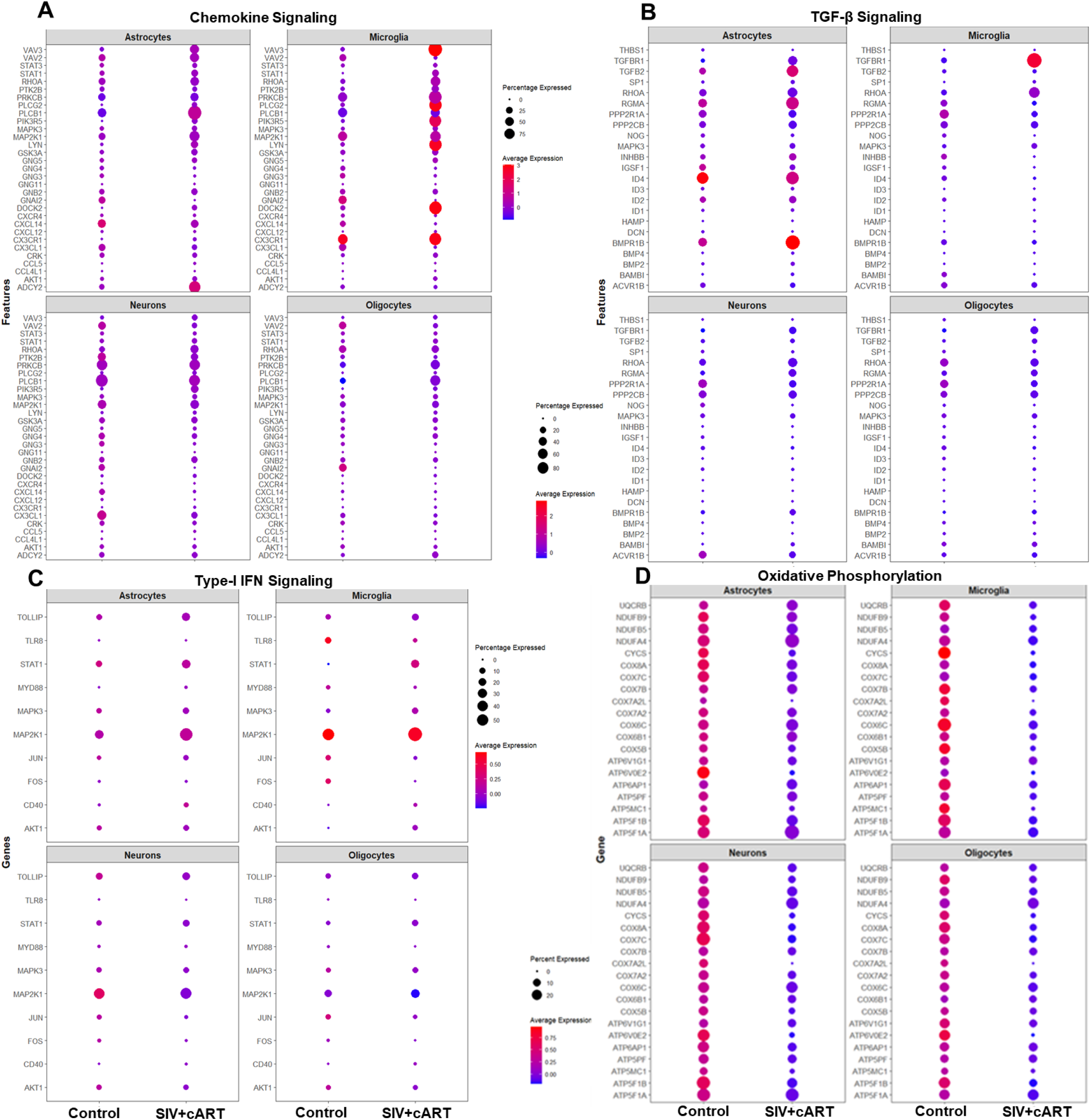
(A-D) Selected genes of specific pathways, including chemokine signaling, TGF-β signaling, type I interferon signaling, and oxidative phosphorylation, were presented as dot plots, where the diameter of the dots indicates the percentage of cells in which specific genes were expressed. NES: Normalized enrichment scores.

**Supplementary figure S6:**
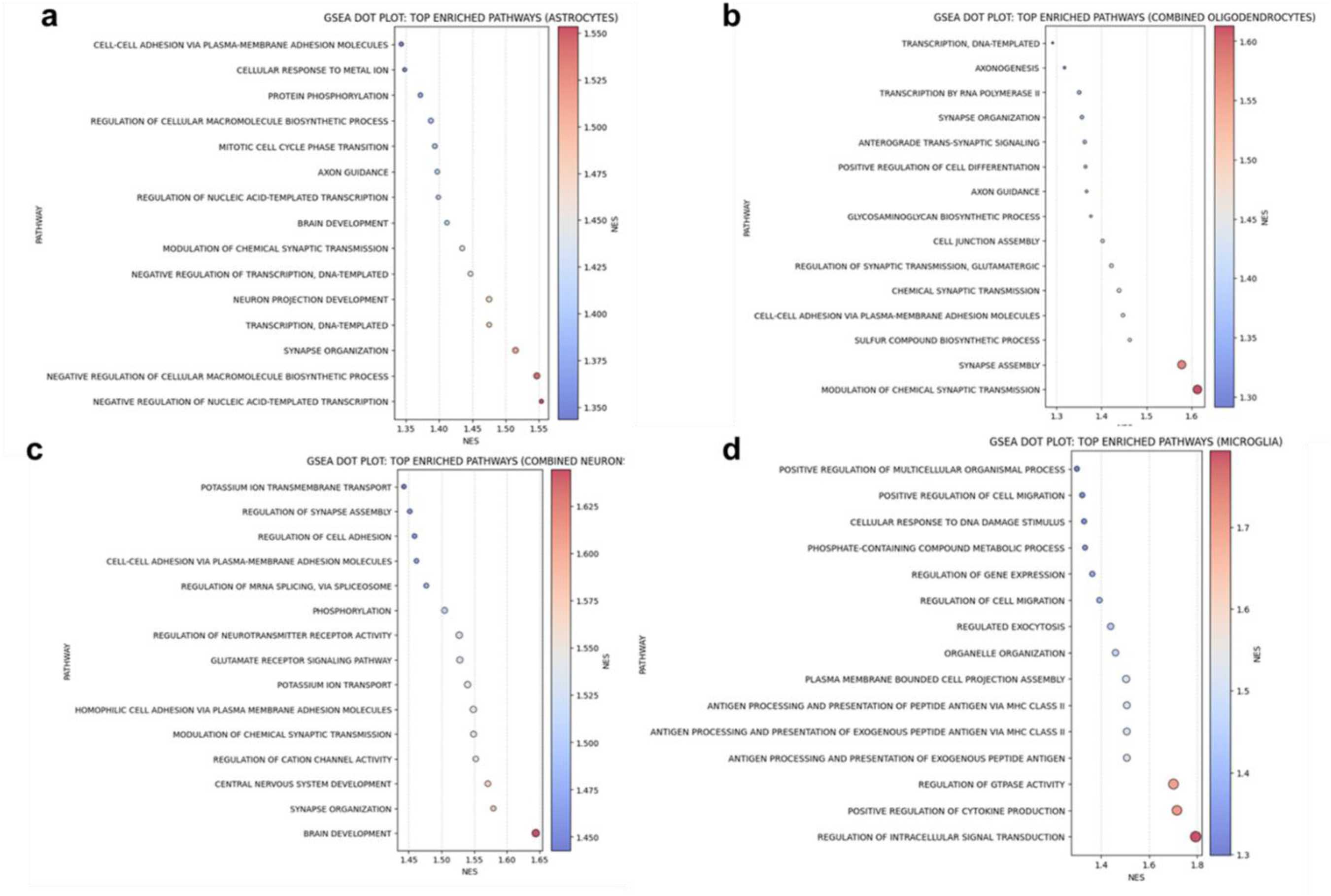
Gene Set Enrichment Analysis (GSEA) of the top enriched pathways in astrocytes (**A),** oligodendrocytes **(B),** neurons (**C),** and microglia (**D)** from control and SIV+ART treated RMs. The size of the dots indicates the number of genes involved in each pathway. NES: Normalized enrichment scores.

**Supplementary figure S7:**
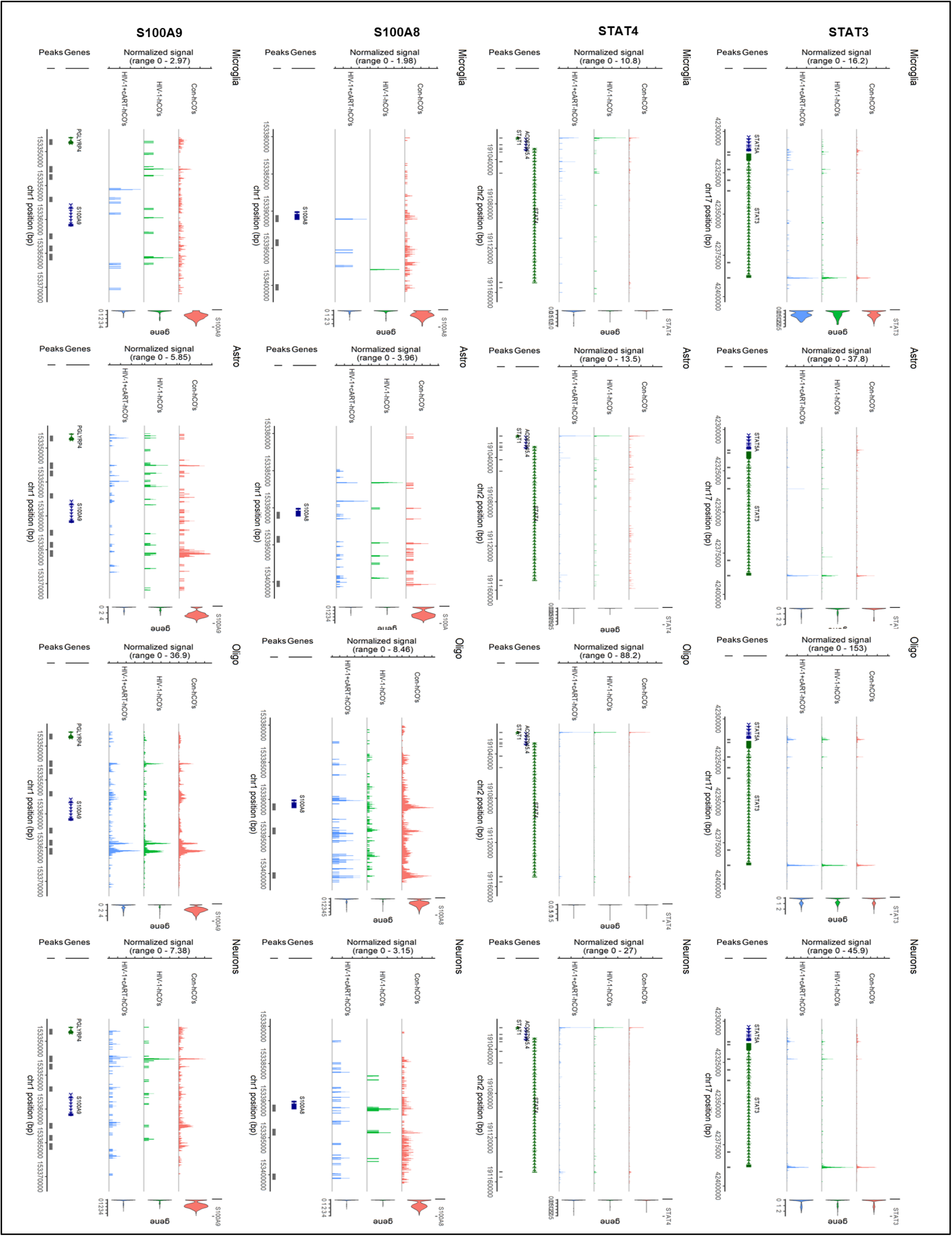
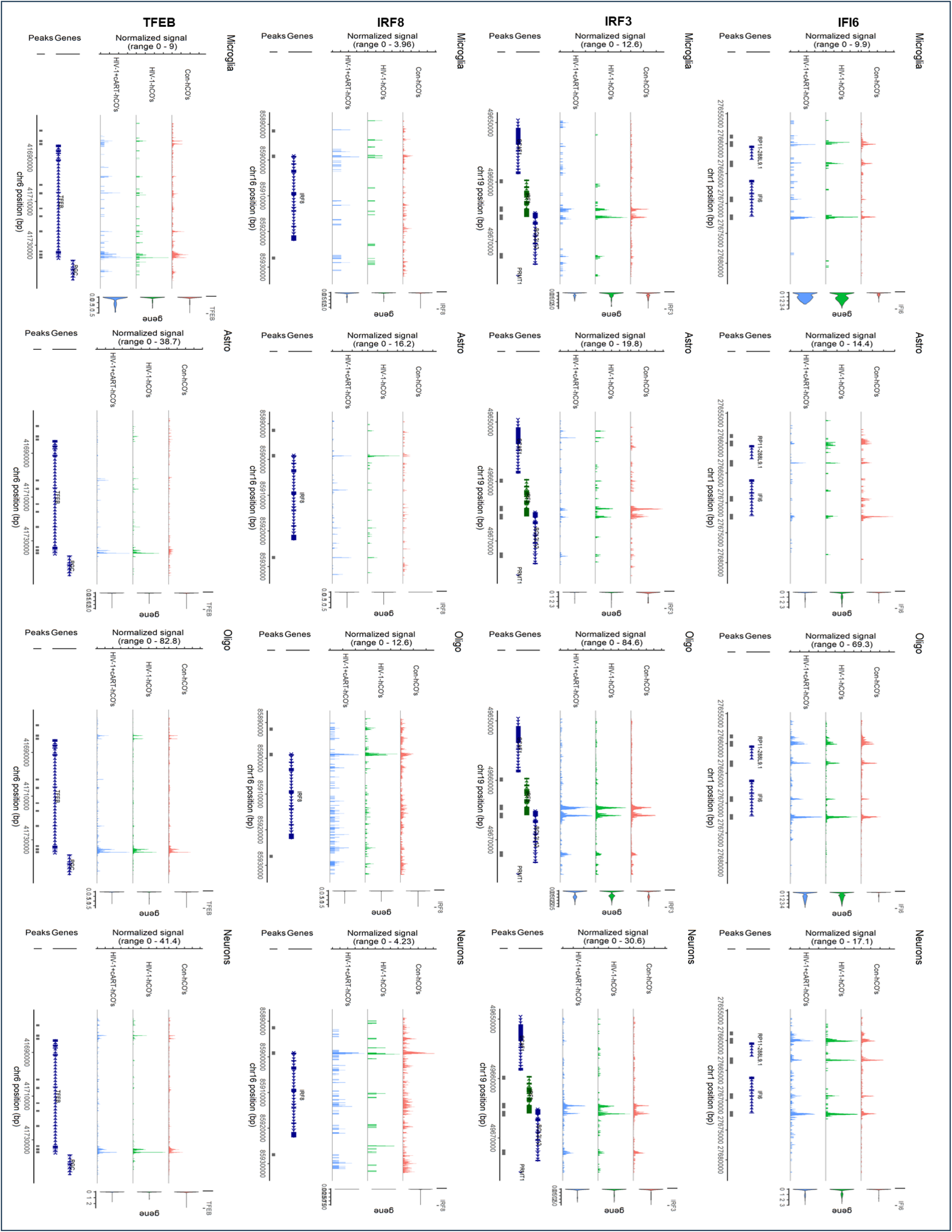
Representative sequencing tracks for the IFI6, IRF3, IRF8, TFEB, STAT3, STAT4, S100A8, and S100A9 gene loci show distinct ATAC-seq peaks at the promoter and the known enhancer in hCOs. The ATAC-seq data have been normalized to take sequencing depth into account, and the scale on the y-axis was chosen for optimal visualization of peaks for each cell type.

**Supplementary figure S8:**
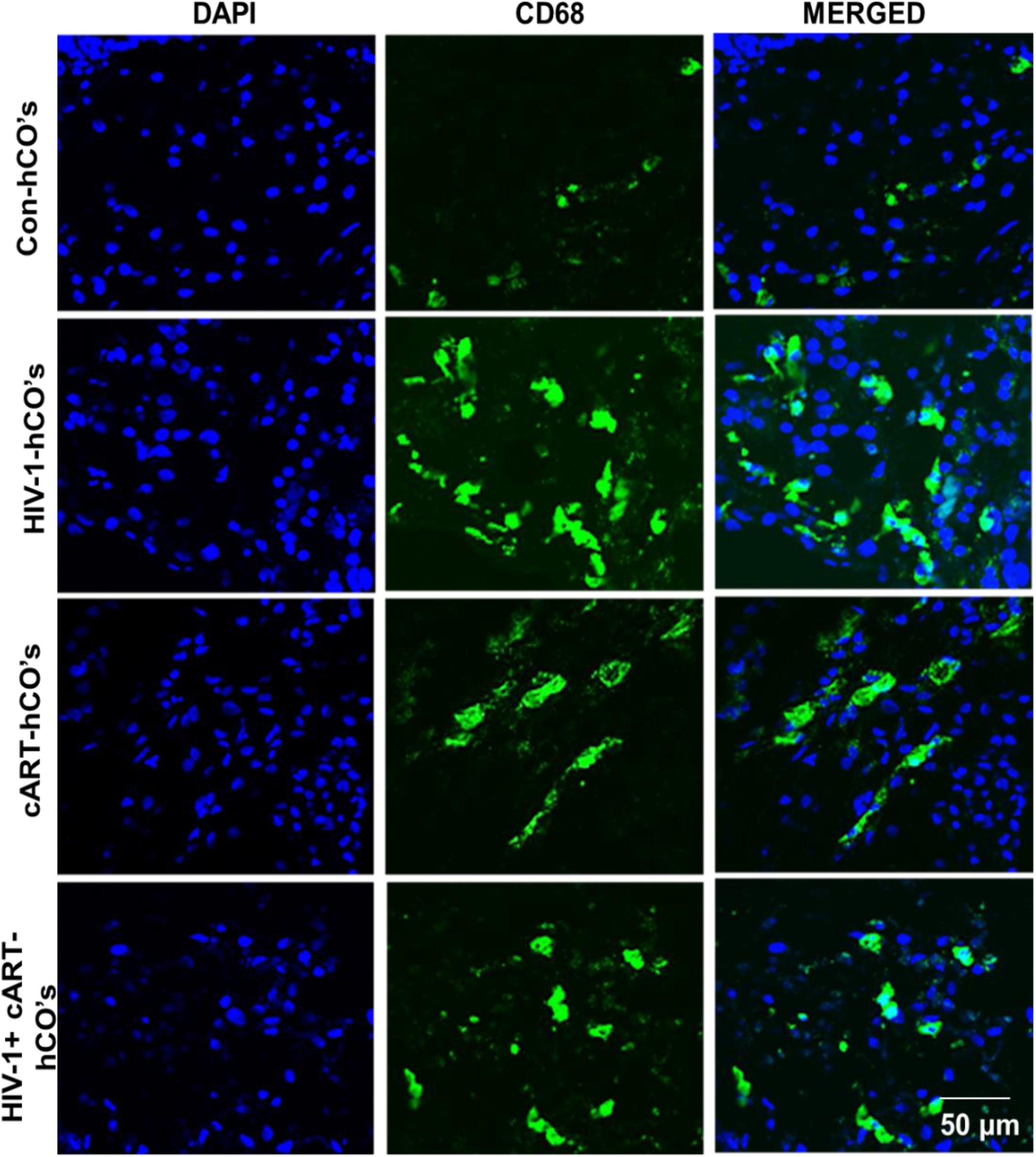
Immunostaining of microglia/macrophages activation marker CD68 is expressed in HIV-1-infected hCOs compared to control and cART treatment. Scale bar: 50μm.

**Supplementary figure S9:**
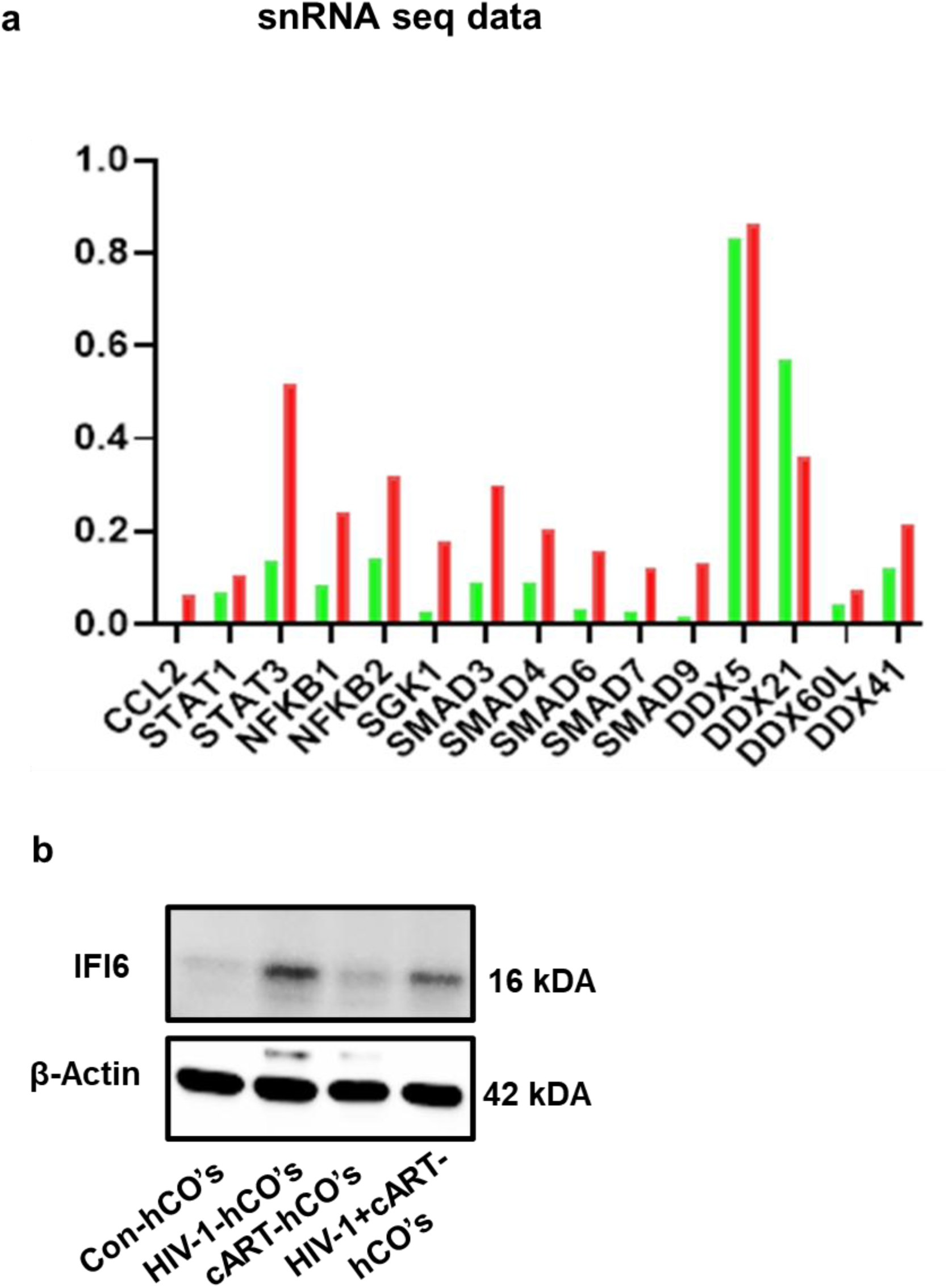
(**A)** RNA-seq analysis showing the expression levels of CCL2, STAT1, STAT3, NFKB1, NFKB2, SGK1, SMAD, and DDX genes in HIV-1-infected hCOs vs control hCOs. The graph represents log2 fold-change (FC) ratios, highlighting transcriptional alterations induced by HIV-1 infection. (**B)** Western blot analysis of IFI6 expression in HIV-infected hCOs compared to control and cART treatment. β-Actin served as an internal control.

**Supplementary figure S10:**
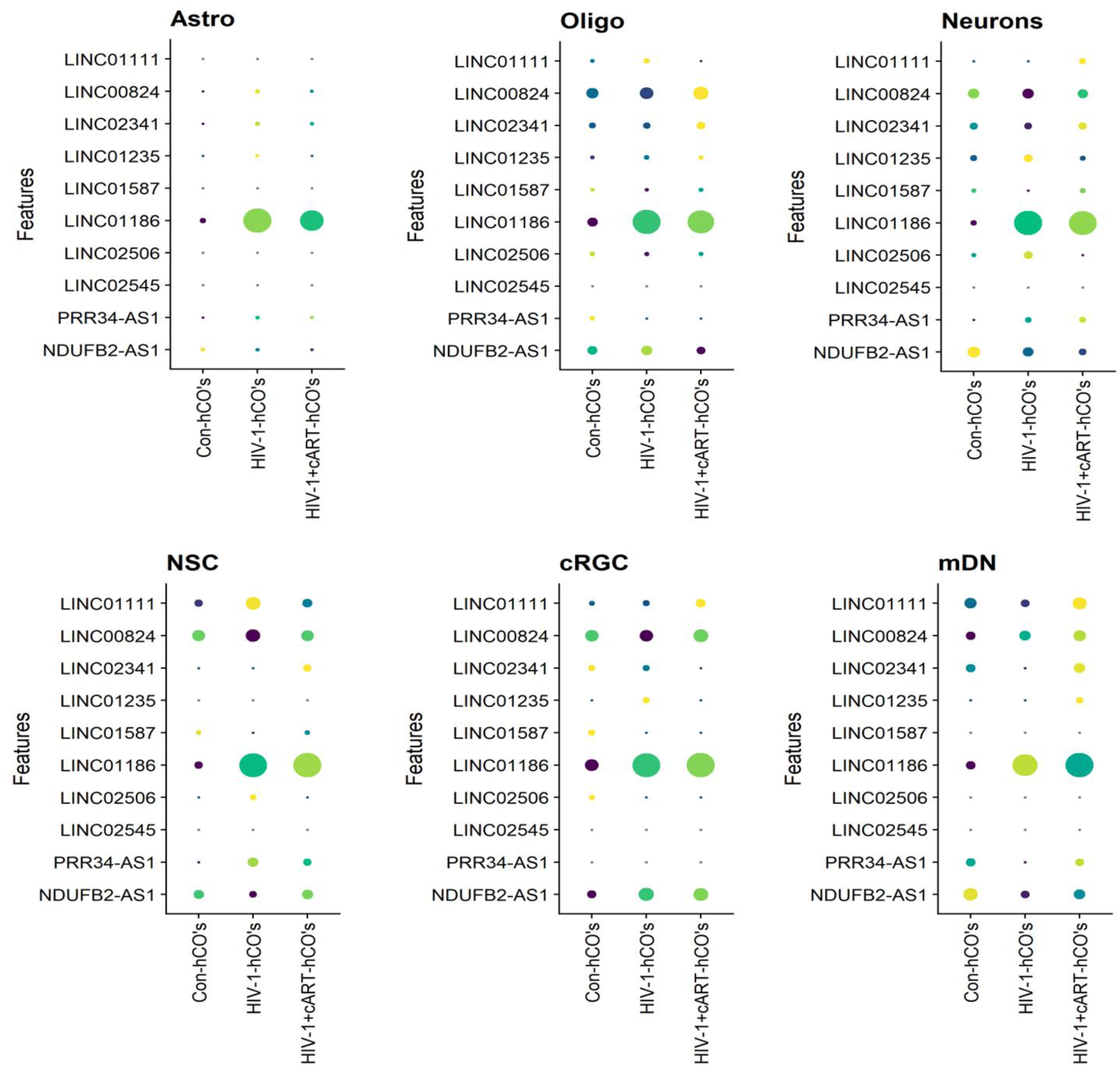
GO enrichment analysis of the astrocytes, neurons, mDN, NSCs, Oligodendrocytes, and cRGC linc-RNA nearby genes. Nearby genes were collected within 100 kb up- or downstream of lincRNAs related to TGF-β signaling and oxidative phosphorylation between each group.

## References

1. Global Task Team on, H.I.V.T.W.G.E.a.m.u.o., Global HIV targets: a roadmap to 2030 and beyond. Lancet, 2025.

2. Saylor, D., et al., HIV-associated neurocognitive disorder--pathogenesis and prospects for treatment. Nat Rev Neurol, 2016. 12(4): p. 234–48.

3. Ziffra, R.S., et al., Single-cell epigenomics reveals mechanisms of human cortical development. Nature, 2021. 598(7879): p. 205–213.

4. Sato, Y., T. Asahi, and K. Kataoka, Integrative single-cell RNA-seq analysis of vascularized cerebral organoids. BMC Biol, 2023. 21(1): p. 245.

5. Lancaster, M.A. and J.A. Knoblich, Generation of cerebral organoids from human pluripotent stem cells. Nat Protoc, 2014. 9(10): p. 2329–40.

6. Dos Reis, R.S., S. Sant, and V. Ayyavoo, Three-Dimensional Human Brain Organoids to Model HIV-1 Neuropathogenesis. Methods Mol Biol, 2023. 2610: p. 167–178.

7. Ormel, P.R., et al., Microglia innately develop within cerebral organoids. Nat Commun, 2018. 9(1): p. 4167.

8. Kong, W., et al., Neuroinflammation generated by HIV-infected microglia promotes dysfunction and death of neurons in human brain organoids. PNAS Nexus, 2024. 3(5): p. pgae179.

9. Kwon, H.S. and S.H. Koh, Neuroinflammation in neurodegenerative disorders: the roles of microglia and astrocytes. Transl Neurodegener, 2020. 9(1): p. 42.

10. Simpson-Abelson, M.R., et al., CCAAT/Enhancer-binding protein β promotes pathogenesis of EAE. Cytokine, 2017. 92: p. 24–32.

11. Schwartz, C., et al., Functional interactions between C/EBP, Sp1, and COUP-TF regulate human immunodeficiency virus type 1 gene transcription in human brain cells. J Virol, 2000. 74(1): p. 65–73.

12. Akira, S., et al., A nuclear factor for IL-6 expression (NF-IL6) is a member of a C/EBP family. The EMBO Journal, 1990. 9(6): p. 1897–1906.

13. Ren, Y., W. Guo, and B. Qiao, Abnormal expression of CEBPB promotes the progression of renal cell carcinoma through regulating the generation of IL-6. Heliyon, 2023. 9(10): p. e20175.

14. Abraham, S., et al., Cooperative interaction of C/EBP beta and Tat modulates MCP-1 gene transcription in astrocytes. J Neuroimmunol, 2005. 160(1-2): p. 219–27.

15. Ko, C.Y., et al., Glycogen synthase kinase-3beta-mediated CCAAT/enhancer-binding protein delta phosphorylation in astrocytes promotes migration and activation of microglia/macrophages. Neurobiol Aging, 2014. 35(1): p. 24–34.

16. Siebel, C. and U. Lendahl, Notch Signaling in Development, Tissue Homeostasis, and Disease. Physiol Rev, 2017. 97(4): p. 1235–1294.

17. Koutelou, E., et al., Neuralized-like 1 (Neurl1) targeted to the plasma membrane by N-myristoylation regulates the Notch ligand Jagged1. J Biol Chem, 2008. 283(7): p. 3846–53.

18. Keerthivasan, S., et al., Notch signaling regulates mouse and human Th17 differentiation. J Immunol, 2011. 187(2): p. 692–701.

19. Dash, P.K., et al., Loss of neuronal integrity during progressive HIV-1 infection of humanized mice. J Neurosci, 2011. 31(9): p. 3148–57.

20. Fleck, J.S., et al., Inferring and perturbing cell fate regulomes in human brain organoids. Nature, 2023. 621(7978): p. 365–372.

21. Schmidtmayerova, H., et al., Human immunodeficiency virus type 1 T-lymphotropic strains enter macrophages via a CD4- and CXCR4-mediated pathway: replication is restricted at a postentry level. J Virol, 1998. 72(6): p. 4633–42.

22. Acharya, A., et al., Chronic morphine administration differentially modulates viral reservoirs in SIVmac251 infected rhesus macaque model. J Virol, 2021. 95(5).

23. Fox, H.S., et al., Morphine suppresses peripheral responses and transforms brain myeloid gene expression to favor neuropathogenesis in SIV infection. Front Immunol, 2022. 13: p. 1012884.

24. Hoffman, G.E., et al., Efficient differential expression analysis of large-scale single cell transcriptomics data using dreamlet. bioRxiv, 2024.

25. Honeycutt, J.B., et al., T cells establish and maintain CNS viral infection in HIV-infected humanized mice. J Clin Invest, 2018. 128(7): p. 2862–2876.

26. Doke, M., et al., HIV-1 Tat and cocaine impact astrocytic energy reservoirs and epigenetic regulation by influencing the LINC01133-hsa-miR-4726-5p-NDUFA9 axis. Mol Ther Nucleic Acids, 2022. 29: p. 243–258.

27. Pellegrini, L., et al., Human CNS barrier-forming organoids with cerebrospinal fluid production. Science, 2020. 369(6500).

28. Giandomenico, S.L., M. Sutcliffe, and M.A. Lancaster, Generation and long-term culture of advanced cerebral organoids for studying later stages of neural development. Nat Protoc, 2021. 16(2): p. 579–602.

29. Min, A.K., et al., HIV-1 infection of genetically engineered iPSC-derived central nervous system-engrafted microglia in a humanized mouse model. J Virol, 2023. 97(12): p. e0159523.

30. Walsh, J.G., et al., Rapid inflammasome activation in microglia contributes to brain disease in HIV/AIDS. Retrovirology, 2014. 11: p. 35.

31. Pan, X., et al., Restrictions to HIV-1 replication in resting CD4+ T lymphocytes. Cell Res, 2013. 23(7): p. 876–85.

32. Rao, V.R., et al., Clade C HIV-1 isolates circulating in Southern Africa exhibit a greater frequency of dicysteine motif-containing Tat variants than those in Southeast Asia and cause increased neurovirulence. Retrovirology, 2013. 10: p. 61.

33. Zhang, K., et al., A single-cell atlas of chromatin accessibility in the human genome. Cell, 2021. 184(24): p. 5985–6001 e19.

34. Phan, B.N., et al., Single nuclei transcriptomics in human and non-human primate striatum in opioid use disorder. Nat Commun, 2024. 15(1): p. 878.

35. Ma, Y., et al., Integrative differential expression and gene set enrichment analysis using summary statistics for scRNA-seq studies. Nat Commun, 2020. 11(1): p. 1585.

36. Oguariri, R.M., et al., Short Communication: S100A8 and S100A9, Biomarkers of SARS-CoV-2 Infection and Other Diseases, Suppress HIV Replication in Primary Macrophages. AIDS Res Hum Retroviruses, 2022. 38(5): p. 401–405.

37. Maarifi, G., et al., Alarmin S100A9 restricts retroviral infection by limiting reverse transcription in human dendritic cells. EMBO J, 2021. 40(16): p. e106540.

38. Sartorius, R., et al., Exploiting viral sensing mediated by Toll-like receptors to design innovative vaccines. NPJ Vaccines, 2021. 6(1): p. 127.

39. Suh, H.S., et al., TLR3 and TLR4 are innate antiviral immune receptors in human microglia: role of IRF3 in modulating antiviral and inflammatory response in the CNS. Virology, 2009. 392(2): p. 246–59.

40. Sacktor, N., et al., Prevalence of HIV-associated neurocognitive disorders in the Multicenter AIDS Cohort Study. Neurology, 2016. 86(4): p. 334–40.

41. Clifford, D.B. and B.M. Ances, HIV-associated neurocognitive disorder. Lancet Infect Dis, 2013. 13(11): p. 976–86.

42. Sun, L., et al., Cyclic GMP-AMP synthase is a cytosolic DNA sensor that activates the type I interferon pathway. Science, 2013. 339(6121): p. 786–91.

43. Zheng, M.Y. and L.Z. Luo, The Role of IL-17A in Mediating Inflammatory Responses and Progression of Neurodegenerative Diseases. Int J Mol Sci, 2025. 26(6).

44. Guo, C., et al., Oxidative stress, mitochondrial damage and neurodegenerative diseases. Neural Regen Res, 2013. 8(21): p. 2003–14.

45. Chapuis, J., et al., Increased expression of BIN1 mediates Alzheimer genetic risk by modulating tau pathology. Mol Psychiatry, 2013. 18(11): p. 1225–34.

46. Anthony, I.C., et al., Accelerated Tau deposition in the brains of individuals infected with human immunodeficiency virus-1 before and after the advent of highly active anti-retroviral therapy. Acta Neuropathol, 2006. 111(6): p. 529–38.

47. Eugenin, E.A., et al., Human immunodeficiency virus infection of human astrocytes disrupts blood-brain barrier integrity by a gap junction-dependent mechanism. J Neurosci, 2011. 31(26): p. 9456–65.

48. Xu, X., et al., T cell-mediated SIV dissemination into the CNS: a single-cell transcriptomic analysis. J Neuroinflammation, 2025. 22(1): p. 226.

49. Ash, M.K., L. Al-Harthi, and J.R. Schneider, HIV in the Brain: Identifying Viral Reservoirs and Addressing the Challenges of an HIV Cure. Vaccines (Basel), 2021. 9(8).

50. Sui, Y., et al., Neuronal apoptosis is mediated by CXCL10 overexpression in simian human immunodeficiency virus encephalitis. Am J Pathol, 2004. 164(5): p. 1557–66.

51. Lawrence, D.M., et al., Astrocyte differentiation selectively upregulates CCL2/monocyte chemoattractant protein-1 in cultured human brain-derived progenitor cells. Glia, 2006. 53(1): p. 81–91.

52. Ramji, D.P. and P. Foka, CCAAT/enhancer-binding proteins: structure, function and regulation. Biochem J, 2002. 365(Pt 3): p. 561–75.

53. Maitra, A., et al., Distinct functional motifs within the IL-17 receptor regulate signal transduction and target gene expression. Proc Natl Acad Sci U S A, 2007. 104(18): p. 7506–11.

54. Nitkiewicz, J., et al., HIV induces expression of complement component C3 in astrocytes by NF-kappaB-dependent activation of interleukin-6 synthesis. J Neuroinflammation, 2017. 14(1): p. 23.

55. Liu, Z., et al., IL-17A exacerbates neuroinflammation and neurodegeneration by activating microglia in rodent models of Parkinson’s disease. Brain Behav Immun, 2019. 81: p. 630–645.

56. Irons, D.L., et al., Overexpression and activation of colony-stimulating factor 1 receptor in the SIV/macaque model of HIV infection and neuroHIV. Brain Pathol, 2019. 29(6): p. 826–836.

57. Bo, L. and X. Bo, Colony stimulating factor 1: friend or foe of neurons? Neural Regen Res, 2022. 17(4): p. 773–774.

58. Guo, S., H. Wang, and Y. Yin, Microglia Polarization From M1 to M2 in Neurodegenerative Diseases. Front Aging Neurosci, 2022. 14: p. 815347.

59. Taal, K., et al., Neuralized family member NEURL1 is a ubiquitin ligase for the cGMP-specific phosphodiesterase 9A. Scientific Reports, 2019. 9(1): p. 7104.

60. Wang, W., et al., Mitochondria dysfunction in the pathogenesis of Alzheimer’s disease: recent advances. Molecular Neurodegeneration, 2020. 15(1): p. 30.

61. Henrich, M.T., et al., Mitochondrial dysfunction in Parkinson’s disease - a key disease hallmark with therapeutic potential. Mol Neurodegener, 2023. 18(1): p. 83.

62. Xia, Y., et al., C/EBPβ is a key transcription factor for APOE and preferentially mediates ApoE4 expression in Alzheimer’s disease. Mol Psychiatry, 2021. 26(10): p. 6002–6022.

63. Brown, L.A., et al., The role of tau protein in HIV-associated neurocognitive disorders. Mol Neurodegener, 2014. 9: p. 40.

64. Saha, O., et al., The Alzheimer’s disease risk gene BIN1 regulates activity-dependent gene expression in human-induced glutamatergic neurons. Mol Psychiatry, 2024. 29(9): p. 2634–2646.

65. Altmann, A., et al., Towards cascading genetic risk in Alzheimer’s disease. Brain, 2024. 147(8): p. 2680–2690.

66. Vallés-Saiz, L., J. Ávila, and F. Hernández, Lamivudine (3TC), a Nucleoside Reverse Transcriptase Inhibitor, Prevents the Neuropathological Alterations Present in Mutant Tau Transgenic Mice. Int J Mol Sci, 2023. 24(13).

