## Supplementary material for "STAT1-Mediated Regulation of IL-17A/CEBPB/NF-κB Axis in HIV-1 Infected Human Cerebral Organoids Reveals Therapeutic Targets for Neuroprotection": Excel file, Full blot, Tables: bioRxiv_Supplementary Material.docx

**Thangavel Samikkannu^1^***

^1^Department of Pharmaceutical Sciences, Irma Lerma Rangel College of Pharmacy, Texas A&M University Health Science Center, College Station, Texas -77843, USA.

^2^Department of Veterinary Integrative Biosciences, School of Veterinary Medicine and Biomedical Sciences, Texas A&M University, College Station, Texas -77843, USA.

^3^Department of Pharmacology and Experimental Neuroscience, College of Medicine, University of Nebraska Medical Center, Omaha, NE, USA

***Corresponding Author:**

Thangavel Samikkannu, Ph.D.

Associate Professor

Department of Pharmaceutical Sciences

Irma Lerma Rangel College of Pharmacy

Texas A&M University

College Station

Texas-77845, USA.

**Supplementary figure S1**

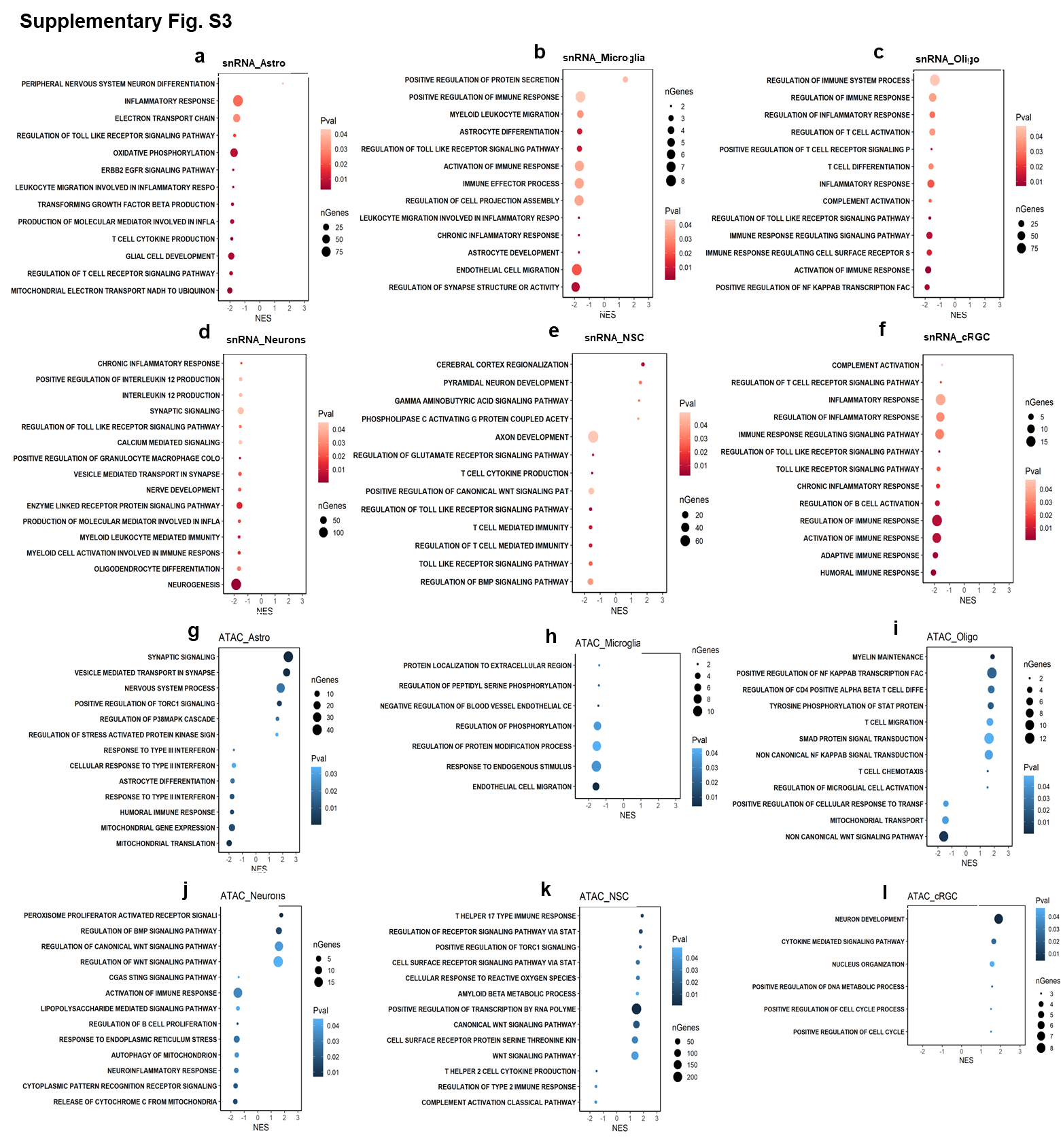

**Supplementary figure S2A**

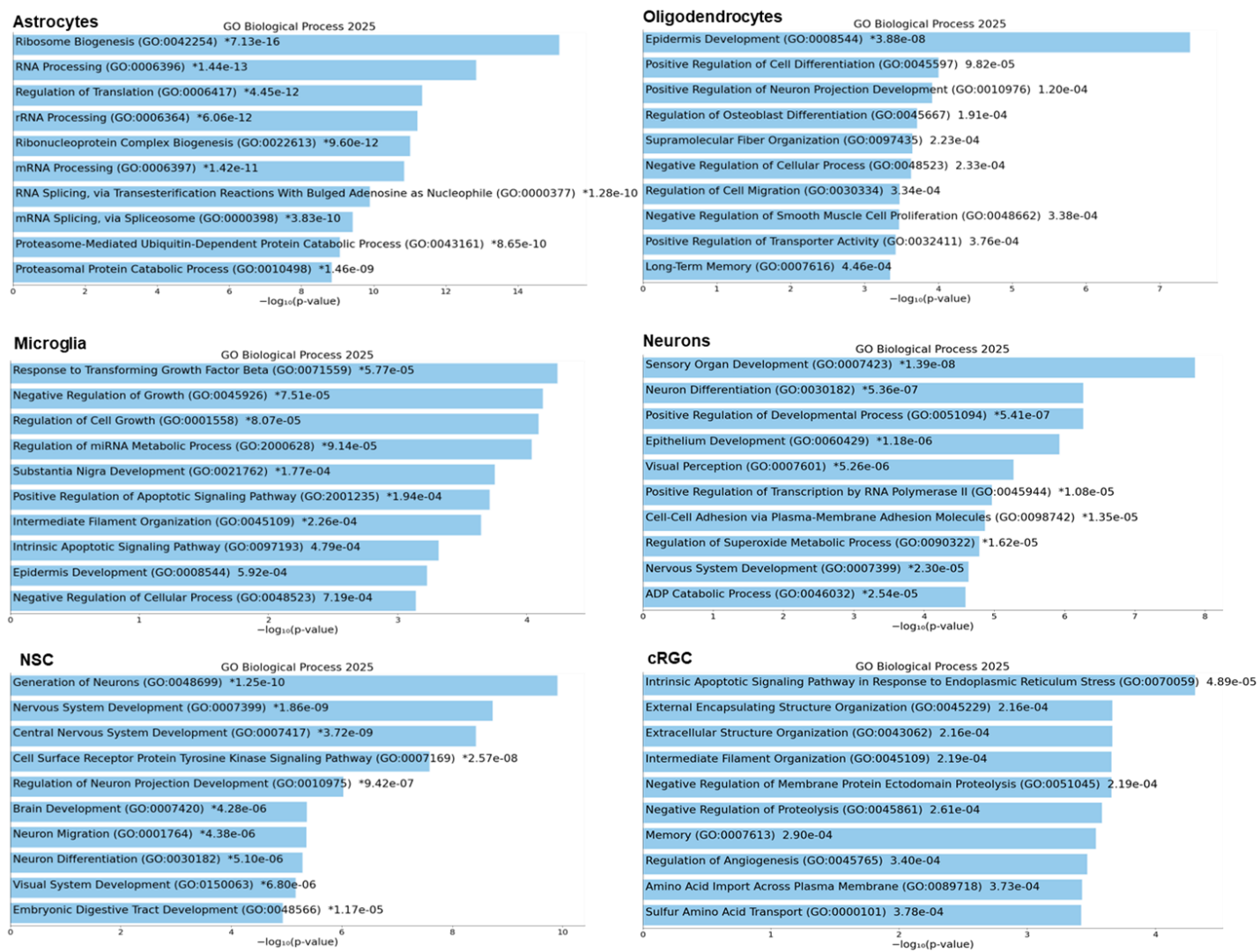

**Supplementary figure S2B**

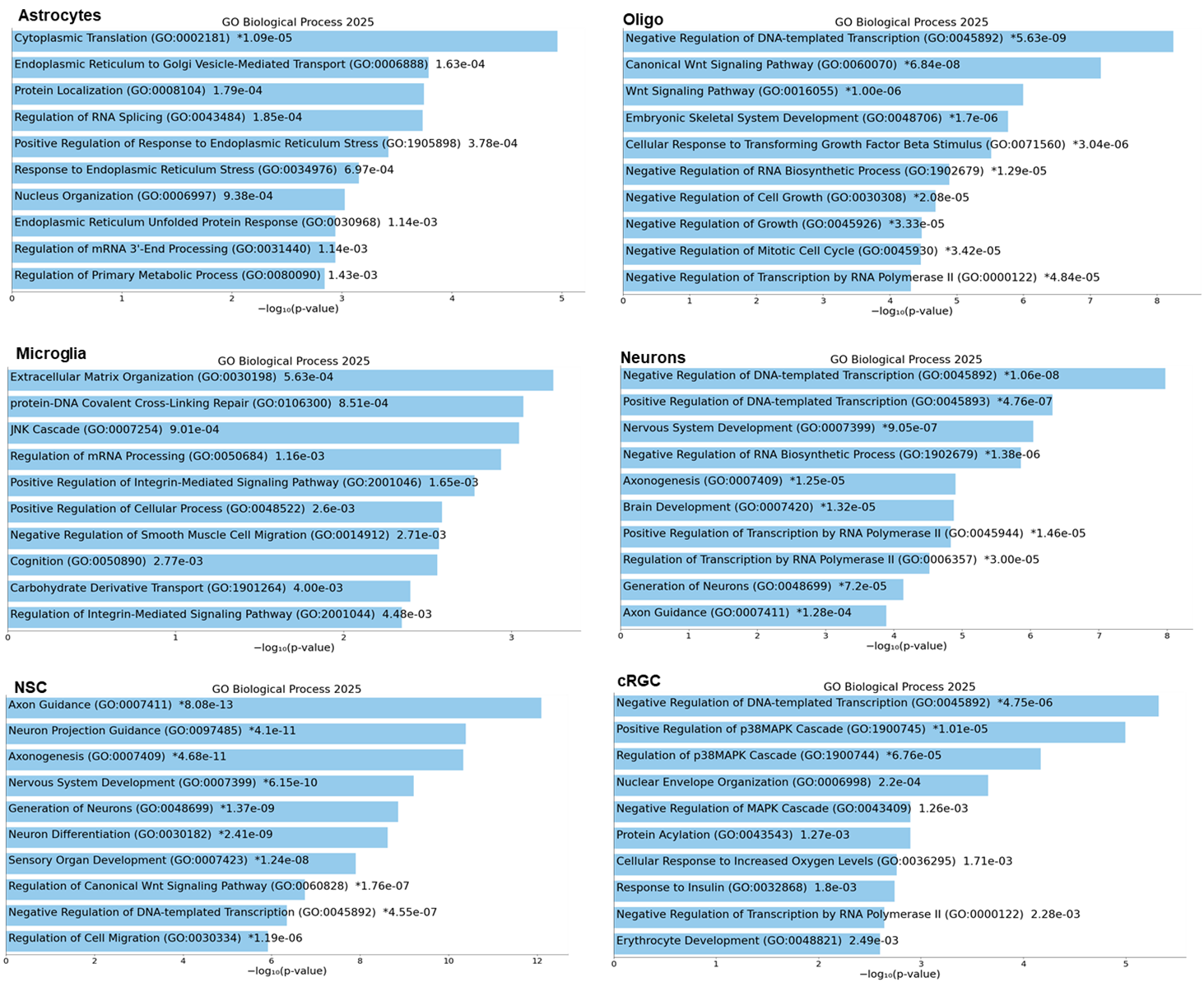

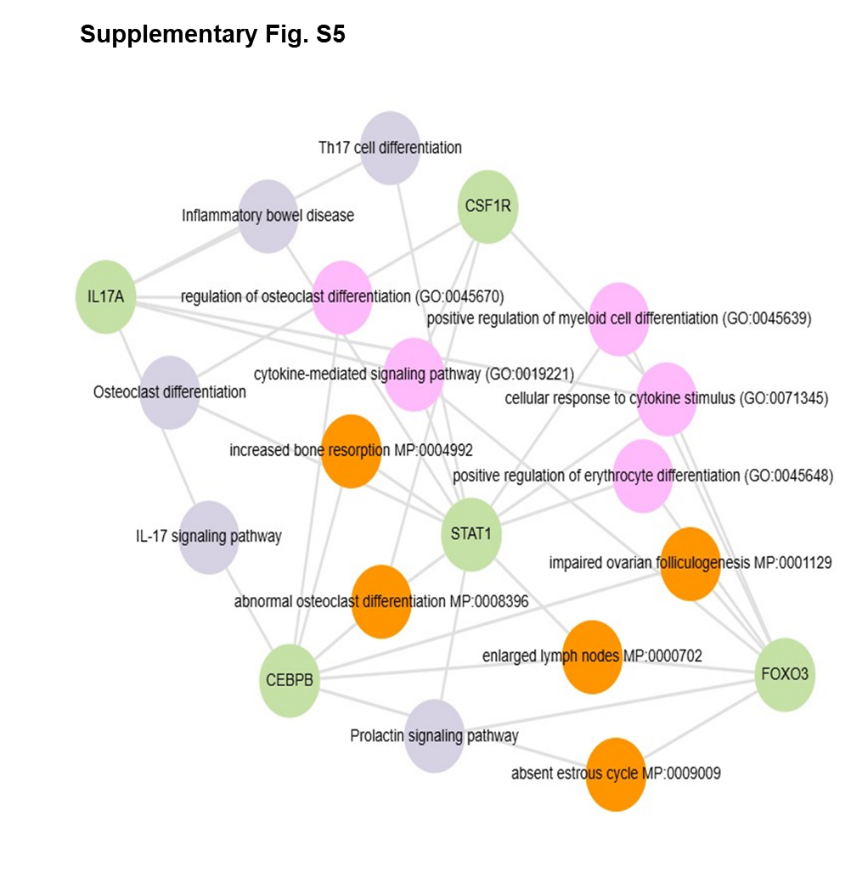
**Supplementary figure S3A**

**Supplementary figure S3B**

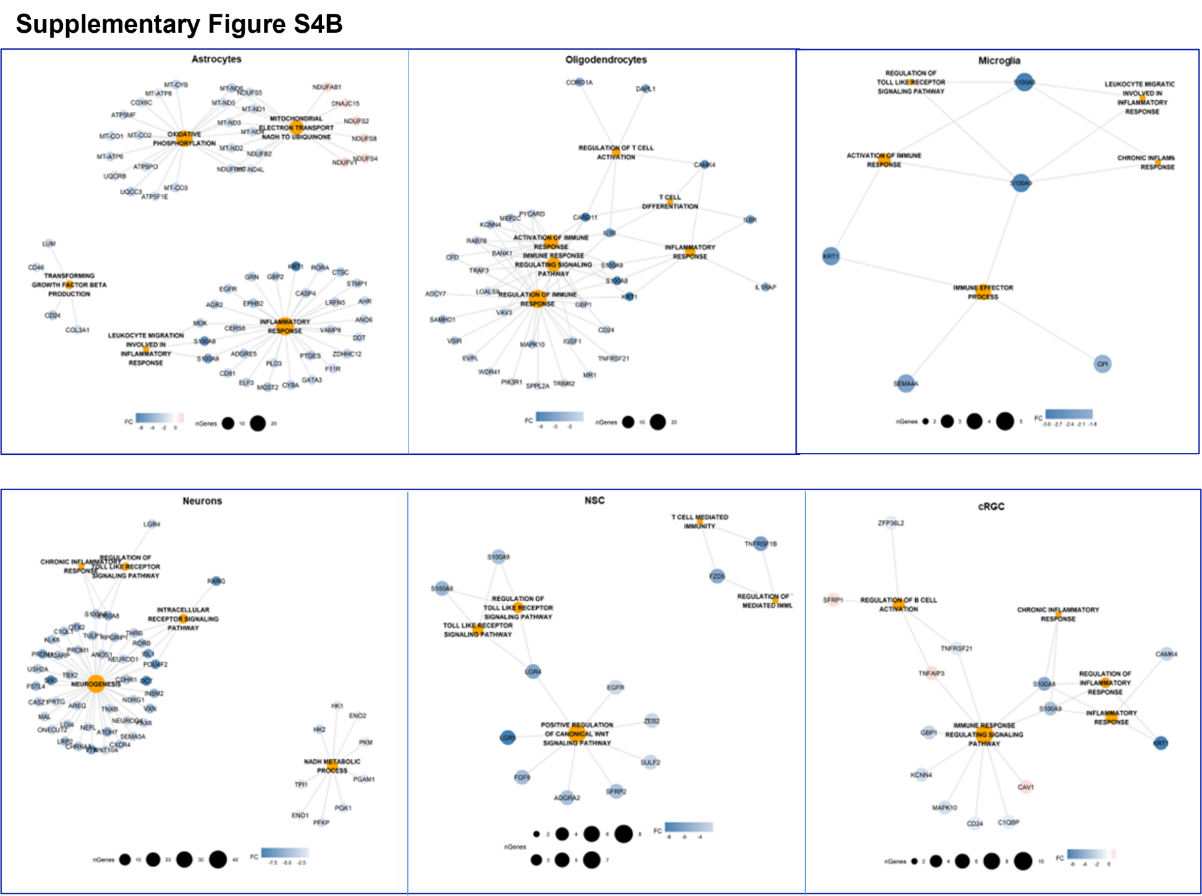

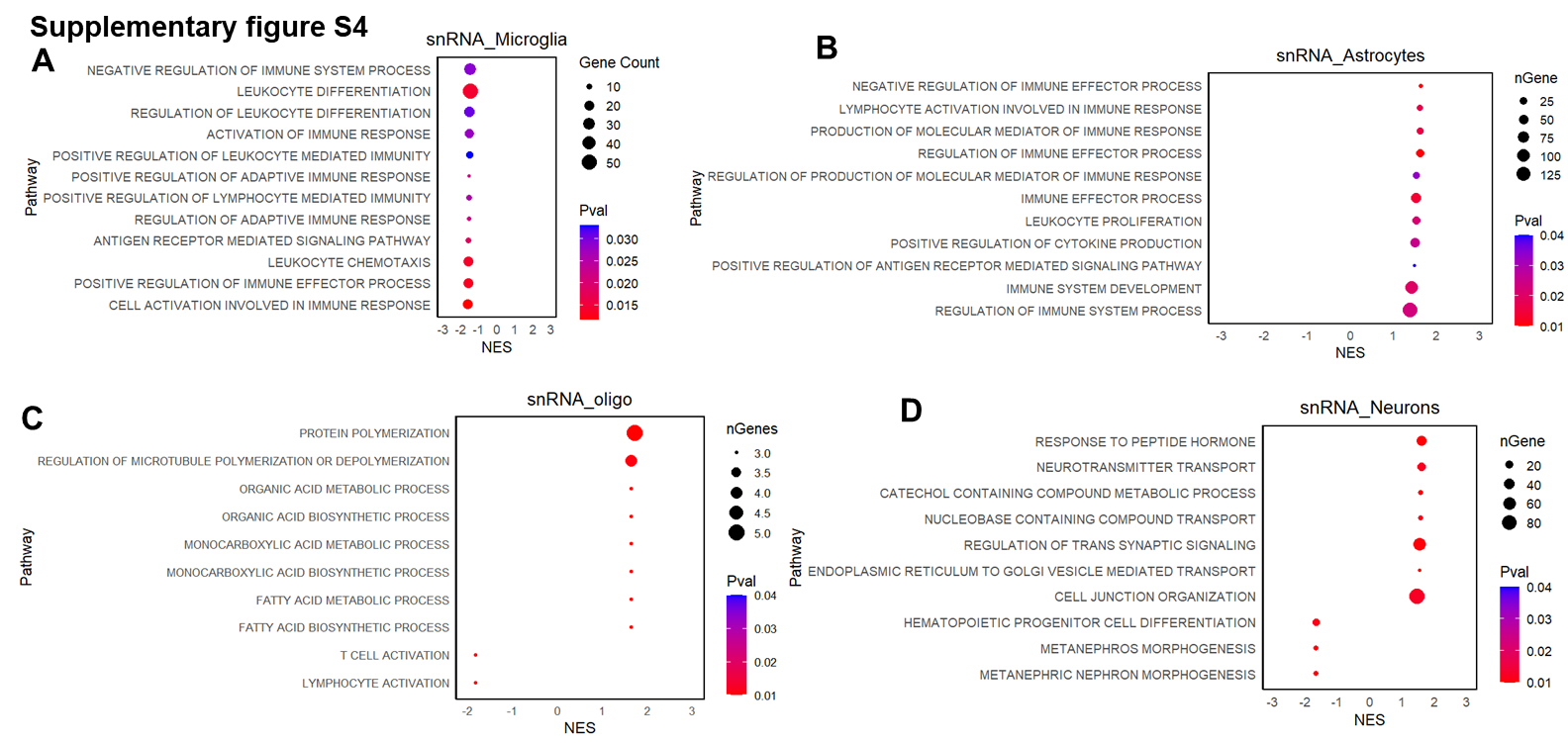

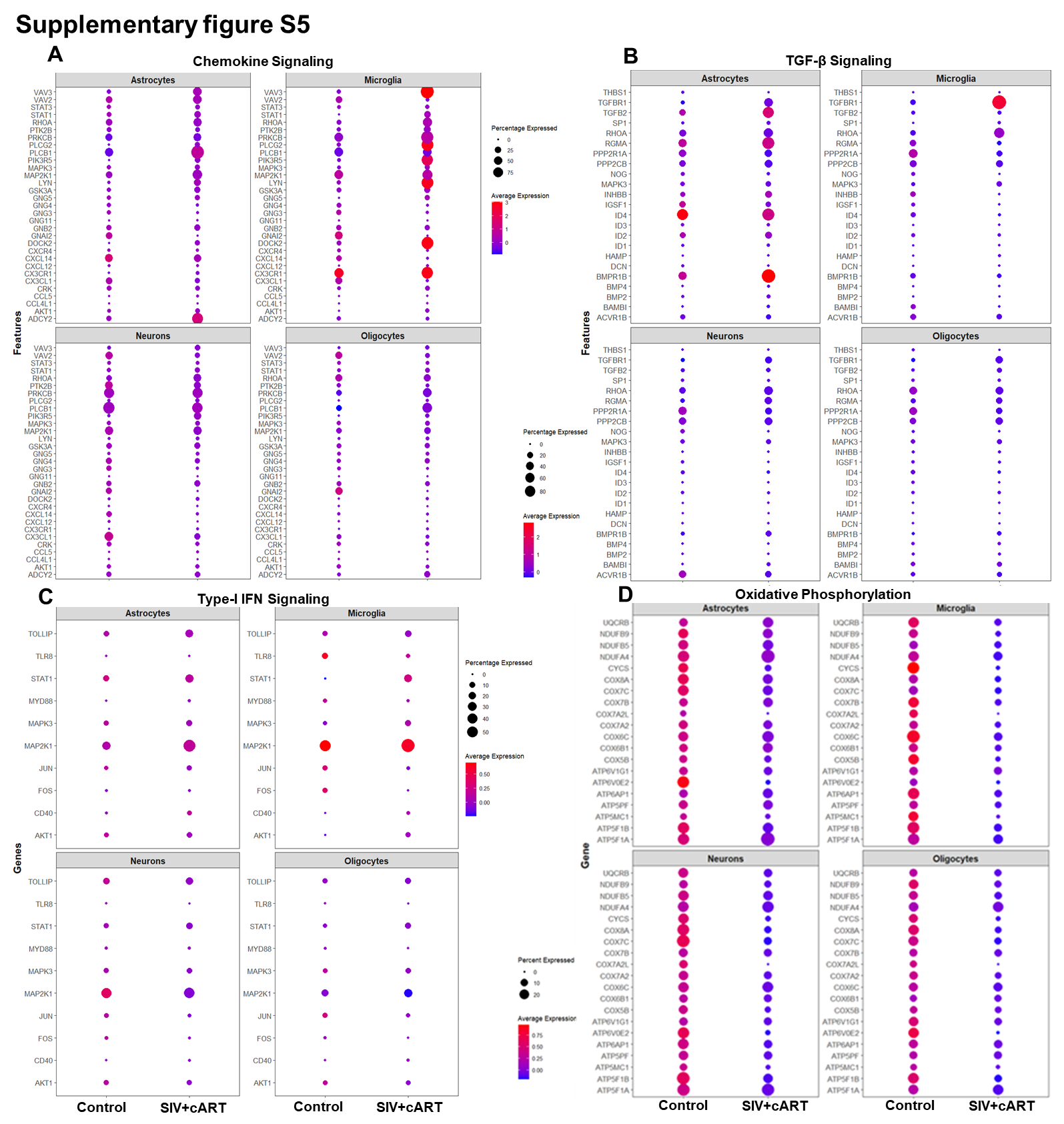

**Supplementary figure S6**

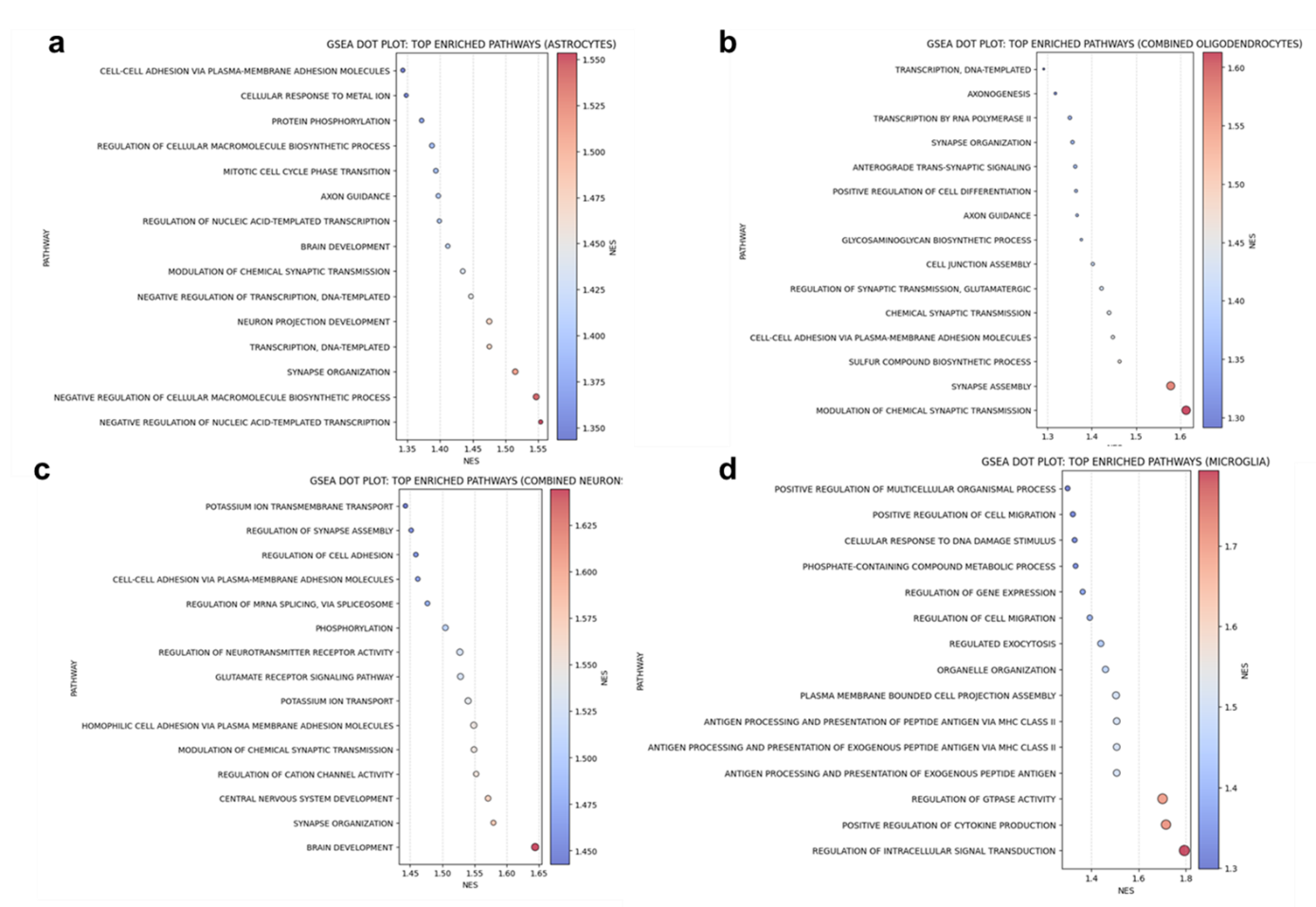

**Supplementary figure S7**

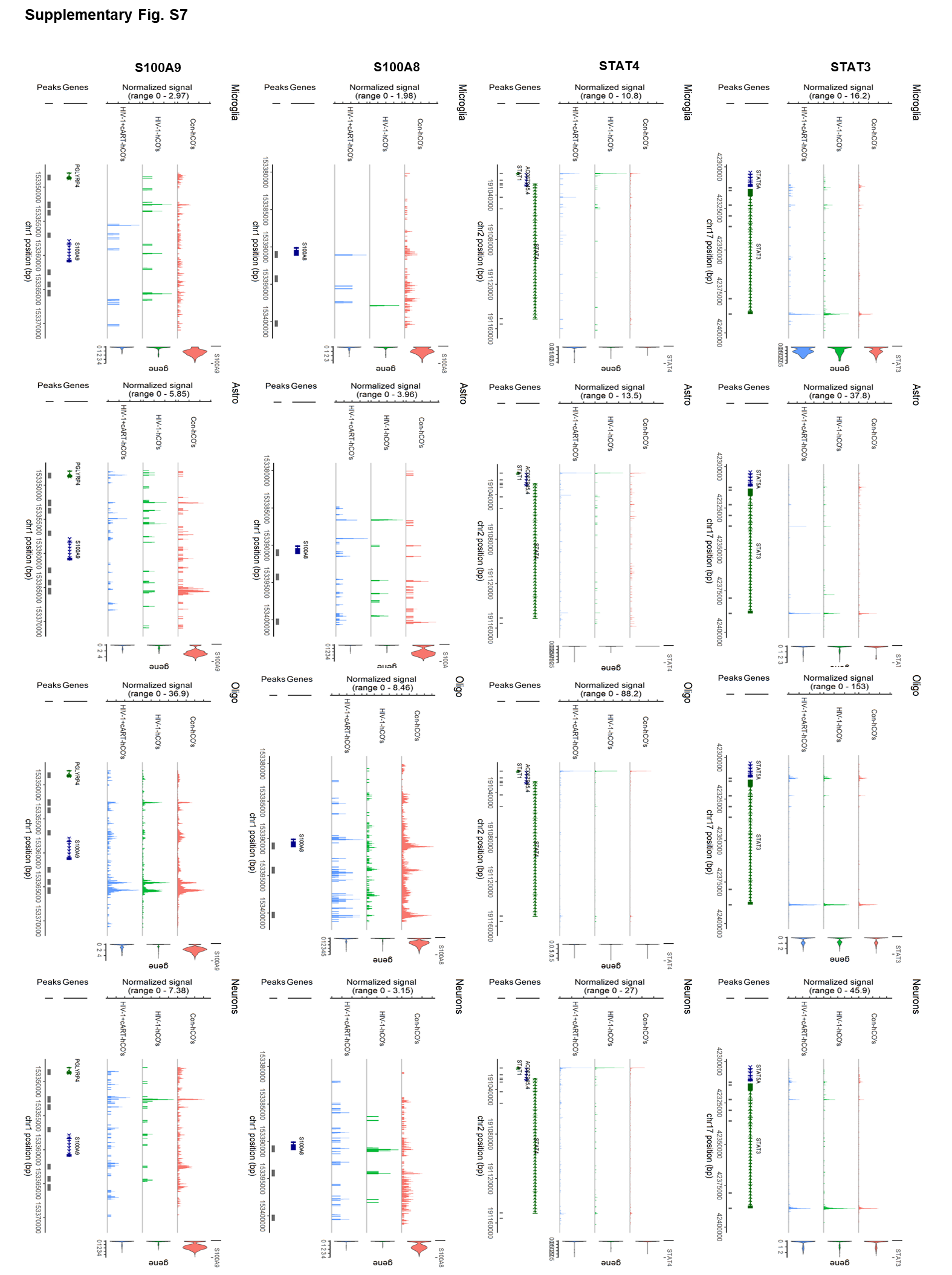

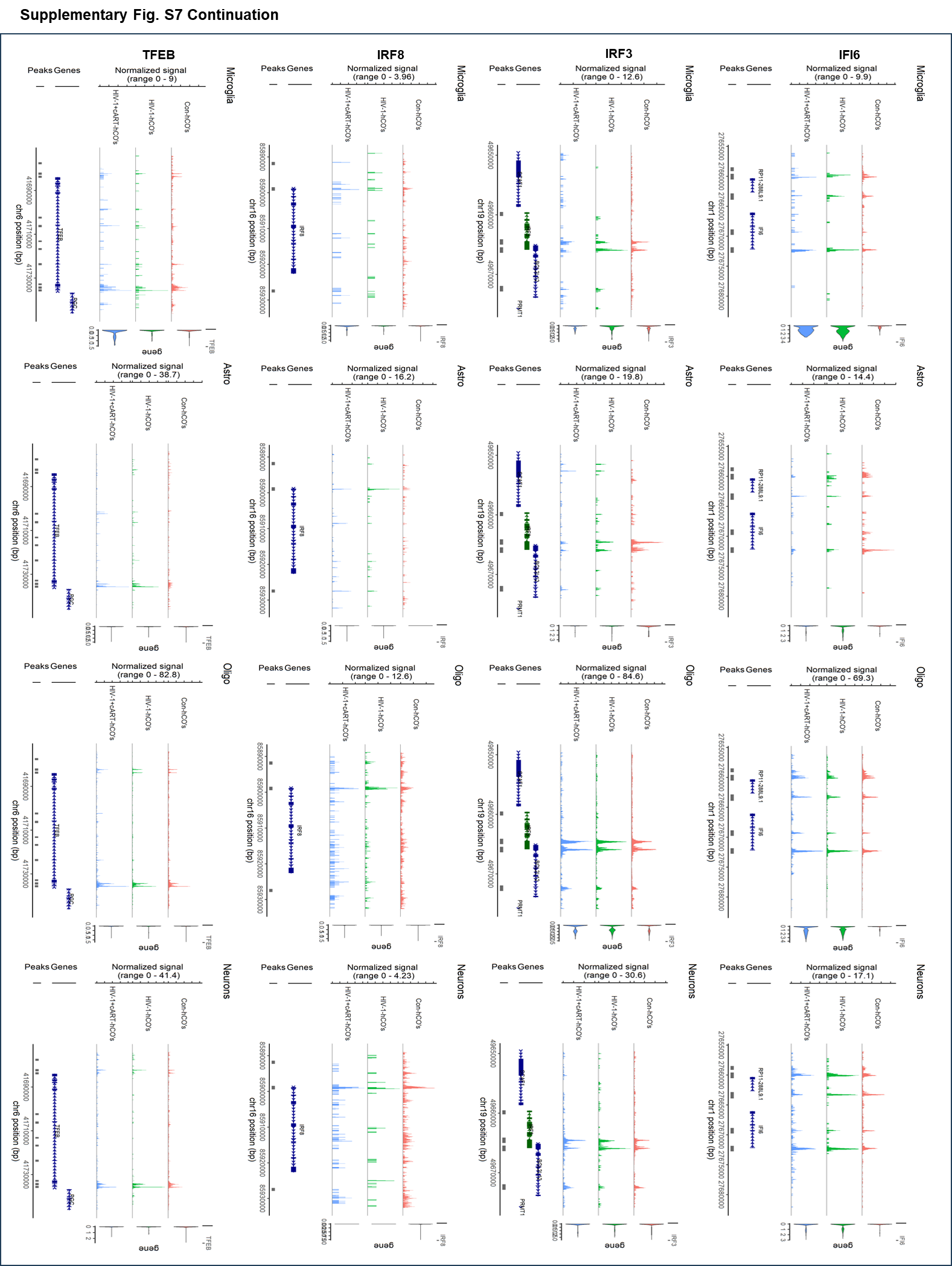

**Supplementary figure S8**

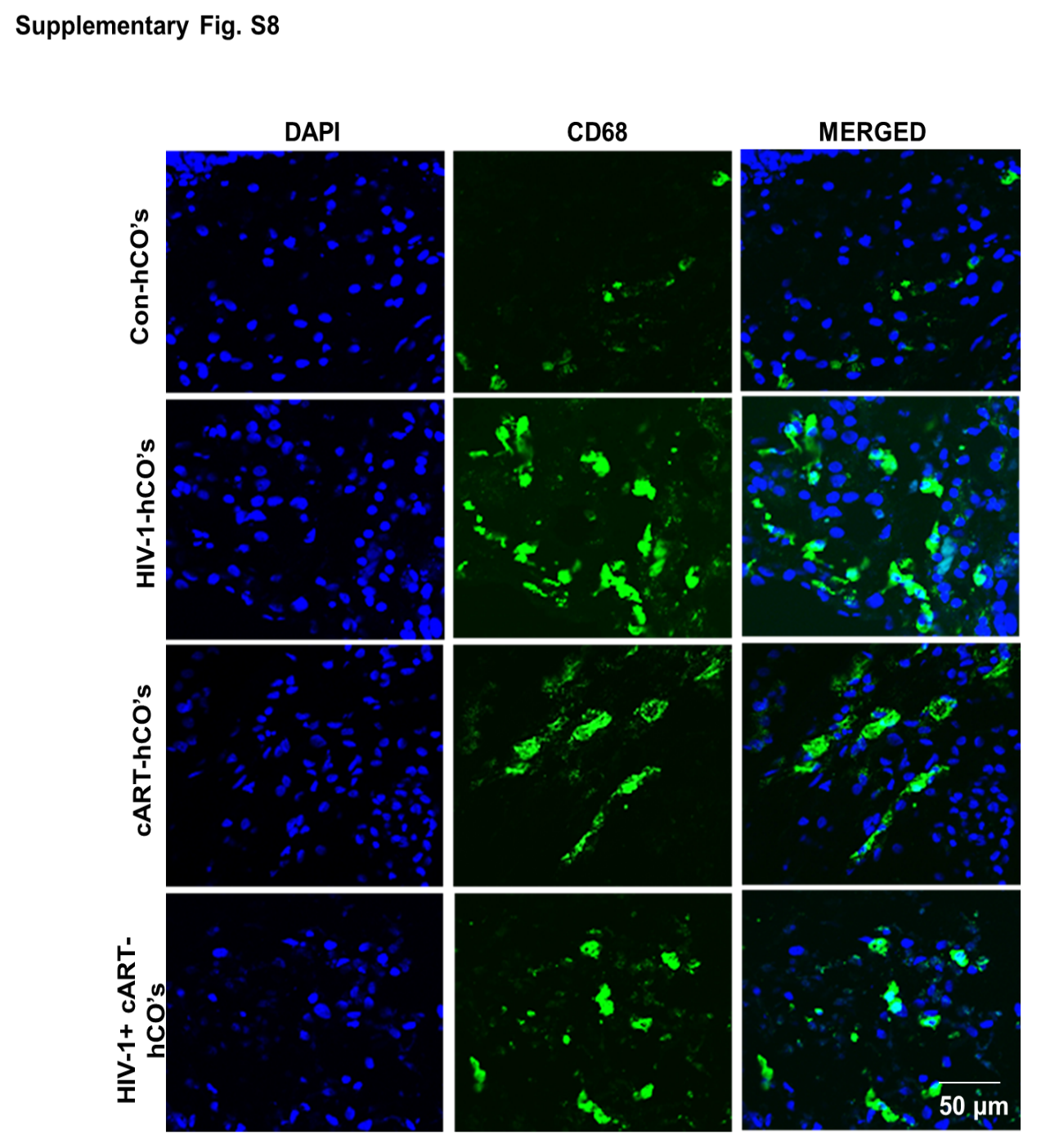

**Supplementary figure S9**

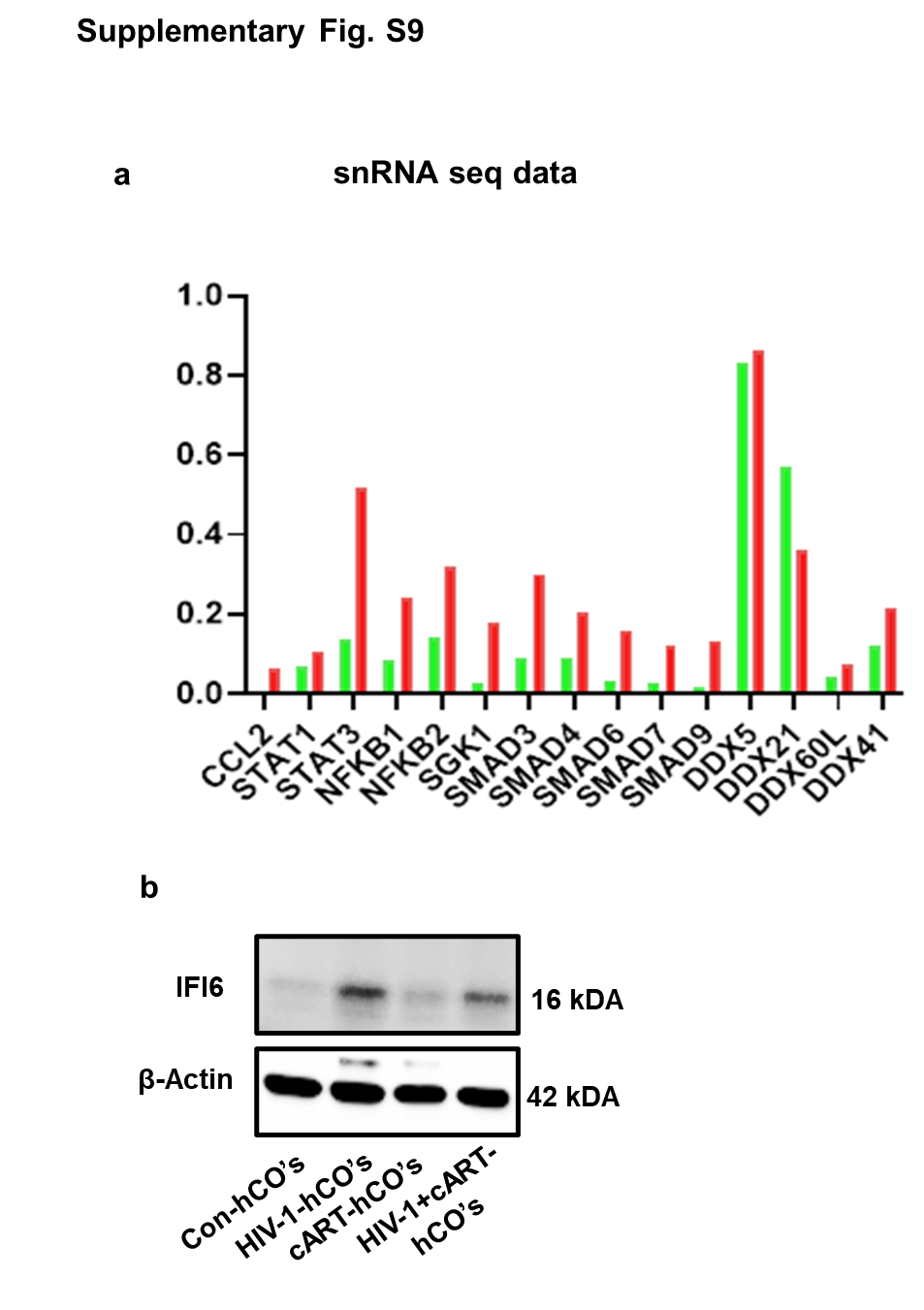

**Supplementary figure S10**

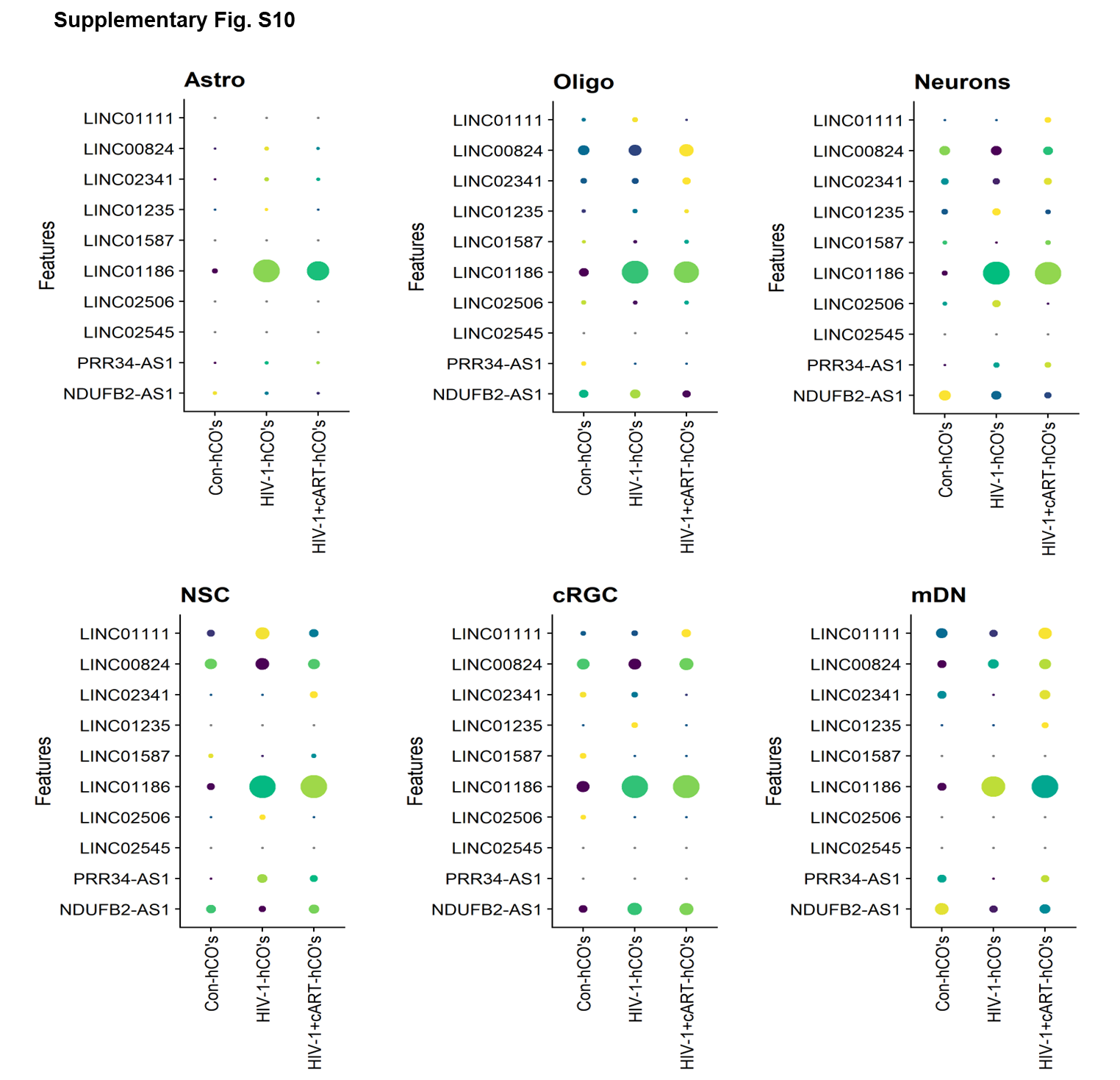

**Figure Legends**

**Supplementary figure S1**: Identifying key regulators of HIV-1-infected cerebral organoids through snRNA seq and ATAC seq analysis**. (A-F)** A dot plot visualizing differentially expressed gene levels of up and down-regulated inflammatory pathways, complement activation, TLR signaling pathway, and oxidative phosphorylation in different cell types of snRNA sequencing and **(G-I)** ATAC sequencing data.
