## Supplementary material for "STAT1-Mediated Regulation of IL-17A/CEBPB/NF-κB Axis in HIV-1 Infected Human Cerebral Organoids Reveals Therapeutic Targets for Neuroprotection": Excel file, Full blot, Tables: supplemental full blot images.docx

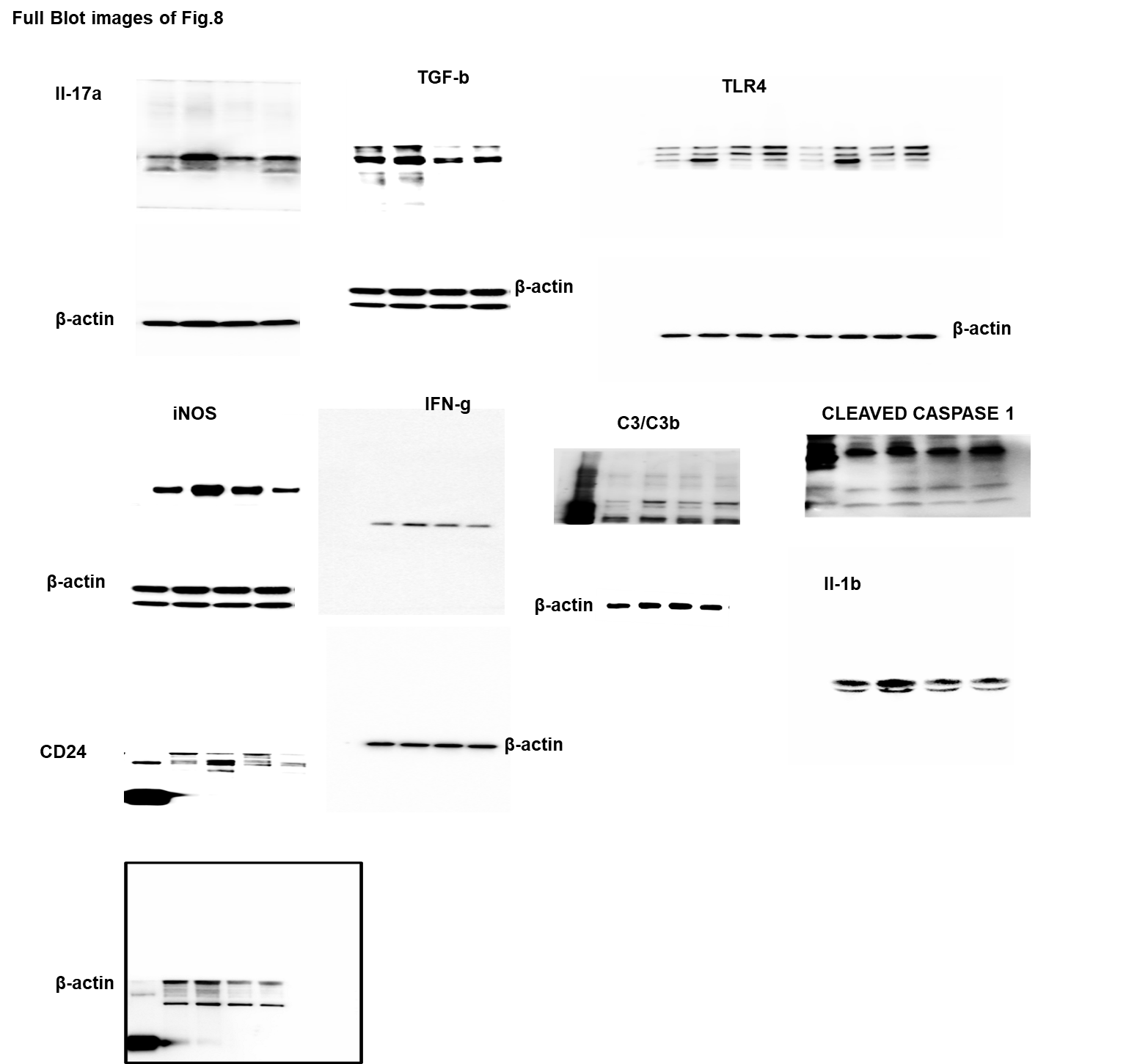
**Full blots Fig 7**

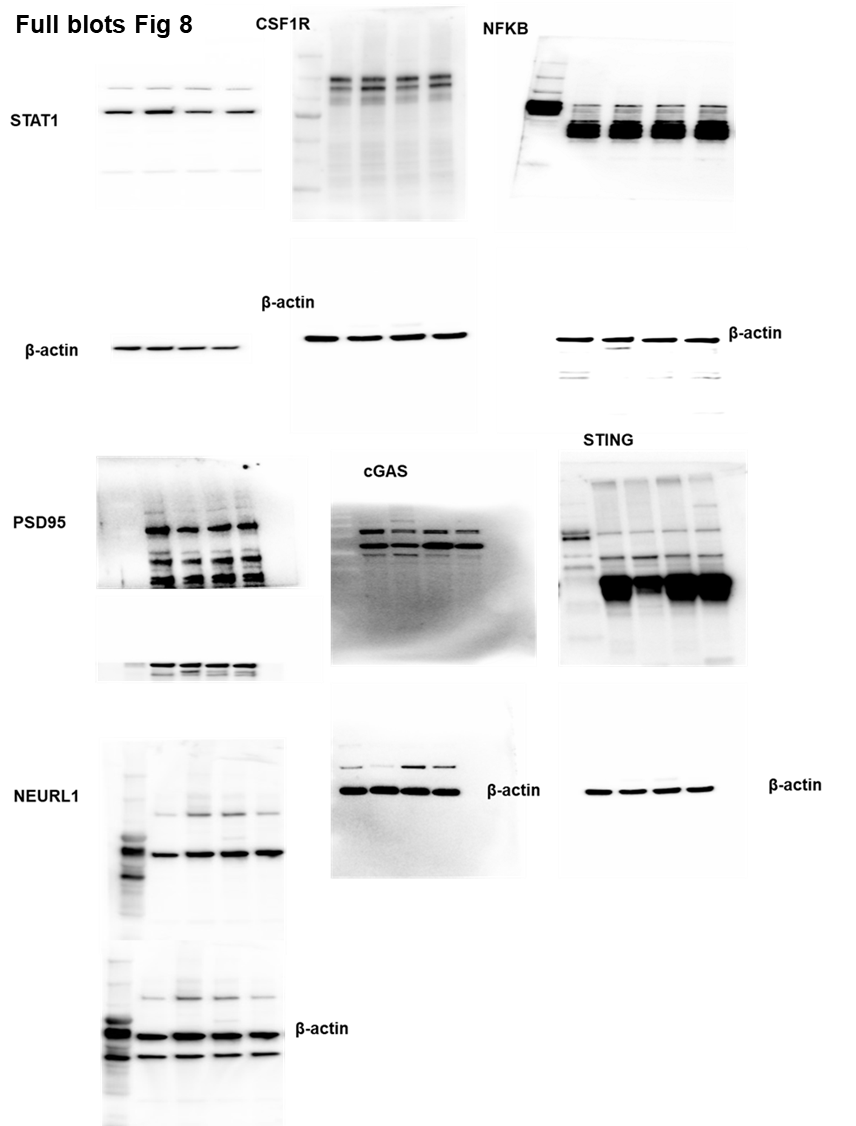

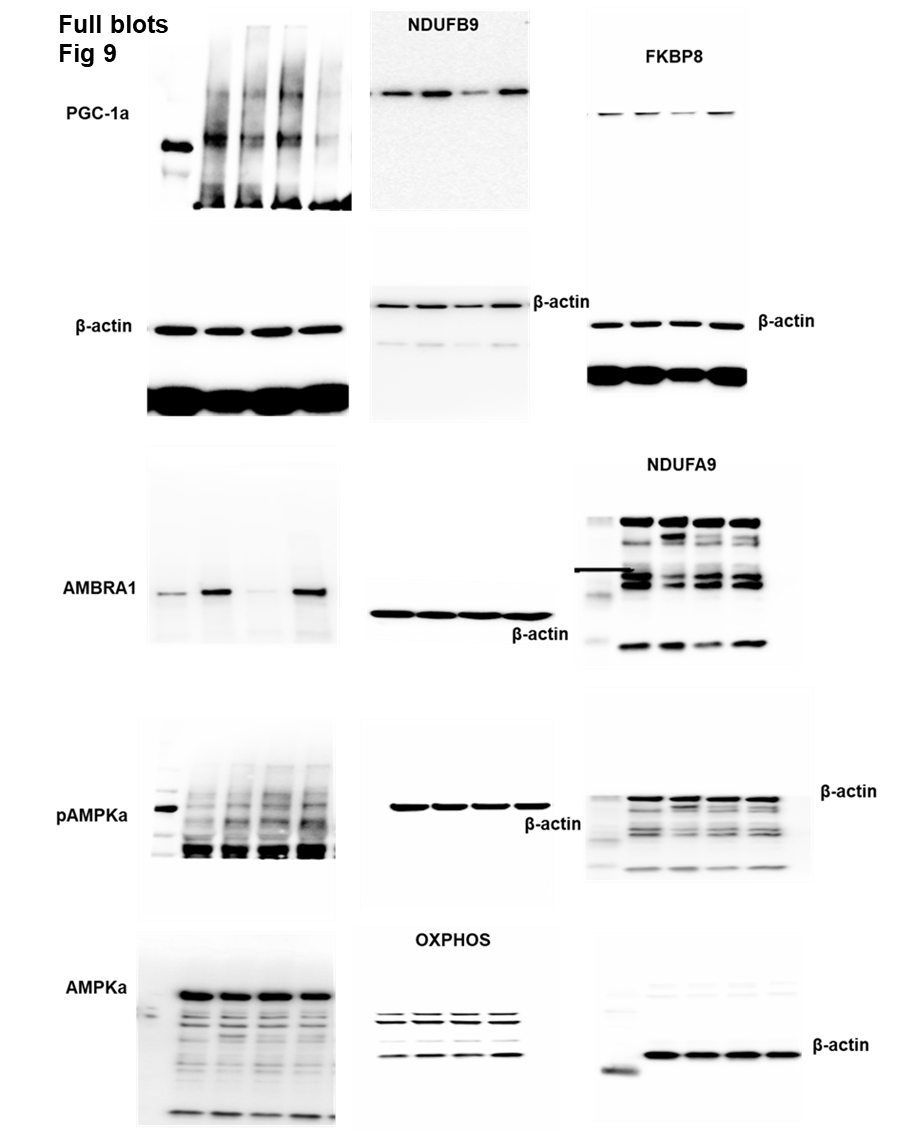

Full Blot images: To conserve samples and maintain experimental consistency, certain membranes were reprobed, and the same β-actin blot was reused as the loading control.
