## Supplementary material for "STAT1-Mediated Regulation of IL-17A/CEBPB/NF-κB Axis in HIV-1 Infected Human Cerebral Organoids Reveals Therapeutic Targets for Neuroprotection": Excel file, Full blot, Tables: Supplemental table.docx

**Table S1: List of Antibodies used for Western blot analysis**

| **Antibody** | **Company** | **Cat #** |
| --- | --- | --- |
| TLR 4 | Protein Tech | 66350-1-Ig |
| C3/C3b/C3c | Protein Tech | 21337-1-AP |
| CD 24 | Protein Tech | 67627-1-Ig |
| IL-17A | Protein Tech | 66148-1-Ig |
| IFN - gamma | Protein Tech | 15365-I-AP |
| TGF-β (56E4) | Cell Signaling | 3709T |
| IL-1 β | Protein Tech | 16806-1-AP |
| Cleaved caspase-1 (Asp297) (D57A2) | Cell Signaling | 4199T |
| Cleaved-IL-1β (Asp116) (D3A3Z) | Cell Signaling | 83186T |
| iNOS | Thermo Fisher Scientific | MA5-17139 |
| cGAS | Santa Cruz Biotechnology | SC-515777 |
| TMEM173/ STING | Protein Tech | 19851-1-AP |
| Stat1 (D1K9Y) | Cell Signaling | 14994T |
| Phospho-Stat1 (Tyr701) (58D6) | Cell Signaling | 9167T |
| PGC1a | Protein Tech | 66369-I-IG |
| NDUFB9 (E7W6K) | Cell Signaling | 99235T |
| NDUFA9 | ABclonal | A3196 |
| FKBP8 | Protein Tech | 11173-1-AP |
| AMBRA1 | Protein Tech | 13762-1-AP |
| Phospho-AMPK alpha1/2 (Thr183/172) | Elabscience | AB-21121 |
| AMPK alpha | Protein Tech | 10929-2-AP |
| OXPHOS | Abcam | ab110411 |
| TAU | ABclonal | A23490 |
| Phospho-Tau (Ser202, Thr205) | Thermo Fisher Scientific | MN1020 |
| MAP2 | Thermo Fisher Scientific | PA1-10005 |
| NF-κB p65 | Protein Tech | 10745-1-AP |
| Anti-CSF1R antibody [EPR21885-161] | Abcam | ab221684 |
| PSD-95 | Thermo Fisher Scientific | 51-6900 |
| Anti-Iba1/AIF1 | Millipore Sigma | MABN92 |
| Beta Actin | Cell Signaling | 12262S |

**Table S2: List of Antibodies used for immunocytochemistry**

| **Antibody** | **Company** | **Cat #** |
| --- | --- | --- |
| CD68 | Cell Signaling | 76437T |
| Anti-Iba1/AIF1 | Millipore Sigma | MABN92 |
| FOXO3A | Protein Tech | 10849-1-AP |
| TAU | ABclonal | A23490 |
| Phospho-Tau (Ser202, Thr205) | Thermo Fisher Scientific | MN1020 |
| SOX2 | Thermo Fisher Scientific | A11501 |
| MAP2 | Thermo Fisher Scientific | PA1-10005 |
| PAX 6 | ABclonal | A7334 |
| Nestin | ABclonal | A11861 |
| BIN1 (E4A1P) | Cell Signaling | 51844S |
| Ki67 | ABclonal | A2094 |
| GFAP | ABclonal | A0237 |
| FOXG1 | ABclonal | A16851 |
| TNF alpha | Thermo Fisher Scientific | PA1-40281 |
| Anti-Beta-Tubulin III Antibody | Stemcell Technologies | 60052 |
| PSD-95 | Thermo Fisher Scientific | 51-6900 |
| HIV-1 Gag p24 | Fisher scientific | MAB73601SP |
